# Maturation of human cardiac organoids enables complex disease modelling and drug discovery

**DOI:** 10.1101/2024.09.05.611336

**Authors:** Mark Pocock, Janice D. Reid, Harley R. Robinson, Natalie Charitakis, James R. Krycer, Simon R. Foster, Rebecca L. Fitzsimmons, Mary Lor, Natasha Tuano, Sara Howden, Katerina Vlahos, Kevin I. Watt, Adam Piers, Holly K. Voges, Patrick R.J. Fortuna, James Rae, Robert G. Parton, Igor E. Konstantinov, Robert G. Weintraub, David Elliott, Mirana Ramialison, Enzo R. Porrello, Richard J. Mills, James E. Hudson

## Abstract

Cardiac maturation is an important developmental phase culminating in profound biological and functional changes to adapt to the high demand environment after birth^1,2^. Maturation of human pluripotent stem cell-derived human cardiac organoids (hCO) to more closely resemble human heart tissue is critical for understanding disease pathology. Herein, we profile human heart maturation *in vivo*^3^ to identify key signalling pathways that drive maturation in hCOs^4,5^. Transient activation of both the 5’ AMP-activated kinase (AMPK) and estrogen-related receptor (ERR) promoted hCO maturation by mimicking the increased functional demands of post-natal development. hCOs cultured under these directed maturation (DM) conditions (DM-hCOs) display robust transcriptional maturation including increased expression of mature sarcomeric and oxidative phosphorylation genes resulting in enhanced metabolic capacity. DM-hCOs have functionally mature properties such as sarcoplasmic reticulum-dependent calcium handling, accurate responses to drug treatments perturbing the excitation-coupling process and ability to detect ectopy CASQ2 and RYR2 mutants. Importantly, DM- hCOs permit modelling of complex human disease processes such as desmoplakin (*DSP*) cardiomyopathy, which is driven by multiple cell types. Subsequently, we deploy DM-hCOs to demonstrate that bromodomain extra-terminal inhibitor INCB054329 rescues the *DSP* phenotype. Together, this study demonstrates that recapitulating *in vivo* development promotes advanced maturation enabling disease modelling and the identification of a therapeutic strategy for *DSP-* cardiomyopathy.

## Introduction

Understanding the maturation process is pivotal for many fields including the application of human pluripotent stem cell technologies for disease modelling and drug discovery. Although recent studies have begun to identify factors *required* for maturation^2,6^, it has been difficult to pinpoint the upstream physiological stimuli that are *sufficient* to drive the maturation process.

hPSC-CMs are typically immature limiting some of their applications for disease modelling and drug screening. They therefore also provide an excellent model to understand the maturation process. Extended culture for up to 1 year does not result in equivalent maturation to that of time matched *in vivo* maturation^7^, indicating that either environmental conditions are inhibitory and/or additional stimuli are required. However, the challenge is that different stimuli sequentially drive distinct aspects of maturation^4^, meaning that combinations of factors and precise timing of factors are essential. To this end, the five most successful drivers of maturation thus far are multicellularity, mechanical loading/structural patterning, hormonal stimulation, metabolic switching, and pacing with electrical stimulation^6^. Multicellular and organoid protocols have been established to promote cellular complexity and cell-cell interactions^5,8–11^. Tissue engineering approaches generally incorporate mechanical loading which drastically improves the morphology, alignment and maturity of the cardiomyocytes^4,12–14^. Metabolic maturation can also provide maturation of key features including metabolism, cell cycle and sarcomeric isoforms^4^. In addition, pacing is one of the most potent inducers of functional maturation in terms of excitation-contraction coupling and drug responses^15–17^. Despite these advances, mechanistic understanding of the maturation stimuli are limited, and the induction of further maturation is challenging. This may be important for modelling complex genetic or environmental driven diseases.

Heart-Dyno® is a 96-well platform that facilitates the self-organisation of cardiac cell types into miniaturised, mechanically loaded hCOs^4,5^. To create hCOs, hPSCs are carefully patterned into pre- cardiac mesoderm^18^, which gives rise to cardiomyocytes and cardiac progenitor cells in a single differentiation protocol. Subsequently hCO fabrication facilitates the differentiation and organisation of multiple cardiac cell types to form hCOs^5^. We have also identified metabolic conditions to promote maturation of hCOs^4^, yet, like all current protocols, adult properties remain elusive. Therefore, our approach provides a platform to screen for maturation conditions.

In this study we use our knowledge of human heart maturation ^3^ to screen for conditions that promote maturation of hCOs *in vitro*. We found that transient activation of AMPK and ERR provided the most robust induction of maturation. Directed maturation of hCOs (DM-hCOs) manifested in changes across the transcriptome and in contractile function, with an increased metabolic capacity. DM-hCOs enabled complex disease modelling including difficult to model genetic variants such as *DSP* cardiomyopathy, which could be functionally rescued with the bromodomain and extra-terminal protein inhibitor INCB054329.

## Results

We recently developed a serum-free protocol for hCO maturation (SF-hCO), featuring switching of metabolic substrates and vascularisation promoting growth factors^5^. We compared our SF-hCOs to various other hPSC-CM 3D cultures using mRNA expression ratios of sarcomeric proteins that are indicative of maturation^4,10,11,15,17^. *MYH7* as a fraction of *MYH7* and *MYH6,* did not correlate with maturation in any system, and is perhaps rate dependent (Extended Data 1a). *MYL2* as a fraction of *MYL2* and *MYL7* (Extended Data 1b) and *TNNI3* as a fraction of *TNNI3* and *TNNI1* (Extended Data 1c) were stronger indicators, with *TNNI3* in particular correlating strongly with maturation and being elevated by pacing protocols that are proven methods of maturation^15,17^. It is noted that our SF-hCOs displayed advanced levels of maturation as a strong baseline to screen for further maturation conditions (Extended Data 1c).

Our previous transcriptional profiling of human heart maturation identified a key oxidative metabolism and interferon signature that strongly underpins cardiomyocyte maturation^3^. In the current study, we screened for conditions that promote these features and determined their impact on cardiac maturation. Cardiac maturation is characterised by many different parameters, with different stimuli influencing distinct maturational properties^2,6^. We focused on key functional changes that occur with maturation – increased force, decreased automaticity and unaltered or reduced time to 50% relaxation (Tr50) and mature sarcomeric markers – cTnI (*TNNI3*)^7^.

### AMPK and ERR Agonists Drive Maturation

We firstly determined critical factors regulating metabolism or interferon signalling for a factorial maturation screen, see Extended Results and Extended Data 2-8 for identification of factors and concentrations. We identified 10 μM progesterone, 3 μM DY131 (ERRβ/γ agonist), 10 μM MK8722 (AMPK activator) and 100 ng/ml IFN-λ1 (Fig. 1a). Conditions with MK8722 consistently reduced rate without impacting Tr50 (Fig. 1b). MK8722 combined with DY131 or IFN- λ1 increased cTnI (Fig. 1c). When MK8722 and DY131 were further combined with either progesterone or IFN-λ1, there was no additional increase in cTnI (Fig. 1c). The timing of DY131 and MK8722 addition was also assessed (Extended Data 9). If added during the maturation medium period (days 17-22) or immediately after this period at the start of the weaning medium period (days 22-27), rate was still reduced but cTnI increase was limited. We next assessed the impact of the most abundant fatty acids in human breast milk and/or blood^19,20^ on maturation (Extended Data 10). Fatty acids were used during maturation (100 μM) and weaning media (10 μM) phases including palmitic (our standard protocol), linoleic, oleic, myristic acids and an equimolar combination of all four. We found that all fatty acids performed equally well, except for a combination of all four which caused functional decline. We therefore continued to use palmitate.

**Figure 1:**
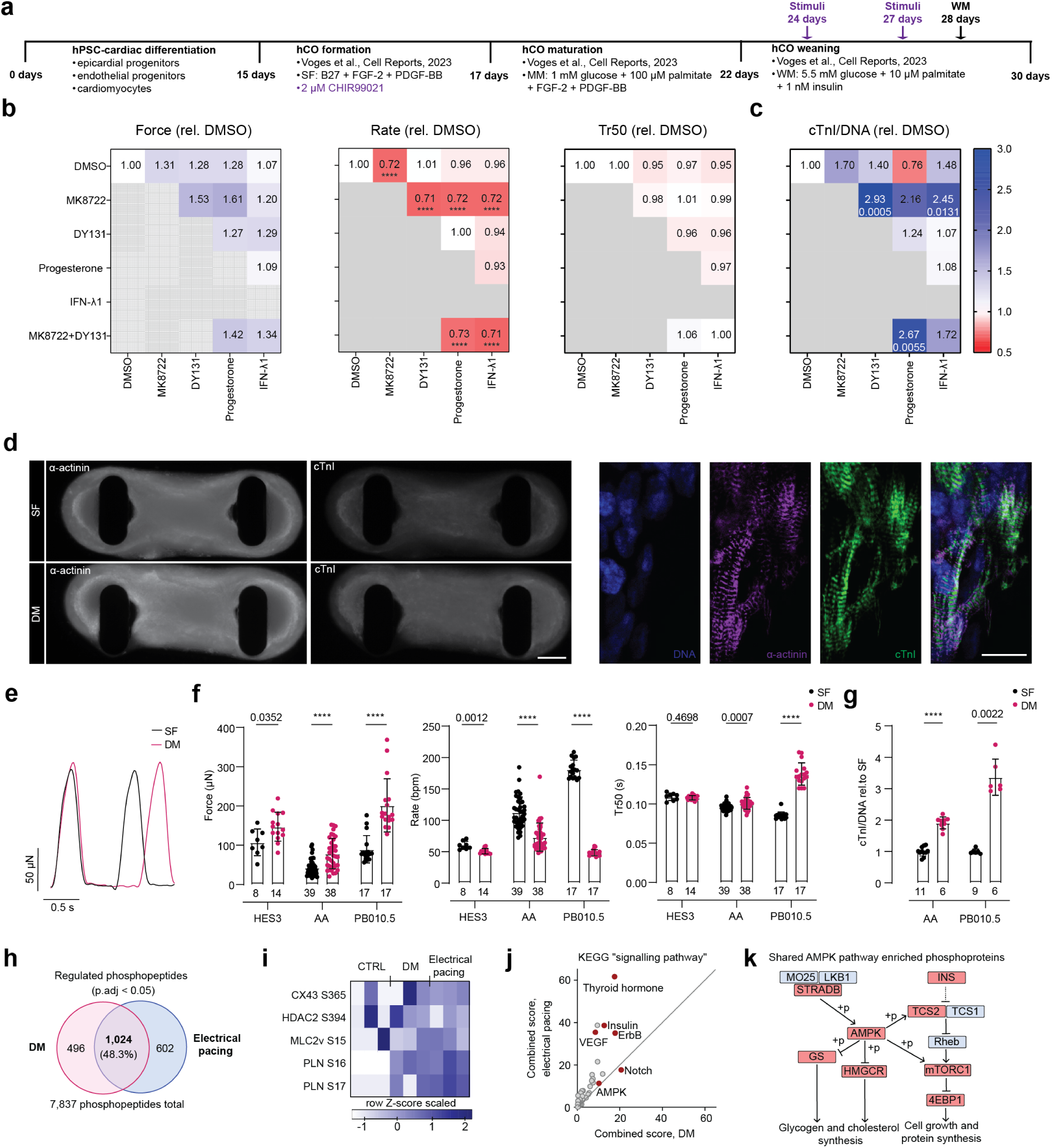
Screening for combinatorial stimuli to promote hCO maturation. **a**, Schematic of the protocol. **b**, Force of contraction, rate and time from peak to 50% relaxation (Tr50) at 30 days normalized to pre-treatment at 24 days. n = 3 experiments. **c**, Cardiac troponin I (cTnI) intensity normalized to DNA and then to DMSO controls. **d**, Cardiac troponin (cTnI) staining of cardiomyocytes (α-actinin). Bar = 200 μm**. e**, Comparison of SF- and DM-hCO raw contraction curves. **f**, Comparison of raw force of contraction, rate and Tr50 between SF- and DM-hCOs across 3 cell lines. **g**, cTnI intensity normalized to DNA and then to DMSO controls. **h**, Regulated phosphosites in MK8722 + DY131 (DM) treated and 120 bpm electrically paced hCOs (5 min stimulation each), highlighting shared sites. n = 3 biological replicates with 15 pooled hCOs each. **i**, Heatmap of phosphosite Z-scores. **j**, Gene ontology analysis of phosphorylated proteins shared between acute DM treatment of hCOs and electrical pacing. **k**, The AMPK signalling network shared between acute DM treatment of hCOs and electrical pacing. Concentrations were: 10 μM progesterone, 3 μM DY131, 10 μM MY8722 and 100 ng/ml IFN-λ1. Krusksal-Wallis test with Dunn’s multiple comparison to DMSO (**b,c**). For heatmaps (**b,c**) the p- values are given beneath the values if significant. Mann-Whitney tests (**f**,**g**).**** P < 0.0001.

We defined our new maturation conditions, designated directed maturation (DM-hCO). In summary, the key additions to our SF-hCO conditions^5^ are the addition of 2 μM CHIR99021 during the first 2 days of hCO formation and transient 4 day addition of 3 μM DY131 combined with 10 μM MK8722 (days 24-28). The ability of these conditions to reduce rate and increase cTnI was confirmed in multiple additional hPSC cell lines (Fig. 1d-g).To determine whether these conditions are broadly applicable to the field, we assessed the impact of our protocol in a 2D culture. The addition of MK8722 during a 4 day period (days 24-28) was also sufficient to reduce rate and increase cTnI in 2D hPSC-CMs (Extended Data 11).

As there is a close relationship between exercise, AMPK and metabolism in the heart^21^, we assessed the similarity in the activation profile of acute pacing and DM treatment using phosphoproteomics (Extended Data 12). We utilized a custom platform, where we added Heart-Dyno inserts into a C-pace system in 24-well plates enabling 120 bpm pacing for 5 minutes without causing toxicity. Phosphoproteomics revealed that there was substantial overlap of 1,024 (48.2% of regulated) phosphosites of DM treatment and electrical pacing (Fig. 1h). Phosphosites known to play a role in cardiac biology and function were detected including HDAC2 S394^22^, GJA1 (CX43) S365^23^, PLN S16/T17 and MLC2v S15^24^. PLN S16/17 consistently increased in both acutely DM treated and paced hCO and has been shown to drive increased SR calcium cycling^25^ (Fig. 1i). Similar KEGG pathways were activated in both DM and electrically paced conditions (Fig. 1j), including an AMPK network (Fig. 1k). This results in inhibition of the mevalonate/cholesterol biosynthesis program, which we have previously shown to be repressed during cardiac maturation^26^.

### hCOs Display Cellular Complexity Consistent with Human Hearts

Using snRNA-seq and integrated clustering with our previous human cardiac maturation dataset^3^, we found that multiple cardiac cell types including cardiomyocytes, endothelial cells, smooth muscle cells, fibroblasts and epicardial cells all co-cluster (Fig. 2a). Thus our single optimised differentiation SF-hCO protocol^5^ yields a complex mixture of cells similar to native human heart tissue, but with a higher percentage of cardiomyocytes, lower percentage of endothelial cells, lack of immune cell populations (Fig. 2a) and lack of a neural population at higher clustering resolution (data not shown).

**Figure 2:**
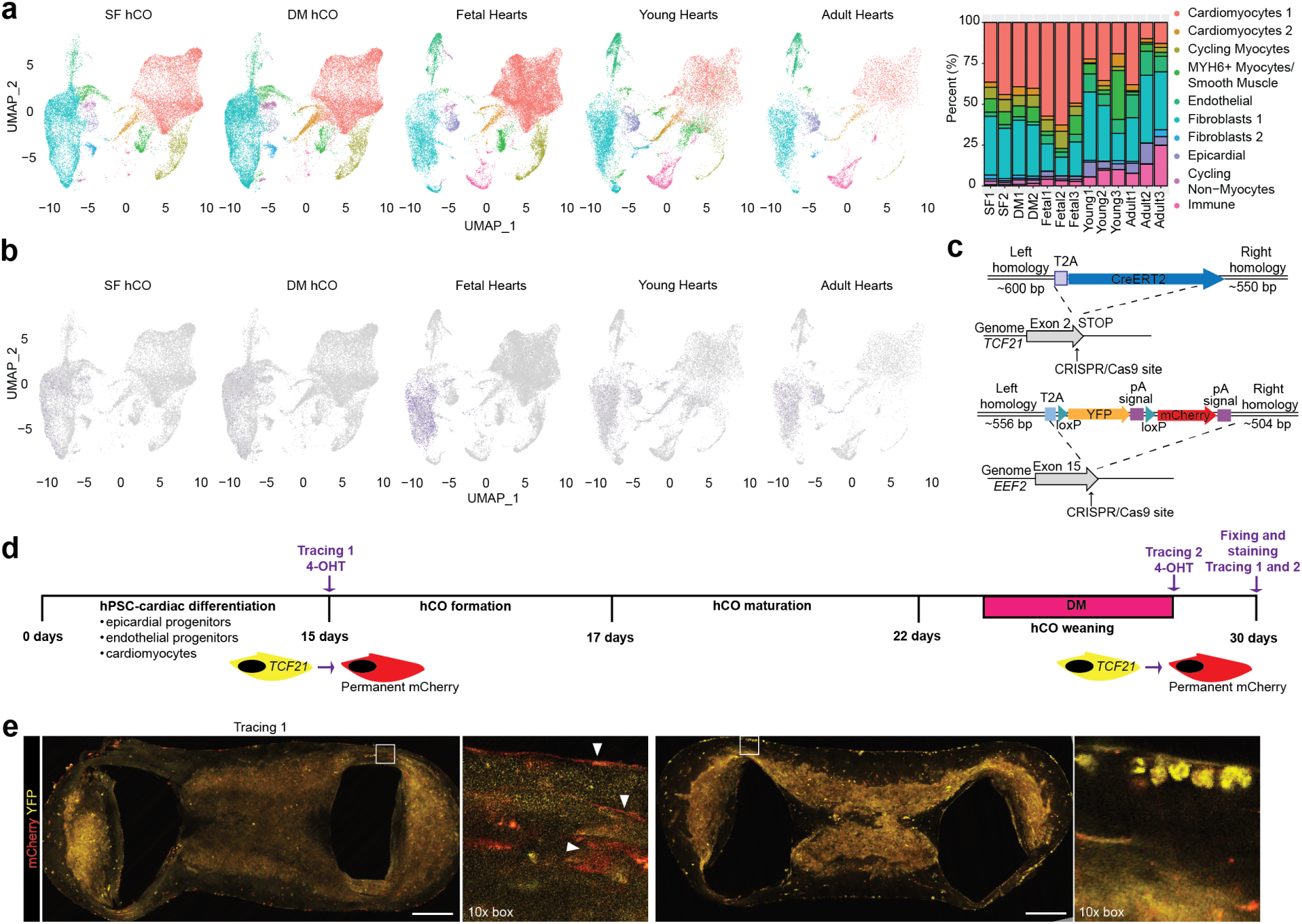
Cellular composition in SF- and DM-hCOs. **a,** Co-clustering of snRNA-seq from SF- and DM- hCOs and human heart data from GSE156707^3^, including proportions of cell populations. n = 2 snRNA-seq replicates for SF- and DM-hCO each from ∼70 pooled hCOs per sample. **b**, Expression of pro- epicardial organ marker *TCF21*. c, Generation of a hPSC *TCF21* lineage tracing line. **d**, Schematic of lineage tracing experiments. **e**, Representative analysis (n = 3 experiments) of lineage Tracing 1 and 2. Bars = 200 μm.

Further exploring fibroblast identity, we found some expression of the epicardial gene *TCF21* in the fibroblastic population indicative of epicardial origins (Fig. 2b). As TCF21 is reduced during differentiation and maturation^27^, we created a *TCF21* lineage tracing tool (Fig. 2c). We found that early, but not late lineage tracing marked the epicardial cells surrounding the SF- and DM-hCOs along with some infiltrating cells (Fig. 2d,e). Thus SF- and DM-hCOs contain TCF21 derived epicardium and stromal cells consistent with the human heart.

### hCOs Display Complex Cell-Cell Interactions

The DM-hCO protocol does not alter cellular composition, and immunostaining confirmed the presence of the major cardiac cell types in the hCOs including cardiomyocytes, epicardial cells, endothelial cells, pericytes and fibroblasts (Fig 2a, Extended Data 13). When clustering hCOs alone, the nuclei also segregate into distinct cellular populations (Fig. 3a, Extended Data 14), which are demarcated by canonical marker genes (Fig. 3b). The SF- and DM-hCO MYH6+ cardiomyocytes express critical *If* channel genes *HCN1* and *HCN4* and a critical pacemaker transcription factor *SHOX2*^28^ (Extended Data 15a,b). This population likely represents the pacemaker population within the SF- and DM-hCOs, which we have previously confirmed using patch-clamping on isolated cells^4^.

**Figure 3:**
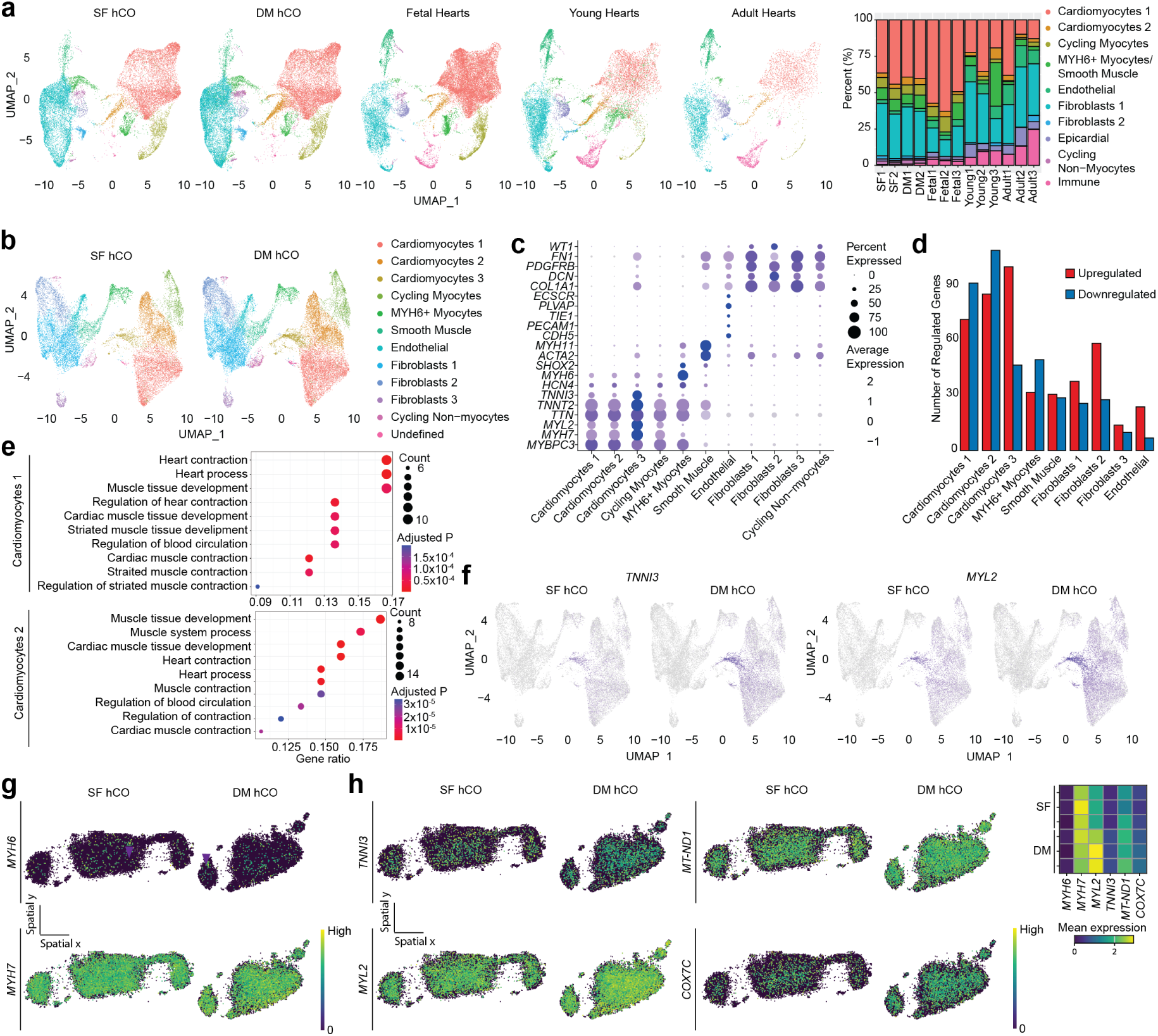
Sarcomeric and metabolic maturation in DM-hCOs. **a**, UMAP projection of nuclei in SF- and DM-hCOs. Nuclei labelled by cell type. **b**, Expression of canonical cell markers in SF- and DM-hCOs. **c**, Number of regulated genes in different cellular populations from comparison DM- versus SF-hCOs using pseudo-bulk analysis (average log2FC > |0.25|, adjusted P < 0.05). **d**, Top 10 gene ontology terms from the sub ontology ‘biological processes’ for upregulated genes in Cardiomyocytes 1 and 2 populations. **e**, Expression of sarcomeric maturation genes *TNNI3* and *MYL2*. **f**, Top 10 gene ontology terms from the sub ontology ‘biological processes’ for upregulated genes in Cardiomyocytes 3. **g**, Oxidation rate in SF- and DM-hCOs over the culture duration. n = 2 cell lines, HES3 and PB005.1. **h**, Response to BAM15 in SF- and DM-hCOs. **i**, Representative spatial expression of cardiomyocyte (*MYH7*) and nodal cardiomyocyte (*MYH6*) markers in a section. Purple arrows point to MYH6 clusters. **j**, Representative spatial expression of sarcomeric and metabolic genes in a section, including a heatmap of average expression. Two-way ANOVA with Tukey’s multiple comparison test. **** P < 0.0001.

During our SF- and DM-hCO protocol we only add exogenous growth factors FGF2 and PDGF-BB during the first week of hCO culture to form robust multicellular populations^5^. This suggests that maintenance of the fibroblasts and endothelial cells occur via endogenous paracrine factor production. Consistent with paracrine support, cardiomyocytes express *FGF10* the cognate ligand for the fibroblast receptor *FGFR2* (Extended Data 16a). While *PDGFRB* was abundantly expressed by the fibroblasts, the ligand *PDGFB* was expressed at very low levels (Extended Data 16b). This is potentially why longer term exogenous addition of PDGF-BB can lead to fibroblast overgrowth and irregular contraction patterns^5^.

The endothelial cells in both SF- and DM-hCOs have strong expression of key endocardial markers including *NRG1*, *NFATC1* and *GATA4*, but do not express coronary markers *APLN* and *FABP4* (Extended Data 17), consistent with our recent publication^5^. The endothelial cells and some fibroblasts express *FLT1*, and cardiomyocytes expressing its ligand *VEGFA* (Extended Data 16c), which is potentially why exogenous VEGF-A is not required in our protocol to support the endothelial population^5^.

### DM-hCOs Display Advanced Sarcomeric and Metabolic Maturation

When comparing the most populous cardiomyocyte cluster (Cluster 1) in SF-hCOs and adult hearts are already similar, and there are only 185 and 98 transcripts two fold lower or higher, respectively. Therefore, cardiomyocyte populations tightly clustered with human heart isolated cardiomyocytes (Extended Data 18), so we instead compared transcriptional changes. The DM-hCO protocol increases 20 and decreases 37 of these transcripts >log2|0.2| towards adult heart expression, respectively. This indicates a maturation response, including a repression of immature cardiac genes highlighted by increased TNNI3 as a fraction of TNNI3 and TNNI1 of 0.43, surpassing all other models examined (Extended Data 1).

DM- versus SF-hCO transcriptional changes were predominantly in the cardiomyocyte and fibroblast populations (Fig 3c). Genes upregulated in DM- compared to SF-hCOs in Cardiomyocytes 1 and 2 clusters were enriched in gene ontologies related to heart contraction and development (Fig. 3d). This includes key sarcomeric maturation markers *TNNI3* and *MYL2* (Fig. 3e) and key transcriptional regulators that control heart development *FOXP1*^29^, *SOX6*^30^, and *CSRP3*^31^. Genes upregulated in DM- compared to SF-hCOs in Cardiomyocytes 3 were enriched for gene ontology terms mostly associated with oxidative phosphorylation (Fig 3f). This includes direct ERRα/γ target genes^32^ particularly those involved with mitochondrial electron transport (*COX5A*, *COX5B*, *COX6A1*, *COX7C*, *COX8A* and *CYCS*) and ATP synthesis (*ATP5F1D*, *ATP5F1B*, *ATP5PO*, *ATP5MC1*, *ATP5PD*, *ATP5MC3* and *ATP5PF*). Real-time monitoring of hCO respiration revealed similar changes in respiration in SF- and DM-hCO protocols, which peaks during the maturation medium phase (Fig. 3g). At the end of the protocol spare respiratory capacity was higher in DM- versus SF-hCOs as assessed by treatment with the mitochondrial un-coupler BAM15 to reveal the maximal respiratory capacity (Fig. 3h). Together, DM- hCOs showed enhanced metabolic capacity consistent with cardiac maturation *in vivo*.

To assess whether these changes occurred homogenously across the tissue spatial transcriptome analysis using the STOMICS platform identified a total of 953 genes, which were mostly cardiomyocyte transcripts. The cardiomyocytes were spread throughout the SF- and DM-hCOs (*MYH7*), whereas the *MYH6*+ pacemaker cardiomyocytes were present sporadically throughout with some clustered regions which may help facilitate initiation of endogenous contractile activity (Fig. 3i). DM-hCOs had uniform induction of the sarcomeric maturation markers *MYL2* and *TNNI3* and metabolic genes *MT-ND1* and *COX7C* (Fig. 3j). This highlights that maturation is homogenous in DM-hCOs, indicating that there are limited diffusion barriers in this miniature system and that the Cardiomyocytes 3 cluster is not confined to a particular region.

While the DM-hCO cardiomyocytes had the most regulated transcripts, there are also some regulated genes in the stromal cells. Some of these are also fibroblast sub-type specific such as *COL15A1*, *COL19A1* and *TGFBR3* which may play a key role in extracellular matrix biology and paracrine interactions (Extended Data 19).

### Mechanisms of DM-hCO Decreased Automaticity are Multi-Factorial

Immature hPSC-CMs typically display robust automaticity largely driven by extracellular membrane currents and sarcoplasmic reticulum (SR) contributions^33,34^. Engineered heart tissues display increased maturity, and the automaticity has been shown to be regulated by both *If* and SR calcium cycling clocks^35^ (consistent with experiments in rabbit sinoatrial nodal cells^36^). In this series of experiments, we investigated whether hCO rate is controlled by similar processes to understand why DM-hCOs have reduced contraction rates. Both SF- and DM-hCOs have similar calcium sensitivities for force and no calcium-rate relationship, thus ruling out changes in calcium sensitivity (Fig. 4a). We blocked *If* using 1 μM cilobradine, because the clinically approved compound ivabradine at <10 μM only partially blocks *If*^37^ and prolonged Tr50 at higher concentrations (data not shown). Following blockade of *If* with 1 μM cilobradine, hCOs became dormant with ‘bursts’ of activity which are far less common in DM-hCOs compared to SF-hCOs (Fig. 4b). The decreased spontaneous activity of DM-hCOs under 1 μM cilobradine blockade is indicative of more functionally mature cardiomyocytes.

**Figure 4:**
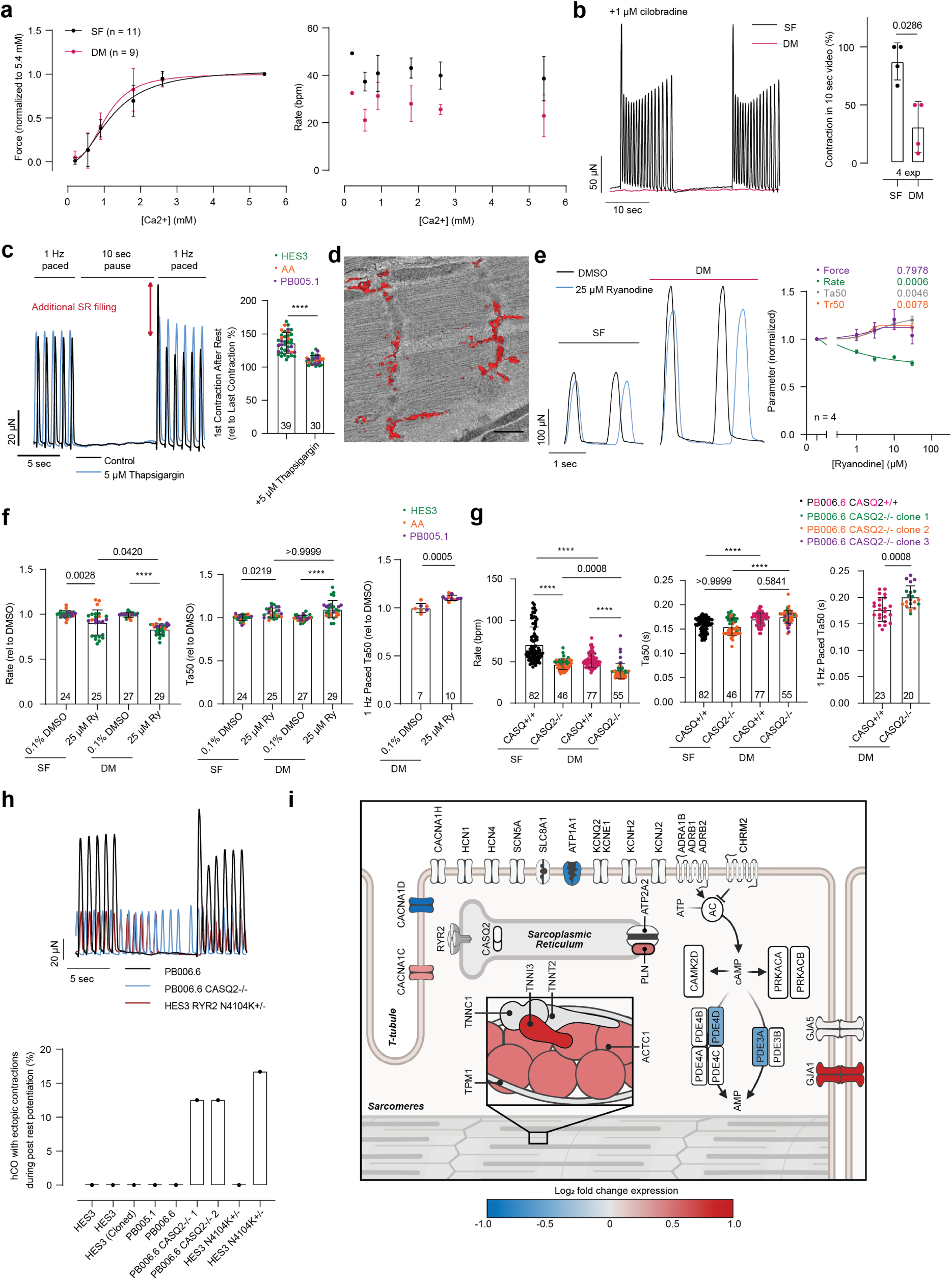
Mechanisms of rate control and SR handling in SF- and DM-hCOs. **a**, Dependence of force and rate on extracellular calcium concentration. **b**, Contractile rate/burst behaviour with blockade of If using 1 μM cilobradine. **c**, Post-rest-potentiation assessment of SR loading and verification using blockade of SERCA using 5 μM thapsigargin. **d**, 3D transmission electron microscopy rendering of the SR (red). Scale bar = 500 nm. **e**, Concentration-response curve of DM-hCOs treated with ryanodine and representative trace curves. Richard’s 5 parameter dose-response was used to fit the curves. **f**, Influence of ryanodine on rate and time from 50% activation to peak (Ta50), including under 1Hz pacing for DM-hCOs. **g**, Influence of CASQ knockout on rate and Ta50, including under 1Hz pacing for DM-hCOs. 3 independent experiments for PB006.6 CASQ+/+. **h**, Quantification of ectopic contractions during the pause phase of post-rest-potentiation experiments. **i**, Critical excitation-contraction genes regulated in DM versus SF-hCO cardiomyocyte populations. Created with BioRender.com. Mann-Whitney (**b**), Welch’s t test (**c,f,g**), Mixed-Effects testing with Dunnett’s post hoc analysis (**e**) and Kruskal-Wallis test with Dunn’s post-hoc analysis (**f,g**). **** P < 0.0001. Schematic produced in Biorender (**i**).

To determine the role of SR cycling in pacing, we first demonstrated that DM-hCOs have a functional SR. The ability to block contractile activity in DM-hCOs allowed us to perform post-rest-potentiation experiments, which enable measurement of functional SR in cardiac muscle^38^. After a 10 second pause the increase in force of the first re-paced contraction is indicative of increased SR filling. In DM-hCOs this was 36% (Fig. 3c), which is similar to human hearts (43%) at the same time interval^38^. This increase no longer occurs when sarcoendoplasmic reticulum calcium ATPase (SERCA) is inhibited using 5 μM thapsigargin, thus confirming a functional SR in DM-hCOs (Fig. 4c). Three-dimensional transmission electron microscopy of a random section in a DM-hCO also revealed the presence of an extensive SR network (Fig. 4d), which was confirmed in both SF- and DM-hCO sections (Extended Data 20).

We next determined the contribution of SR calcium release on rate control by blocking RyR2 with 25 μM ryanodine, a concentration that blocks the channel^39^. This decreased the rate with a larger magnitude in DM-hCOs (15%) compared to SF-hCOs (6%), indicating greater basal RyR2 leakiness in SF-hCOs (Fig. 4e). Force remained constant and Ta50 increased, which remained elevated when paced at 60 bpm to correct for frequency-dependent acceleration of relaxation (Fig 4f). To further confirm the role of SR handling in dictating hCO rate, we knocked out the SR calcium buffering protein CASQ2 using CRISPR gene editing (Extended Data 21a). In comparison to an isogenic control, rate declined in both SF- and DM-hCOs (Fig. 4g). There were no changes in force, and Ta50 was elevated in DM-hCOs when paced at 60 bpm (Fig 4g). These consistent forces combined with increased Ta50, are consistent with SR blockade in larger mammals (that are less reliant on SR calcium for contraction) and experiments performed with higher extracellular calcium concentrations^35,40,41^. Furthermore, the mechanism of increased Ta50 is potentially due to L-type calcium channel compensation^42,43^. Collectively, these data confirm that DM-hCOs have a functional SR that regulates endogenous pacing rate, contributes to contraction, and is potentially less leaky in DM- versus SF-hCOs.

In the DM-hCOs where spontaneous contractions are less frequent, we were able to model SR dysfunction driven ectopy. For this we used the CASQ2 knockout cell lines and also used CRISPR to generate a pathogenic variant cell line RYR2^+/N4104K^, which increases risk of arrhythmia in human patients^44^ (Extended Data 21b). Using our post-rest potentiation protocol, we observed ectopy in CASQ2^-/-^ and RYR2^+/N4104K^ DM-hCOs (Fig. 4h). This also provides further confirmation of a functional SR in DM-hCOs.

These changes between SF- and DM-hCOs are underpinned by multifactorial expression changes in excitation-contraction coupling genes. In DM- versus SF-hCOs *PLN* was increased (Fig. 4i), which may be in part responsible for the reduced SR impact on rate and reduction in SR leak. However, there were also additional changes that could alter contraction rate including decreased expression of *CACNA1D* (with a concomitant increase in *CACNA1C*), and increased *GJA1* (*CX43*) expression (Extended Data 14a,c,d and Fig. 4i). A reduction in *CACNA1D* activity has been shown to reduce rate in nodal cells^45^, and CX43 enhances cell-cell coupling, which also alters rate in hPSC-CM^46^. Collectively, all of these changes in DM- versus SF-hCOs may lead to the reduced rate of contraction and are consistent with the impact of AMPK (gamma 2 subunit) activation in decreasing heart rate in humans^47^.

### DM-hCOs Accurately Predicts Responses to Cardioactive Drugs

Accurate prediction of *in vivo* drug responses is a property required for widespread use of hCOs in drug discovery applications. DM-hCOs were benchmarked using 12 *Comprehensive In Vitro Proarrhythmia Assay* (CiPA) compounds that bind the human ether-or-go-go (hERG) channel with different risk stratifications for arrhythmia^48^ and a set of 17 compounds that are inert, negative or positive inotropes covering a diverse array of calcium transient and sarcomere regulators.

In the CiPA screen we found that time to 50% relaxation (Tr50) was the most predictive parameter for risk stratification (extended Data 22). This was not surprising as prolonged contraction duration is a clinical predictor of long QT syndrome, and is the contractile analogue to calcium transient duration (CaTD) and action potential duration (APD), which are commonly used in CiPA assays^48^ and contraction duration is also a key arrhythmia predictor clinically^49^. All high risk CiPA compounds that we tested increased Tr50 by 32-64% at the concentration closest to Cmax. Further increases in concentrations either stopped contraction (quinidine) or induced arrhythmias (ibutilide and dofetilide). Low risk compounds (ranozaline, mexiletine, diltiazem, and verapamil) did not increase Tr50, even at superphysiological concentrations. None of the intermediate risk compounds (cisapride, terfenadine, ondansetron, and chlorpromazine) increased Tr50 at Cmax. Concentration-dependent responses above Cmax were variable in this group. Cisapride and terfenadine lowered force, and odansetron substantially increased Tr50 (50%). Together this indicates that the hCOs are reliable in segregating low and high risk compounds, but assigning intermediate risk is more challenging..

In the boutique set of cardioactive compounds, the inert compounds paracetamol and pravastatin did not alter any of the contractile parameters (> 10%) at the highest concentrations (Extended Data 23a). Drugs inhibiting systemic cardiovascular factors including atenolol (β-adrenoreceptor inhibitor) and captopril (angiotensin converting enzyme inhibitor) did not directly alter any of the contractile parameters (> 10%) at the highest concentrations (Extended Data 23a), confirming that there is limited basal adrenergic drive or a renin-angiotensin system in the DM-hCOs. Likewise, clonidine (α_2_- adrenorecptor agonist) which primarily acts on neurons to alter heart function had no impact on DM- hCOs, consistent with the lack of neurons in our culture (Fig 2).

Negative inotropes sunitinib (RTK inhibitor), verapamil (ICa,L inhibitor), flecainide (INa inhibitor), mavacamten (myosin conformation) and aficamten (myosin conformation) all concentration-dependently decreased force (Extended Data 23b). Flecainide also initially increased Tr50 at low concentrations as expected. These drugs also impact other contractile parameters as forces substantially decline.

Inotropic compounds were screened at 0.6 mM Ca^2+^ given the higher calcium sensitivity in DM-hCOs (Fig 3a) compared to the human heart ∼2.6 mM^50^. BAYK-8644 (ICa,L activator) also increased force, but with a large liability on diastolic function with 250% Tr50 increase. Isoprenaline (β-adrenoreceptor agonist) concentration-dependently increased rate, then force at higher concentrations together with decreasing Ta50 and Tr50 (Extended Data 24a). Phenylephrine (α1-adrenorecptor agonist) increased force at the highest concentrations (Extended Data 24a). We also found that ouabain (INa/K inhibitor) increased force, prior to contraction cessation at higher concentrations (Extended Data 24a). Milrinone (PDE3 inhibitor) increased the force of contraction in the presence of 10 nM isoprenaline, with limited changes in other parameters (< 10%) (Extended Data 24b). We confirmed that PDE inhibition, particularly PDE4 by rolipram, potentiated the isoprenaline response increasing both the efficacy and potency of the isoprenaline response in DM-hCOs from multiple cell lines (Extended Data 24b).

For sarcomeric acting inotropes, CK-136 (nelutroctiv, troponin activator), which is under clinical assessment^51^, increases force prior to increasing Tr50 (102%) at the highest concentration tested (Extended Data 24c). Omecamtiv mecarbil (myosin activator) concentration dependently increases force until >1 μM where it starts to substantially increase Tr50 and force declines (Extended Data 24c) similar to other experimental paradigms^52^. This is consistent with data in human ventricular muscle preparations, and as an inotropic effect was not observed in human atrial preparations, this also further confirms the predominantly ventricular phenotype of our DM-hCOs^53^. More recently, danicamtiv (myosin activator) has been developed to overcome this liability and we find it substantially increases force with limited increases in Tr50 (20%) (Extended Data 24c). Specifically, when comparing omecamtiv mecarbil and dancamtiv at their upper serum concentrations in the clinic (1 μM^54^ and 8 μM^55^, respectively) we found that danicamtiv consistently increased force without the contraction duration liability of omecamtiv mecarbil in DM-hCOs from multiple cell lines (Fig. 5 a-c).

**Figure 5.**
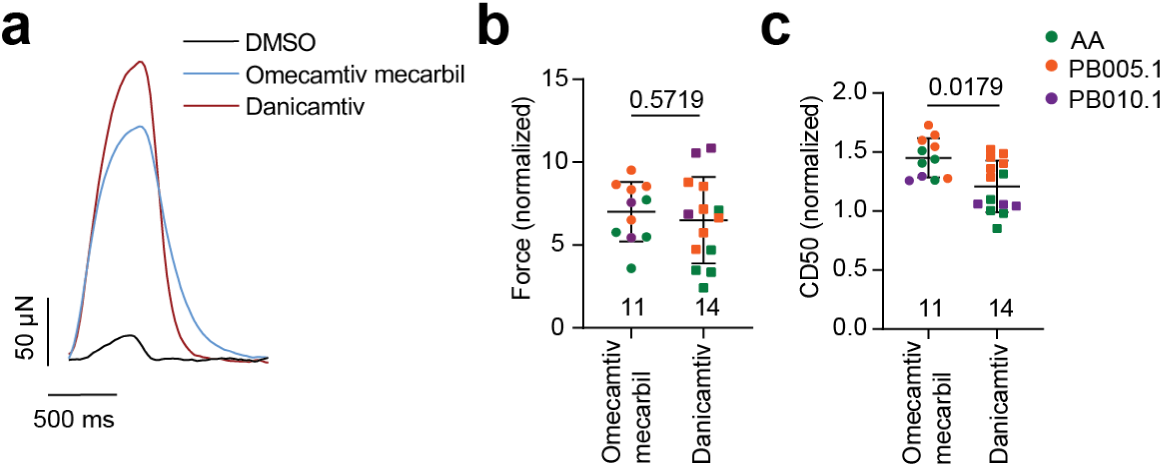
Myosin activators differentially affect contraction duration in DM-hCOs. **a-c**, Testing of omecamtiv mecarbil and dancamtiv at Cmax values, 1 μM and 8 μM, respectively. Experiments were performed at 0.6 mM Ca^2+^ by mixing weaning medium made in RPMI and DMEM base. **a**, Representative force curves. **b**, Force of contraction normalised to pre-drug baseline. **c**, Contraction duration between 50% activation and 50% relaxation. Mann-Whitney (**b**,**c**).

Taken together this suggests that DM-hCOs accurately predict many drug responses with similar IC50/EC50 to those reported in the literature (Extended Data Table 1). However, there are still attributes of this model consistent with an immature phenotype. In comparison to adult hearts, DM-hCOs have a higher calcium sensitivity, PDE4>PDE3 activity (although similar EC50 for milrinone as the human heart), and less sensitivity to α1-adrenoreceptor stimulation (Extended Data Table 1). These features are all consistent with the expression of the key genes involved in our RNA-sequencing data: *PDE3* (lower than human heart), *PDE4* (higher than human heart) and *ADRA1A* (lower than human heart). We also found that using our weaning medium culture conditions, positive inotropes can still be detected based on increases in force production (Extended Data 25), which is important for statistical power and comparisons in screening applications as otherwise small changes in calcium EC50 can have large impacts on the magnitude of force increases.

### DM-hCOs Enable Modelling of Complex DSP Cardiomyopathy

DSP cardiomyopathies are driven by complex cell-cell interactions and changes in excitation-contraction coupling^56^. As DM-hCOs faithfully recapitulate the multicellular composition of heart tissue and display features of mature excitation-contraction coupling, we chose to model a DSP mutation to demonstrate the utility of this platform for disease modelling. We identified a patient presenting with dilated cardiomyopathy and arrhythmia (MCHTB11), diagnosed with a homozygous 2 bp deletion in the *DSP* gene DSP c.4246_4247del; p.Leu1416AsnfsTer23. This results in leucine being replaced by asparagine at amino acid position 1416, followed by a termination codon after 22 amino acids in the new reading frame. The variant affects the longer isoform of DSP (NM_004415.3; 2871 amino acids), but not the shorter isoform (NM_001319034.2; 2428 amino acids). The family was screened, and the parents found to be heterozygous, and two deceased siblings homozygous for the mutation (Fig. 6a). We created an induced pluripotent cell line (DSPmut) and corrected line (DSPcorr) using simultaneous CRISPR correction with hiPSC reprogramming^57^ (Fig. 6b, Extended Data 26a). DM- hCOs displayed normal junctional expression of DSP and CX43 in the corrected H.3 line, but largely absent DSP expression and aberrant CX43 localisation together with disordered sarcomeres in the DSPmut line (Fig. 6c).

**Figure 6.**
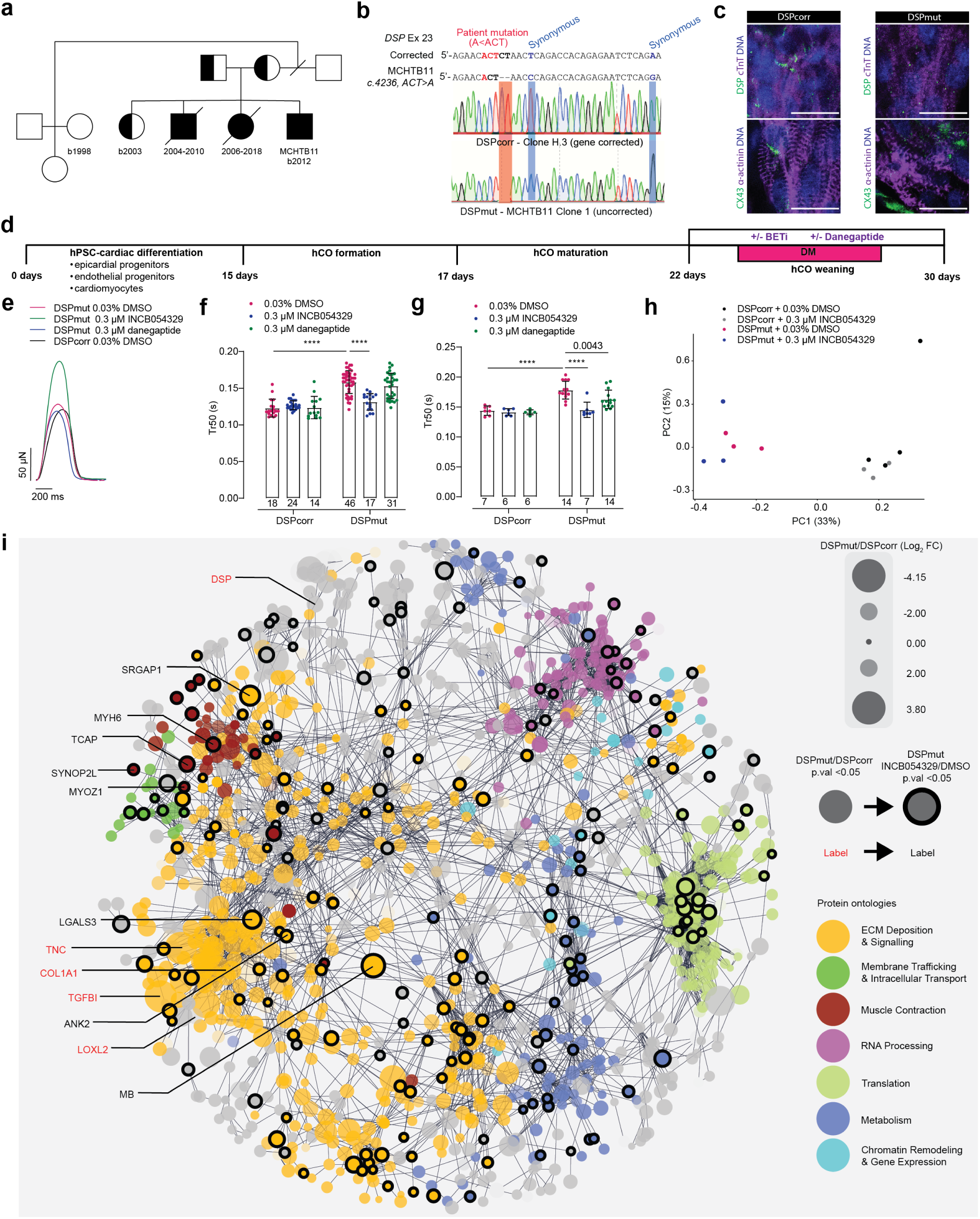
Modelling DSP cardiomyopathy and identification of therapeutic candidates in DM-hCOs. **a**, Family tree for the patient with dilated cardiomyopathy and arrhythmia, MCHTB11. **b**, Patient specific and CRISPR corrected human induced pluripotent stem cell lines. **c**, DM-hCO staining for DSP and CX43. bars = 20 μm. **d**, Schematic of functional assessment in DM-hCOs. **e**, Representative force traces. **f**, Time from peak to 50% relaxation (Tr50) with treatment of INCB054329 and danegaptide in DM-hCOs. n = 3-4 experiments. **g,** Tr50 under acute 60 bpm pacing with treatment of INCB054329 and danegaptide in DM hCO. n = 3-4 experiments. **h**, Principal component plot of proteomics data, n = 3-4 experiments with 3 DM-hCOs pooled in each. **i**, Interaction map of regulated proteins in DSPmut versus DSPcorr with highlighted circles also regulated by INCB054329 in DSPmut. Two-way ANOVA with Sidak’s post-hoc test between lines for DMSO or relative to DMSO for DSPmut (**f,g**). **** p < 0.0001.

We next sought to determine whether there were functional differences between DSPmut and DSPcorr SF- and DM-hCOs (Fig. 6d). There were large differences in DSPmut versus DSPcorr contraction rate in SF-hCOs, which may mask other key phenotypes (Extended Data 26b). In contrast, in DM-hCO the contraction rate in DSPmut and DSPcorr was relatively stable, which unmasked a phenotype whereby Tr50 was increased by 30% (Extended Data 26b).

In an attempt to improve the relaxation phenotype, we trialled two drugs: INCB054329 and danegaptide. INCB054329 is a bromodomain and extra-terminal protein inhibitor that we recently discovered can correct inflammation induced diastolic dysfunction^58^. Danegaptide is a gap junction enhancer^59^ that was also assessed due to the CX43 remodelling we observed (Fig. 6c), as well as recent interest in CX43 gene therapies for *DSP* cardiomyopathy. INCB054329 and danegaptide had no substantial effects in the DSPcorr DM-hCOs other than a slightly reduced rate for INCB054329 (Extended Data 26c). In DSPmut DM-hCOs INCB054329 increased force, slightly decreased rate and fully corrected the Tr50 phenotype (Fig. 6f,g, Extended Data 26c). In contrast, in DSPmut DM-hCOs, danegaptide slightly improved Tr50 which was only revealed under acute pacing conditions (Fig. 6g).

Given the improvements with INCB054329, we investigated the underlying mechanisms using proteomics (Extended Data 27a). There was distinct clustering of DSPmut versus H.3 DM-hCOs indicating the DSP mutation drives substantial proteomics changes (Fig. 6h). While there were no major changes in cellular population markers (Extended Data 27b), desmosomal proteins including DSP and DSG2 were decreased in DSPmut DM-hCOs (Extended Data 27c) and there was a strong upregulation of immunomodulatory proteins including CD47, THBS1, IL-10, TGF-β1 (Extended Data 27d). The changes in DSPmut versus DSPcorr DM-hCOs were dominated by upregulation of extracellular matrix proteins, which was strongly underpinned by TGF-β1 activation (Fig. 6i). However, there were protein changes across multiple additional processes including membrane proteins,, muscle contraction, RNA processing, translation, metabolism and chromatin regulation indicating a strong remodelling response (Fig. 6i). INCB054329 reverted a subset of the DSPmut regulated proteins in DM-hCOs (Extended Data 27e-g) without rescuing the expression of the desmosome proteins or repressing THBS1, IL-10 or TGF-β1 (Extended Data 27c,d). However, INCB054329 reverted multiple proteins in each of the highlighted dysregulated ontologies in the DSPmut (Fig. 6i), indicating broad remodelling activity toward DSPcorr. Collectively this indicates that there is a strong immunomodulatory response caused by DSP mutation which drives fibrosis and alters contraction kinetics, and INCB054329 improves function by reverting broad protein dysregulation rather than distinct control of the overall fibrotic response^60^.

## Discussion

In this study we use our Heart-Dyno hCO platform to investigate whether key *in vivo* signatures we identified in human hearts play a role in *in vitro* cardiac maturation^3^. We assessed key drivers of increased metabolic capacity, activation of AMPK, activation or ERRβ/γ and progesterone^3,21,32^. Together MK8722 and DY131 were the most effective drivers of increased maturation. In contrast to previous studies^61^, activation of ERR (using DY131) had a relatively subtle impact whereas activation of AMPK using MK8722 dramatically enhanced maturation. Critically, these factors were most effective following - but not during - the phase where hCOs are metabolically switched to fatty acid oxidation^4^. This may be because during this phase the cardiomyocytes are already near maximum capacity as indicated by our data profiling the oxidation over the full hCO culture time-course. Therefore, further metabolic stress under conditions of low glucose provision and high fatty acid provision is detrimental. Together, these data further support our previous findings that oxidative metabolism is not just a feature, but a driver of cardiac maturation^4,62^.

The acute application of DM induced a similar signature to pacing with over 48 % of activated phosphopeptides shared. Therefore, the critical AMPK signalling network we identify may play a key role in maturation under both pacing and DM-hCO protocols. We found that high contraction rates alone do not promote maturation, as PB010.5 SF-hCOs had endogenous beating rates >150 bpm but low levels of cTnI expression, which were then greatly enhanced in DM-hCOs. Therefore, increases in metabolic capacity are essential for maturation. Even after the DM stimuli are removed the increase in maturation indicators such as cTnI and reduced endogenous contraction rate are sustained. These data are consistent with DM-hCO maintaining a stable maturation cellular state. This may be a key adaptation during cardiac maturation to enable large increases in demand from basal function to increased function during exercise or stressed conditions.

One key feature of DM-hCOs is a more consistent rate across different hPSC lines. This made it possible to 1) assess SR function under reduced If in >90% of DM-hCOs, 2) assess ectopy as a surrogate readout of arrhythmia for SR protein mutations, and 3) reveal a stiffening phenotype in DSP mutant DM-hCOs. Automaticity in hPSC-CM results from relatively high expression of HCN4, CACNA1H and SLC8A1, with low expression of KCNJ2^34^. We show that DM-hCOs have a reduction in contraction rate via multiple alternative mechanisms including a more stable SR, a switch from *CACNA1D* to *CACNA1C*, and increased CX43 expression as key changes consistent with human heart maturation. This DM-hCO protocol may therefore also have some applications in cellular therapies to help facilitate a reduction in arrhythmic risk conferred by automaticity^34^.

One of the key features of the DM-hCO protocol is the cellular complexity, which was critical for revealing the *DSP* phenotype. This *DSP* mutation induced a pro-fibrotic signature underpinned by TGF- β1 and kinetic dysfunction in DM-hCOs. TGF-β1 and IL-10 are anti-inflammatory^63^, and increased expression may serve as a cardio-protective response to the inflammation caused by desmosome breakdown. In *DSP* cardiomyopathy patients often present with multiple inflammatory “myocarditis” episodes that cause cardiac remodelling and eventually lead to decreased heart function^64^. This indicates that the *DSP* mutation may be an initiating event that drives immune and fibroblast driven pathological remodelling leading to decreased function and arrhythmia. Indeed, myocardial injury is one of the major risk factors for arrhythmia and heart failure for *DSP* probands^65^. Further, an analogous structural remodelling basis for heart disease has recently been proposed to be a key determinant of arrhythmia in Brugada syndrome^66^. Therefore, complex, multicellular hPSC-models such as the DM-hCOs presented here provide an important tool to identify the drivers of cardiac pathology and identify preventative therapeutics. The utility of this platform is illustrated by showing that bromodomain and extra-terminal protein inhibitors may be effective treatments for both *DSP-* cardiomyopathy and related myopathies with disruptive inflammatory signalling and fibrosis.

## Methods

### hPSC Cell Lines and Culture

#### hPSC Lines

Ethical approval for the generation and/or use of human pluripotent stem cells (hPSCs) were obtained from QIMR Berghofer’s Ethics Committee and Murdoch Children’s Research Institute (MCRI) and were carried out in accordance with the National Health and Medical Research Council of Australia (NHMRC) regulations. hPSCs were obtained from WiCell, Coriell, MCRI and the CIRM hPSC Repository funded by the California Institute of Regenerative Medicine as detailed in the Methods Table 1.

hPSC lines were maintained in mTeSR PLUS (Stem Cell Technologies) in Matrigel coated flasks Matrigel (Millipore) and passaged using ReLeSR (Stem Cell Technologies). Routine quality control was performed for Karyotyping using G-banding (Sullivan Nicolaides) and more recently Molecular Karyotyping Analysis (Victorian Clinical Genetics Service and Ramaciotti Centre for Genomics) and mycoplasma testing. Some HES3 clones had gained a copy number at q11.21, which is likely mosaic in the parental line.

#### Generation of hPSC transgenic lines

For the LED pacing line, neomycin resistance followed by CAG-C1V1 (E122T/E162T) channel rhopdopsin-P2A-super ecliptic pHurlion was cloned into an AAVS1 targeting construct with ∼800 bp homology arms. For the TCF21 reporter line a homologous directed recombination template was synthesized as depicted in the relevant figure. Transgenic cell lines were generated, cloned and quality controlled. A detailed protocol for the generation of the TCF21 line is given below.

Reprogramming and gene-editing factors (Cas9-Gem mRNA, TCF21:CreERT2 template and plasmid encoding TCF21-specific sgRNA) were introduced into perhipheral blood mononuclear cells using the Neon transfection system (1150 V, 30 ms, 2 pulses). Transfected cells were plated over 3 wells of a Matrigel/MEF coated 6-well dish in StemSpan Media. E8 medium was added to each well two days post-transfection and half medium changes were performed every other day until adherent iPSC colonies became visible (∼1 week post-transfection) when full E8 media changes were performed. Individual iPSC colonies were isolated and expanded in E8 medium.

Successfully edited cells were identified by PCR using primers flanking the recombination junctions. Homozygous knock-in of the CreERT2 was confirmed by PCR analysis using primers which flank the target site. A clone harbouring homozygous insertion of CreERT2 was selected for incorporation of the fate-mapper cassette into the *EEF2* locus. Gene-editing factors (Cas9-Gem mRNA, EEF2 fate-mapper HDR template and plasmid encoding EEF2-specific sgRNA) were introduced into iPSCs using the Neon transfection system (1100 V, 30 ms, 1 pulse). Transfected cells were plated over 4 wells of a Matrigel coated 6-well dish in mTesR medium supplemented with 10 μM Y-27632 which was removed from medium after 24 h. Successfully edited iPSC colonies (expressing EYFP) were identified by fluorescence microscopy. EYFP+ colonies were isolated and expanded in E8 medium. A clone with a high proportion of EYFP+ cells was selected for subcloning to attain a pure population. Subcloning was performed by dissociating cells with TryPLE and plating at low density in mTesR supplemented with Y-27632 (removed from medium after 24 h). Individual colonies were picked and expanded in E8 medium. One of these clones was confirmed by flow cytometry to consist entirely of EYFP+ cells and showed homozygous knock-in of the EEF2 fatemapper transgene as evidenced by PCR analysis using primers flanking the *EEF2* target site. sgRNAs are details are provided in the relevant figures.

#### Generation of CASQ2^-/-^ and RYR2^+/N4104K^ hPSC lines

Two hours prior to electroporation, media on the PSCs was changed to fresh mTeSR Plus supplemented with 10 µM ROCK inhibitor Y-27632 (Stem Cell Technologies). For the CASQ2-/- mutation, sgRNAs (Synthego) was conjugated to TrueCut Cas9 Protein v2 (Thermo Fisher) in a 9:1 molar ratio of sgRNA:Cas9 in a 10 µL reaction. For the RyR2^+/N4104K^ cell line, 1.0 µL of 60 µM Alt-R CRISPR-Cas9 sgRNA (Integrated DNA Technologies) was conjugated to 1.0 μL of 3 mg/mL TrueCut Cas9 Protein v2, in a 10 µL reaction. Cells were passaged with ReLeSR, 1.6 x 10^5^ cells re-suspended in Buffer R (12 μL) with the the sgRNA-Cas9 RNP prepared above. For RyR2^+/N4104K^ cell line, 0.6 μL of each 100 μM ssODN was added. sgRNAs and ssODN details are provided in the relevant figures.

The Neon Transfection System 10 μL Kit (mCell Technologies) was used according to the manufacturer’s instructions. Electroporation settings were 1200V, with 1 pulse, and 30 ms pulse width. Cells were immediately plated onto 24-well tissue culture plates coated with Matrigel, containing mTeSR Plus with 10 µM ROCK inhibitor Y-27632, or 1X CloneR2 (Stem Cell Technologies). Post-electroporation survival was assessed the next day and media changed to mTeSR Plus without further supplements. Media was changed every two to three days with mTeSR Plus until PSCs reached 70% confluency and subsequently cryopreserved with mFreSR (Stem Cell Technologies) upon reaching 70% confluency. Concurrently, hPSCs were also harvested for DNA with the DNeasy Blood & Tissue Kit (Qiagen) according to the manufacturer’s instructions.

Amplicons of target sites were generated from DNA of electroporated hPSCs with Hifi Platinum Taq DNA Polymerase High Fidelity (Thermo Fisher Scientific) according to manufacturer’s instructions. PCR purification was performed with the QIAquick PCR Purification Kit (Qiagen) according to the manufacturer’s instructions. Sanger sequencing was performed by the QIMR Berghofer Analytical Facility. The sequence traces from the wild-type parental cell lines and the electroporated hPSC populations were analysed with Synthego’s ICE Analysis tool (available at https://ice.synthego.com/) to determine editing efficiency and incorporation rate of ssODNs.

hPSC pools that had a rate of successful edits greater than 1% were thawed into T25 flasks for subsequent single-cell cloning. Cloning Medium was prepared by supplementing mTeSR Plus with 10% of CloneR or CloneR2. Cells were passaged using ReLeSR and sorted using a BD FACSARIA IIIu Cell Sorter (BD Biosciences) into 96-well plates at a density of 1 cell per well. Culture continued in CloneR2 Cloning Medium, and once all colonies were over 25% confluency they were each passaged with ReLeSR into one well of a 96-well plate and 24-well plate and then maintained in mTeSR Plus. The Extract-N-Amp for Blood Kit (Sigma-Aldrich) was used according to the manufacturer’s instructions to generate DNA lysates from the 96-well plate, and to produce PCR amplicons of the edited sites. PCR amplicons were purified with the QIAquick PCR Purification Kit and submitted for Sanger Sequencing as above. For each clone the Sanger chromatograms were manually interrogated to determine the presence and mono- or bi- allelic editing. hPSC clones were maintained in 24-well plates until genotyping was complete. Positive clones were expanded and banked, negative clones were discarded.

For each gRNA sequence, the top five non-overlapping sites predicted by COSMID (available at https://crispr.bme.gatech.edu/) and CRISPOR (available at http://crispor.tefor.net/) were amplified with PCR and Sanger sequenced. Traces for each off-target site from clones were manually compared to traces from the relevant parental cell line. Clonal edited hPSC cell lines were also karyotyped as described above.

#### Generation of DSP mutant and corrected hPSC lines

Episomal reprogramming plasmids and gene-editing factors were introduced into patient PBMCs (MCRI Biobank ID; MCHTB11.9) using the Neon transfection system (1150 V, 30 ms, 2 pulses). Transfected cells were plated over 3 wells of a Matrigel 6-well dish Essential 8 medium. Media changes were performed every other day. Individual iPSC colonies were picked and expanded in Essential 8 medium. Individual iPSC colonies were genotyped by PCR using primers (AGAAAACGCCCTTCAGCAA and CTCCAGCTTCTTCCTCTTGC) flanking the patient-specific mutation. Amplicons were Sanger sequenced to confirm gene-correction and absence of indel mutations.

### hCO formation and culture

#### Cardiac differentiation and hCO culture

The single step cardiomyocyte and stromal cell differentiation protocol, dissociation and hCO formation was performed as recently described^5,67^.

For pacemaker differentiation, hPSC were differentiated using RPMI base media (Thermo Fisher Scientific) supplemented with 1% penicillin/streptomycin (Thermo Fisher Scientific), 200 mM L-Ascorbic acid 2-phosphate sesquimagnesium salt hydrate (Sigma), and B27 (insulin minus) supplement (Thermo Fisher Scientific). For mesodermal induction, 5 ng/ml BMP4, 9 ng/ml Activin A, 5ng/ml bFGF (all RnD Systems), 1 μM CHIR99021 (Stem Cell Technologies) was added on each day for the first 3 days of differentiation. On day 4 of differentiation 5 μM IWP4 (Stem Cell Technologies), 2.5 ng/ml BMP4, 5.4 mM SB431542 (Sigma), 0.24 mM all-trans retinoic acid (Stem Cell Technologies), 0.5 mM PD173074 (Tocris) was added to RPMI base medium to pattern the cardiac cells. B27 (plus insulin) RPMI basal medium supplemented with 5 μM IWP4 was added for the next 7 days with media changes every 2/3days. On day 15 of differentiation B27 (insulin minus) DMEM no glucose, no glutamine, no phenol red (Thermo Fisher Scientific) supplemented with 5 mM lactic acid (Sigma) was added for the next 7 days with media changes every 2/3 days. On day 22 of differentiation, cells were dissociated and hCO formed as recently described^5,67^.

#### Pacing conditions

hCOs were paced using different methods. Isoprenaline (Sigma) (Extended Data 1). They were exposed to pulses of green light using a 96-well green LED array (Lumidox) connected to a Panlab/Harvard Apparatus Digital Stimulator as recently described^68^ (Extended Data 3). The Heart-Dyno inserts were also fabricated in a custom plate format to fit into a 24 well plate for acute pacing using a C-Pace Cell Culture EP Stimulator (Ionoptix) using 10 V with 1ms pulses at 120 bpm (Fig 1).

#### Maturation factors

Various factors were added at the times and concentrations outlined in the different figures. Small molecules GSK4716, DY131, MK8722, O304 were purchased from MedChem Express. Interferons IFN- γ, IFN-λ1, IFN-λ2, IFN-β and IFN-ω were purchased from Peprotech. Fatty acids palmitic acid, oleic acid, myristic acid and linoleic acid were purchased from Sigma.

### Analysis of hCO

#### Force analysis

hCOs were imaged under environmentally controlled conditions at 37°C with 5% CO2 on a Leica Thunder microscope. Image series were taken of each hCO at 50 frames per second generally for 10 seconds, but up to 40 seconds for experiments using cilobradine. Contraction analysis was performed using custom Matlab scripts^67^ or Tempo.ai analysis software developed by Dynomics.

For calcium sensitivity experiments Tyrode’s solution (120 mM NaCl, 5 mM KCl, 22.6 mM NaHCO3, 2 mM MgCl2 and pH adjusted to 7.4) was used and calcium concentration adjusted using a 0.2M CaCl2 stock solution.

To assess post-rest potentiation, hCOs were pre-treated with 1 µM cilobradine (and in some cases 5 µM thapsigargin) for 2 hours before the experiment. For experiments with acutely paced hCO using custom designed gold plated electrode lids. Pacing was performed at 30-40 mA with 5 ms square pulses for at least 30 seconds prior to recordings.

#### qPCR

hCO were manually homogenised in 500 µL Trizol (Thermo Fisher Scientific) with a stainless-steel ball bearing and 10 sec vortex pulses until solubilised. Samples were stored at −80°C before RNA was extracted using Trizol as per manufacturer’s instructions. RNA was DNase treated as per the manufacturer’s instruction (Roche) before cDNA synthesis (Thermo Fisher Scientific). Powerup SYBR Green Master Mix (Thermo Fisher Scientific) was used for RT-qPCR using StepOne software v2.3 to determine gene expression.

#### Immunostaining

hCOs were fixed with 1% paraformaldehyde (Sigma) for 60 min. Cells were stained with primary antibodies (see Methods Table 1) in 5% FBS (Thermo Fisher Scientific) and 0.2% Triton X-100 (Sigma) in PBS (Blocking Buffer) at 4°C overnight on a rocker. Cells were washed 2X for 1 h with Blocking Buffer and labelled with secondary antibodies (see Methods Table 1) and Hoechst33342 at 4°C overnight on a rocker. Cells were again washed with Blocking Buffer 2X for 1 h and imaged in the Heart-Dyno or mounted on microscope slides in ProLong Glass (ThermoFisher Scientific). Low magnification images were taken on a Leica Thunder microscope. High magnification images were taken using Zeiss 780- NLO Point Scanning Confocal or a Leica Stellaris 5.

#### Drug screening

Individual compounds were purchased from Sigma to form our boutique drug libraries. These were added to hCO for at least 15 min for equilibration prior to video recordings. For concentration-response curves, cumulative drug additions were applied to the same hCO.

#### Transmission Electron Microscopy

hCOs were fixed in 2.5% glutaraldehyde in PBS for 1 hour at room temperature and then processed for embedding *in situ* as described previously^69^. For electron tomography 200-300 nm sections were prepared parallel to the base of the plate on a Leica Ultracut 6 ultramicrotome. The grid was then coated with a thin carbon layer. Tomography was performed as described previously^70^ on a Tecnai F30 transmission electron microscope (FEI) at 300 kV. A dual-axis tilt series spanning ± 60° with 1° increments was acquired with a Gatan OneView camera under the control of Serial EM. Tilt series were reconstructed using IMOD (https://bio3d.colorado.edu/imod/) with segmentation performed by density thresholding using the Isosurface Render program in IMOD.

#### DSP proteomics

Each sample contained 3 hCOs and stored (-20°C) prior to preparation. Samples were thawed on ice, and residual supernatant removed prior to addition of 150 μl chilled lysis buffer (4% sodium deoxycholate in TRIS-buffered saline). Samples were immediately boiled, 95°C, for 5 min to inactivate endogenous enzymatic activity and assist lysis. Four volumes of chilled acetone was added to each sample, and sonicated in 1 min bouts alternating with incubation on ice for a total of 5 cycles. Protein was pelleted by centrifugation at 20,000 x g for 30 min at 4°C. Supernatant was removed, and protein pellet washed twice in acetone prior to resuspending in 150μL 50mM TEAB (Triethylammonium bicarbonate buffer, Sigma). Reduction alkylation buffer was added to a final concentration of 10mM TCEP (Sigma) and 40 mM 2-CAA (Sigma, equilibrated with KOH to pH 8) and incubated for 10min 45°C with 1500 rpm agitation. 3 µg Trypsin enzyme mix (Thermo Fisher Scientific) was added to sample and incubated overnight (17 hours) at 37°C, with 1500 rpm agitation. 50 μl of 10% Trifluoroacetic acid solution was added to each sample to halt digest. Peptides were desalted using SDB-RPS tips, according to standard protocol^71^. Samples were resuspended in 2% ACN/0.3% TFA, with sonication to assist solubility.

For each sample, a uniform volume (3 µl) was resolved across a 65 min gradient in data-independent acquisition mode using a Thermo PepMap100 analytical column equipped on a Thermo Ultimate 3000 LC interfaced with a Thermo Exactive HF-X mass spectrometer. Single injection DIA method utilised parameters established previously^72^. Briefly, MS1 included survey scans of 1e6 ions at resolution of 60,000, with a max isolation time of 60ms within the scan range of 390-101m/z and collision energy of 27. MS2 was performed with isolation windows of 12m/z spanning 400-1000m/z range, AGC target of 1e6 and resolution of 15,000.

#### CASQ2-KO proteomics

PB006.6 wild-type and CASQ2-KO hCOs were snap frozen on dry ice and stored at −80°C until extraction. 8-16 hCOs were lysed in 300 μl of 1% SDS in Milli-Q water with cOmplete ULTRA Protease Inhibitor Cocktail (Roche). Samples were vortexed, transferred to 2 mL Precellys Lysing Kit tubes and then pulsed 3 times for 30 seconds at 5000 rpm with a Minilys homogenizer (Bertin Technologies). Samples were centrifuged at 8000 xg for 10 minutes to remove bubbles. Transferred to Eppendorf tubes and stored at −20°C. Four volumes of chilled acetone was added to each sample, and sonicated in 1 min bouts alternating with incubation on ice for a total of 5 cycles. Protein was pelleted by centrifugation at 20,000 x g for 30 min at 4°C. Supernatant was removed, and protein pellet was washed twice in acetone prior to resuspending in 1X TBS Buffer (ThermoFisher Scientific).

Sodium deoxycholate was added to protein extracts for improved protein solubility, to a final SDC concentration of 1% (v/v). Reduction alkylation buffer was added to a final concentration of 10mM TCEP (Sigma) and 40 mM 2-CAA (Sigma, equilibrated with KOH to pH 8) and incubated for 10 min 45°C with 1500 rpm agitation. 1 µg Trypsin (ThermoFisher Scientific) was added to sample and incubated overnight (17 hours) at 37°C, with 1500 rpm agitation. 50 μl of 10% Trifluoroacetic acid solution was added to each sample to halt digest. Peptides were desalted using SDB-RPS tips, according to standard protocol^73^. Samples were resuspended in 2% ACN/0.3% TFA, with sonication to assist solubility.

For each sample, a uniform volume (3 µl) was resolved across a 55 min gradient in data-independent acquisition mode using a Thermo PepMap100 analytical column equipped on a Thermo Vanquish Neo UHPLC interfaced with a Thermo Exactive HF-X mass spectrometer. Single injection DIA method utilised parameters established previously^74^. Briefly, MS1 included survey scans of 1e6 ions at resolution of 60,000, with a max isolation time of 60ms within the scan range of 390-1010m/z and collision energy of 27. MS2 was performed with isolation windows of 16m/z spanning 400-1000m/z range, AGC target of 1e6 and resolution of 15,000.**DSP and CASQ2-KO proteomics bioinformatics**

Raw spectra was analysed using DIA-NN, version 1.8.1 against the human proteome (20,399 sequences, downloaded April 19, 2021 from UniProt) using the library-free method. Carbamidomethylation of cysteines and N-term N excision was set as a fixed modification, matching between runs enabled, cross-run normalisation used RT-dependent method, and small-profiling method was used to generate *in silico* library from the human proteome.

For the DSP samples the mean values for each protein was calculated for each condition, and log2 fold change values generated, including p value (two-sided parametric t test) and false discovery rate corrected p value (Benjamini Hochberg method). PCA plots were generated using R (v4.2.2) and ggfortify package (v0.4.16). Within Cytoscape (v3.10.2) a protein-protein interaction network was generated by querying the STRING database (stringApp v2.1.1) with the 4473 proteins detected in the DSP hCO dataset. A confidence score of 0.9 was used, and the maximum number of additional interactors was set to 40. Default edge filters were used. The largest network of 3516 proteins was isolated from all other smaller networks and nodes. An Edge-weighted Spring Embedded layout was used. The Markov clustering algorithm was used to cluster the network and functional enrichment was performed on the 10 largest sub-clusters. Clusters were coloured according to the consensus description from ‘GO Biological Process’ and ‘Reactome Pathways’. Node size and transparency was mapped to log2 fold change between DSP and WT groups. Proteins that were regulated by INCB in the DSP hCOs (p value < 0.05) were given a black border.

For each CASQ2 sample and the WT PB006.6 sample, GAPDH was used as the normalization factor for CASQ2, HRC, RYR2, ATP2A2, PLN, and CALR. Normalized spectral intensities for each protein in the CASQ2 samples were then expressed as fold difference to the respective protein from the WT PB006.6 sample.

#### Phosphoproteomics

Control hCO or hCO treated with 3 μM DY131 and 10 μM MK8722 or pacing at 120 bpm for 5 mins were immediately quenched in ice cold tris-buffered saline. 16 hCO were pooled per condition for phosphoproteomics preparation, based on the EasyPhos protocol^73^. Briefly, samples were suspended in sodium deoxycholate containing buffer (4% SDC in tris-buffered saline, pH 7.5) and heated for 5min at 95°C, with 1500 rpm agitation, to assist lysis and inhibit endogenous enzymatic activity. This was followed by sonication, using five 30 sec on/off intervals in a chilled water bath sonicator. Reduction alkylation buffer was added to a final concentration of 10 mM TCEP (Sigma) and 40 mM 2-CAA (Sigma, equilibrated with KOH to pH 8) and incubated for 10 min 45°C with 1500 rpm agitation. 1 µg Trypsin/Lys-C enzyme mix (Thermo Fisher Scientific) was added to sample and incubated overnight at 37°C, with 1500 rpm agitation. The following day, 75 µl of isopropanol was added to each sample and thoroughly mixed, before adding 25 µl enrichment buffer (48% v/v TFA and 8 mM KH2PO4). 45 mg TiO2 titansphere beads were wash 3 times with 6% TFA/80% ACN, resuspended and 5 mg added to each sample. All samples were incubated for 10 min at 40°C, with 2000 rpm agitation. All samples were moved to new tubes, and centrifuged (2,000 xg for 1 min) to remove non-phosphorylated peptides (supernatant). Beads were washed five times with 1 ml 5% TFA/60% isopropanol wash buffer using centrifugation. A final wash used 0.1% TFA/60% isopropanol. Lastly, 50 µl elution buffer (200 µl of ammonia solution to 800 µl of 40% ACN) was added to beads twice, to collect two subsequent eluents. Phosphopeptides were dried, using a Genevac sample concentrator, approximately 25 min on aqueous setting. Peptides were desalted using SDB-RPS tips, according to standard protocol^71^. Samples were resuspended in 2% ACN/0.3% TFA, with sonication to assist solubility.

For each sample, a uniform volume (5 µl) was resolved across a 65 min gradient in data-dependent acquisition mode using a Thermo PepMap100 analytical column equipped on a Thermo Ultimate 3000 LC interfaced with a Thermo Exactive HF-X mass spectrometer. The mass spectrometer performed survey scans of 3e6 ions at a resolution of 60,000 from 300–1600 m/z. The 10 most abundant precursors from the survey scan with charge state >1 and <5 were selected for fragmentation. Precursors were isolated with a window of 1.6 m/z and fragmented in the HCD cell with NCE of 27. Maximum ion fill times for the MS/MS scans were 50 ms, with a target of 2e4 ions. Fragment ions were analyzed with high resolution (15,000) in the Orbitrap mass analyzer. Dynamic exclusion was enabled with duration 30 sec.

#### Phosphoproteomics bioinformatics

Raw LC-MS data was searched against the reviewed Uniprot human database (20399 sequences, downloaded April 19, 2021) using Sequest HT on the Thermo Proteome Discoverer software (Version 2.3), with matching between runs enabled. Precursor and fragment mass tolerance were set to 20 ppm and 0.05 Da respectively. A maximum of two missed cleavages were allowed. A strict false discovery rate (FDR) of 1% was used to filter peptide spectrum matches (PSMs) and was calculated using a decoy search. Carbamidomethylation of cysteines was set as a fixed modification, while oxidation of methionine, N-terminal acetylation and S/T/Y phosphorylation were set as dynamic modifications, up to 3 phosphosites per peptide.

Differential expression analysis was performed in Proteome Discoverer software, to return log2 fold change values and an adjusted p value. Fold changes were made between electrical pacing or directed maturation conditions to control conditions. Differentially altered phosphoproteins (adj.p < 0.05) were analysed in EnrichR^74^ using the KEGG 2021 Human gene set. All phosphoproteomics graphs were generated in GraphPad Prism, and diagrams made in Inkscape (v1.2.2).

#### Metabolism

Live-cell respiration was measured in real-time using Resipher (Lucid Scientific). hCO were cultured in 96-well microplates (Thermo Fisher Scientific) using hCO culture inserts (height approximately 1.5 mm) that were fabricated by PDMS molding^67^. The Resipher sensor lid (9.4 mm probe length, Lucid Scientific, NS32-94A) was pre-equilibrated with naïve media in a cell-free 96-well microplate at 37 °C and 5% CO2 overnight prior to seeding. After hCO formation, the Resipher sensor lid was transferred to the hCO culture plate and media O2 was measured with an operating height between 1600 and 2100 µm. During subsequent media changes, the sensor lid was stored in pre-equilibrated naïve media in a cell-free microplate. On Day 30 of culture, hCO were treated with DMSO control or 10 µM BAM- 15 (MedChemExpress) for 6 h. Respiration rates of hCO were normalised to cell-free control wells on the same plate, which contained naïve media for each condition (SF or DM culture conditions).

### Transcriptional profiling

#### Bulk RNA-sequencing datasets

For maturation marker analysis bulk RNA-sequencing data from published datasets were collected. To ensure cell compositions did not impact analysis sarcomeric isoform fractions were used for the analysis including MYH7-MYH6, MYL2-MYL7 and TNNI3-TNNI1. Data were collated from hPSC-CM cultures including GSE93841^4^, GSE148025^10^, GSE116464^11^, GSE201437^17^, GSE114976^15^ and human hearts including ERP109940^75^ and Hahn et al., 2021^76^.

#### hCO nuclei isolation for snRNA-sequencing

hCO were matured using the SF or DM protocol. At the conclusion of the experiment, media was aspirated and hCO were washed with ice cold PBS^-^. Intact hCO were pooled with up to 70 hCO per sample (n = 2 per maturation protocol). All PBS^-^ was removed and hCO were snap frozen in liquid nitrogen and stored at −80°C. Nuclei isolation was completed using the 10x Chromium Nuclei Isolation Kit (PN-1000494, 10x Genomics) and 10x Genomics Chromium Nuclei Isolation Kit Protocol for Single Cell 3’ Gene Expression (RevA) with modifications. All subsequent steps in the nuclei isolation steps were performed on ice. The samples were homogenised in 200 μl Lysis Reagent using the pestle supplied in the 10x kit. 300 μl Lysis Reagent was added and samples were pipette mixed and left to incubate on ice for 5 minutes. The sample was then transferred into a pre-chilled Nuclei Isolation Column and centrifuged at 16 000 x g for 20 sec at 4°C. Samples were vortexed for 5 seconds at max speed to resuspend nuclei and centrifuged again at 500 x g for 5 min at 4°C. 300 μl of supernatant was removed and nuclei re-suspended in 500 μl of Debris Removal Buffer. Samples were centrifuged at 700 x g for 10 min at 4°C. 500 μl of supernatant was removed and nuclei were gently re-suspended in 1 ml of Wash and Resuspension Buffer. Samples were centrifuged at 500 x g for 5 min at 4°C. Nuclei pellets were then re-suspended in 250 µl Wash and Resuspension Buffer containing 4,6-diamidino-2- phenylindole (DAPI, Thermo Fisher Scientific). Samples were sorted on the FACSAria III Cell Sorter (BD Biosciences) using a 70 µM nozzle. Nuclei were captured in 1.5 ml Eppendorf tubes containing 1 ml Wash and Resuspension Buffer. Recovered nuclei were centrifuged again at 500 x g for 5 min at 4°C. The supernatant was removed to leave ∼100 µl and nuclei were resuspended by pipette mixing. Nuclei density counting was completed on the Countess III FL (Thermo Fisher Scientific).

Nuclei were processed using the Chromium Next GEM Single Cell 3’ GEM, Library & Gel Bead Kit v3.1 (PN-1000128, 10X Genomics). Nuclei were loaded into a Chip G (PN-1000127, 10X Genomics) and run on the Chromium Controller (10X Genomics) for gel bead emulsion (GEM) formation. Reverse transcription, barcoding, complementary DNA amplification and purification for library preparation were performed according to the Chromium Single Cell 3’ Reagent Kits User Guide (CG000204_ChromiumNextGEMSingleCell3’v3.1_Rev). GEM formation and library preparation was completed by the Sequencing Facility at the Institute of Molecular Biosciences. All libraries were pooled and sequenced across two NovaSeq 6000 S1 flow cells (Illumina) to a depth of ∼50,000 reads per nuclei (∼10,000 per sample).

#### snRNA-sequencing bioinformatics

For human fetal, young, and adult hearts snRNAseq Fastq files were obtained from GEO accession GSE156707^3^. Fastq files for human and hCO samples were aligned using 10x Cell Ranger (version 7.0). The cellranger count command with default parameters was used to align the sequencing reads to the GRCh38 build of the human transcriptome (refdata-gex-GRCh38-2020-A) and generate a gene expression count matrix.

Sample filtering and quality control analysis: All subsequent analysis was performed using the R statistical programming language, using the Seurat package (version 4.3.0.1). There was an initial filtering step to keep genes that were expressed in 3 or more nuclei and nuclei with at least 200 detected genes. The quality of the cells was assessed for each sample independently by examining the total number of nuclei, the distributions of total unique molecular identifier (UMI) counts, the number of unique genes detected, and the proportions of ribosomal and mitochondrial content per nuclei. Nuclei displaying high expression of mitochondrial genes (using a cut-off of < 5%) were removed. For the 9 human samples, nuclei with a UMI count depth of under 1,000 and higher than 40,000 were filtered out to remove debris and potential doublets. For the 4 hCO samples, nuclei with a UMI count depth of under 1,000 and higher than 30,000 were filtered out. No filtering was applied based on nFeature_RNA. SoupX (version 1.6.2) was used to estimate ambient RNA from empty droplets and correct the expression metrics by removing counts related to ambient RNA molecules.

Normalisation, integration and clustering: For each Seurat object, transformation and normalisation was performed using SCTransform to fit a negative binomial distribution and regress out mitochondrial read percentage. Integration of the 4 samples was performed using the following functions with default parameters: SelectIntegrationFeatures, FindIntegrationAnchors, IntegrateData. Principle components (PCs) were then calculated using the RunPCA function, and an elbow plot generated to select the cut-off for significant PCs to use for downstream analysis. UMAP dimensional reduction was then computed using the top 20 PCs using the RunUMAP command. Unsupervised clustering was then performed using the FindNeighbors and FindClusters functions. For exploratory purposes, Leiden clustering with a resolution range of 0 – 1 increasing at increments of 0.1 was performed to identify clusters within the data.

Differential gene expression testing: Differential gene expression analysis was performed using the FindMarkers command using the default Wilcoxon rank-sum. The logFC_cutoff was set to 0.25, assay set to ‘SCT’, and slot set to ‘data’. Mitochondrial genes were excluded from differential expression analysis. P-value adjustment was performed using bonferroni correction based on the total number of genes in the dataset. clusterProfiler (version 4.2.2) was used for enrichment analysis of gene ontology (GO) terms, using the enrichGO function. Adjusted p-values were calculated by the BH method and results were visualised using the dotplot function from enrichplot (version 1.14.2).

#### Spatial RNA-sequencing

hCOs were washed in PBS and evenly coated in pre-chilled Tissue-Tek O.C.T (Sakura), then placed on a 1 x 1cm plastic mold containing OCT. hCOs were arrayed in different orientations to contain 6 SF- hCO and 6 DM-hCO embedded in Tissue-Tek O.C.T and frozen in an ethanol slurry. Tissue sections were sectioned at 10 μm thickness using a CM3050S cryostat (Leica), mounted onto a Stereo-seq T-Chip slide (Stereo-seq, 210CT114) pre-coated with 0.01% poly-L-lysine (Sigma) and allowed to incubate for 5 min at 37°C. Tissue sections were then fixed in methanol for 30 min at −20°C then stained with ssDNA fluorescent staining solution to identify cell boundaries and imaged using Z2 Axio Imager (Ziess) using the FITC channel. The sections were permeabilised using 1 x Permeabilsation Reagent Solution (Stereo-seq Transcriptomics T Kit, 111KT114) for 18 minutes at 37°C, then rinsed in 1x PR Rinse Buffer.

RNA was released and reverse transcribed using Reverse Transcription Mix (Stereo-seq Transcriptomics T Kit, 111KT114) at 42°C for 3 hours. Following reverse transcription, hCOs were digested with TR Buffer (Stereo-seq Transcriptomics T Kit, 111KT114) at 55°C for 10 minutes. To release cDNA from the chip, samples were incubated with cDNA Release Mix for 16 hours at 55°C. cDNA was collected, purified using 0.8X AMPure XP Beads (Beckman Coulter) then amplified using cDNA Amplification Mix (Stereo-seq Transcriptomics T Kit, 111KT114). Samples were incubated at 95°C for 5 min, then 15 cycles of 98°C for 20 secs, 58°C for 20 sec, 72°C for 3 min and a final elongation step at 72°C for 5 min.

DNA concentration was quantified using Qubit™ dsDNA Assay Kit (Thermo Fisher Scientific) and cDNA fragmentation was carried out using 20 ng of cDNA and 1X Fragmentation Reaction Mix (Stereo-seq Library Preparation Kit, 111KL114) at 55°C for 10 min. Fragmented product was amplified using PCR Barcode Primer and Amplification Mix (Stereo-seq Library Preparation Kit, 111KL114) using the following thermal cycler settings: 95°C for 5 min, 13 cycles of 98°C for 20 sec, 58°C for 20 sec, 72°C for 3 min and a final step at 72°C for 5 min. The PCR product was purified using 0.55X AMPure XP Beads and sequenced using MGI DNBSEQ™-T7 sequencer.

#### Spatial RNA-sequencing bioinformatics

The stitched ssDNA image was first processed through ImageStudio for compatibility with the SAW pipeline. The single Stereo-seq chip was first processed using the SAW pipeline v6.12 and aligned to the human reference genome GRCh38, using the default parameters including cellbinning. The tissue.gef output file was then analyzed using Stereopy v1.1.0. For each maturation protocol, three of the six available dynos with the highest data quality were selected and segregated into individual data objects using spatial coordinates. Each of these dynos were then analysed separately at bin size 10 and filtered for percentage of mitochondrial genes per bin, counts per bin, and genes per bin. Data was normalized for total counts and log transformed before performing Gaussian smoothing.

## Statistical analysis

Unless otherwise indicated each n is designated as an individual hCO, cultured independently for the entire period following hCO formation. An experiment is designated as an independent cardiac differentiation performed on an entirely different week from a different passage number of hPSCs.

For functional data, in most cases pre-treatment baseline functional recordings of hCO were performed and post-treatment recordings normalized to these as the baseline. An additional normalization to the control conditions was also performed in many cases to account for any time-dependent changes in function.

Most data were analyzed using Prism v8.2 (GraphPad) with the appropriate statistical tests depending on normality, equal variances, experimental setup and required comparisons. The tests applied and the p-values are given in each figure and legend. For omics data, data were analyzed as described in the bioinformatics sections.

## Methods Table 1: Critical resources

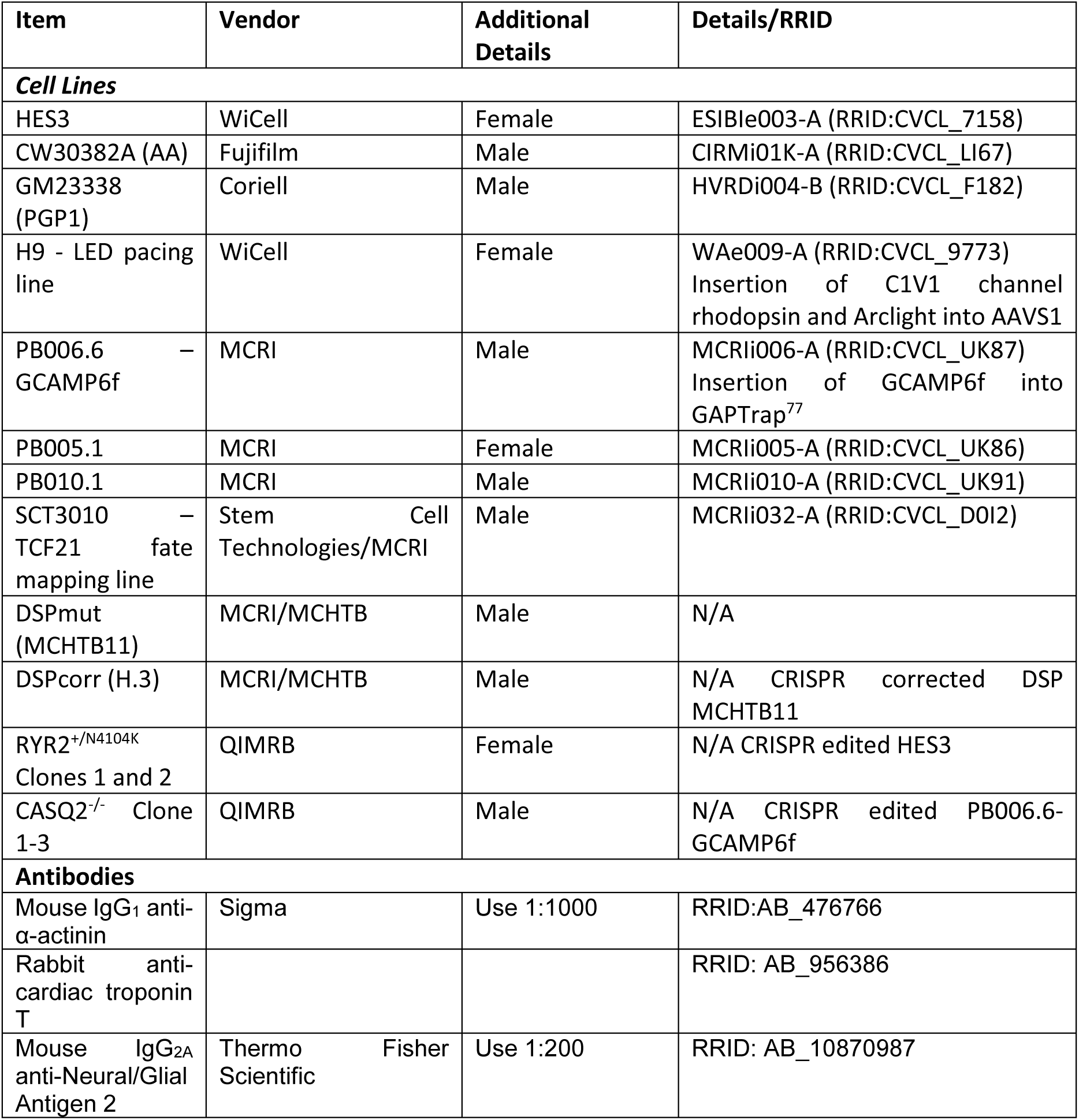
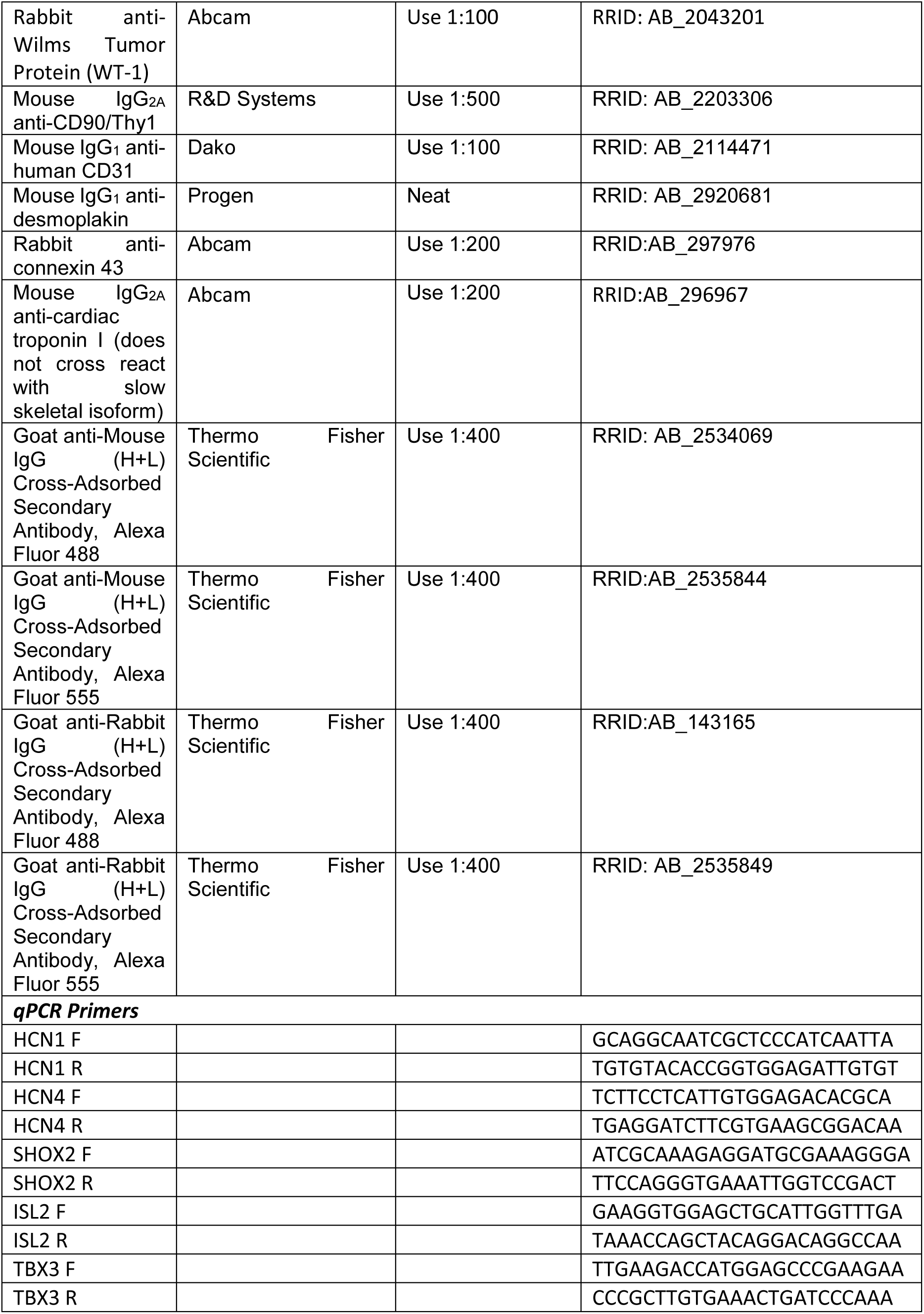
Methods Table 1: Critical resources.

## Acknowledgements

The Heart-Dyno molds were fabricated using the NCRIS enabled Australian National Fabrication Facility – Queensland, New South Wales and South Australian (Government of South Australia supported) Nodes. We especially thank Simon Doe and Mark Cherrill.

We thank Tam Nguyen and Nigel Waterhouse for microscopy, Tu Parsons and Paul Collins for Sanger sequencing, and Grace Chojnowski and Michael Rist for FACS (QIMR Berghofer). Mark Hodson and Rebekah Ziegman for proteomics support. Charles Ferguson for assistance with transmission electron microscopy. Chris Semsarian for cardiomyopathy genetics.

J.E.H. was supported by funding from a Snow Medical Fellowship and J.R.K. QIMRB funding. E.R.P. is supported by an Investigator Grant from the National Health and Medical Research Council of Australia (GNT2008376). The Novo Nordisk Foundation Center for Stem Cell Medicine (E.R.P. and R.J.M.) is supported by Novo Nordisk Foundation grants (NNF21CC0073729). R.G.P. is supported by an Australian Research Council (ARC) Laureate Fellowship (FL210100107). This was was also supported by the STOmics Grant Program.

The services and technical assistance of the Murdoch Children’s Research Institute iPSC & GE Core Facility. This facility was established using a generous donation from the Stafford Fox Medical Research Foundation and is currently supported by Phenomics Australia (PA), and the Novo Nordisk Foundation reNEW Center for Stem Cell Medicine (NNF21CC0073729). PA is supported by the Australian Government through the National Collaborative Research Infrastructure Strategy program. The use of the Microscopy Australia Research Facility at the Centre for Microscopy and Microanalysis at The University of Queensland.

## Author contributions

M.P., R.J.M. and J.E.H. conceived the project, all authors contributed to the acquisition and analysis of data, all authors contributed to the interpretation of data, J.E.H. wrote the original manuscript, all authors revised the manuscript.

## Materials & Correspondence

James Hudson

## References

1 Guo, Y. & Pu, W. T. Cardiomyocyte Maturation: New Phase in Development. Circ Res 126, 1086–1106, doi:10.1161/circresaha.119.315862 (2020).

2 Mills, R. J. & Hudson, J. E. Bioengineering adult human heart tissue: How close are we? APL Bioeng 3, 010901, doi:10.1063/1.5070106 (2019).

3 Sim, C. B. et al. Sex-Specific Control of Human Heart Maturation by the Progesterone Receptor. Circulation 143, 1614–1628, doi:10.1161/circulationaha.120.051921 (2021).

4 Mills, R. J. et al. Functional screening in human cardiac organoids reveals a metabolic mechanism for cardiomyocyte cell cycle arrest. Proc Natl Acad Sci U S A 114, E8372–e8381, doi:10.1073/pnas.1707316114 (2017).

5 Voges, H. K. et al. Vascular cells improve functionality of human cardiac organoids. Cell Rep 42, 112322, doi:10.1016/j.celrep.2023.112322 (2023).

6 Karbassi, E. et al. Cardiomyocyte maturation: advances in knowledge and implications for regenerative medicine. Nat Rev Cardiol 17, 341–359, doi:10.1038/s41569-019-0331-x (2020).

7 Bedada, F. B. et al. Acquisition of a quantitative, stoichiometrically conserved ratiometric marker of maturation status in stem cell-derived cardiac myocytes. Stem Cell Reports 3, 594–605, doi:10.1016/j.stemcr.2014.07.012 (2014).

8 Drakhlis, L. et al. Human heart-forming organoids recapitulate early heart and foregut development. Nat Biotechnol 39, 737–746, doi:10.1038/s41587-021-00815-9 (2021).

9 Lewis-Israeli, Y. R. et al. Self-assembling human heart organoids for the modeling of cardiac development and congenital heart disease. Nat Commun 12, 5142, doi:10.1038/s41467-021-25329-5 (2021).

10 Hofbauer, P. et al. Cardioids reveal self-organizing principles of human cardiogenesis. Cell 184, 3299–3317.e3222, doi:10.1016/j.cell.2021.04.034 (2021).

11 Giacomelli, E. et al. Human-iPSC-Derived Cardiac Stromal Cells Enhance Maturation in 3D Cardiac Microtissues and Reveal Non-cardiomyocyte Contributions to Heart Disease. Cell Stem Cell 26, 862–879.e811, doi:10.1016/j.stem.2020.05.004 (2020).

12 Tiburcy, M. et al. Defined Engineered Human Myocardium With Advanced Maturation for Applications in Heart Failure Modeling and Repair. Circulation 135, 1832–1847, doi:10.1161/circulationaha.116.024145 (2017).

13 Schaaf, S. et al. Human engineered heart tissue as a versatile tool in basic research and preclinical toxicology. PLoS One 6, e26397, doi:10.1371/journal.pone.0026397 (2011).

14 Zhang, D. et al. Tissue-engineered cardiac patch for advanced functional maturation of human ESC-derived cardiomyocytes. Biomaterials 34, 5813–5820, doi:10.1016/j.biomaterials.2013.04.026 (2013).

15 Zhao, Y. et al. A Platform for Generation of Chamber-Specific Cardiac Tissues and Disease Modeling. Cell 176, 913–927.e918, doi:10.1016/j.cell.2018.11.042 (2019).

16 Ronaldson-Bouchard, K. et al. Advanced maturation of human cardiac tissue grown from pluripotent stem cells. Nature 556, 239–243, doi:10.1038/s41586-018-0016-3 (2018).

17 Shen, S. et al. Physiological calcium combined with electrical pacing accelerates maturation of human engineered heart tissue. Stem Cell Reports 17, 2037–2049, doi:10.1016/j.stemcr.2022.07.006 (2022).

18 Raad, F. S. et al. Chalcone-Supported Cardiac Mesoderm Induction in Human Pluripotent Stem Cells for Heart Muscle Engineering. ChemMedChem 16, 3300–3305, doi:10.1002/cmdc.202100222 (2021).

19 Simon Sarkadi, L., et al. Fatty Acid Composition of Milk from Mothers with Normal Weight, Obesity, or Gestational Diabetes. Life (Basel) 12, doi:10.3390/life12071093 (2022).

20 Zeleniuch-Jacquotte, A., Chajès, V., Van Kappel, A. L., Riboli, E. & Toniolo, P. Reliability of fatty acid composition in human serum phospholipids. Eur J Clin Nutr 54, 367–372, doi:10.1038/sj.ejcn.1600964 (2000).

21 Arad, M., Seidman, C. E. & Seidman, J. G. AMP-activated protein kinase in the heart: role during health and disease. Circ Res 100, 474–488, doi:10.1161/01.Res.0000258446.23525.37 (2007).

22 Pedram, A. et al. Estrogen regulates histone deacetylases to prevent cardiac hypertrophy. Mol Biol Cell 24, 3805–3818, doi:10.1091/mbc.E13-08-0444 (2013).

23 Solan, J. L. et al. Phosphorylation at S365 is a gatekeeper event that changes the structure of Cx43 and prevents down-regulation by PKC. J Cell Biol 179, 1301–1309, doi:10.1083/jcb.200707060 (2007).

24 Sheikh, F. et al. Mouse and computational models link Mlc2v dephosphorylation to altered myosin kinetics in early cardiac disease. J Clin Invest 122, 1209–1221, doi:10.1172/jci61134 (2012).

25 Traaseth, N. J. et al. Structural and dynamic basis of phospholamban and sarcolipin inhibition of Ca(2+)-ATPase. Biochemistry 47, 3–13, doi:10.1021/bi701668v (2008).

26 Mills, R. J. et al. Drug Screening in Human PSC-Cardiac Organoids Identifies Pro-proliferative Compounds Acting via the Mevalonate Pathway. Cell Stem Cell 24, 895–907.e896, doi:10.1016/j.stem.2019.03.009 (2019).

27 Fernandes, I., Funakoshi, S., Hamidzada, H., Epelman, S. & Keller, G. Modeling cardiac fibroblast heterogeneity from human pluripotent stem cell-derived epicardial cells. Nat Commun 14, 8183, doi:10.1038/s41467-023-43312-0 (2023).

28 Espinoza-Lewis, R. A. et al. Shox2 is essential for the differentiation of cardiac pacemaker cells by repressing Nkx2-5. Dev Biol 327, 376–385, doi:10.1016/j.ydbio.2008.12.028 (2009).

29 Wang, B. et al. Foxp1 regulates cardiac outflow tract, endocardial cushion morphogenesis and myocyte proliferation and maturation. Development 131, 4477–4487, doi:10.1242/dev.01287 (2004).

30 Saleem, M., Barturen-Larrea, P. & Gomez, J. A. Emerging roles of Sox6 in the renal and cardiovascular system. Physiol Rep 8, e14604, doi:10.14814/phy2.14604 (2020).

31 Vafiadaki, E., Arvanitis, D. A. & Sanoudou, D. Muscle LIM Protein: Master regulator of cardiac and skeletal muscle functions. Gene 566, 1–7, doi:10.1016/j.gene.2015.04.077 (2015).

32 Sakamoto, T. et al. A Critical Role for Estrogen-Related Receptor Signaling in Cardiac Maturation. Circ Res 126, 1685–1702, doi:10.1161/circresaha.119.316100 (2020).

33 Kim, J. J. et al. Mechanism of automaticity in cardiomyocytes derived from human induced pluripotent stem cells. J Mol Cell Cardiol 81, 81–93, doi:10.1016/j.yjmcc.2015.01.013 (2015).

34 Marchiano, S. et al. Gene editing to prevent ventricular arrhythmias associated with cardiomyocyte cell therapy. Cell Stem Cell 30, 396–414.e399, doi:10.1016/j.stem.2023.03.010 (2023).

35 Mannhardt, I. et al. Human Engineered Heart Tissue: Analysis of Contractile Force. Stem Cell Reports 7, 29–42, doi:10.1016/j.stemcr.2016.04.011 (2016).

36 Bogdanov, K. Y., Vinogradova, T. M. & Lakatta, E. G. Sinoatrial nodal cell ryanodine receptor and Na(+)-Ca(2+) exchanger: molecular partners in pacemaker regulation. Circ Res 88, 1254–1258, doi:10.1161/hh1201.092095 (2001).

37 Peters, C. H. et al. Bidirectional flow of the funny current (I(f)) during the pacemaking cycle in murine sinoatrial node myocytes. Proc Natl Acad Sci U S A 118, doi:10.1073/pnas.2104668118 (2021).

38 Pieske, B. et al. Diminished post-rest potentiation of contractile force in human dilated cardiomyopathy. Functional evidence for alterations in intracellular Ca2+ handling. J Clin Invest 98, 764–776, doi:10.1172/jci118849 (1996).

39 Miotto, M. C. et al. Structural analyses of human ryanodine receptor type 2 channels reveal the mechanisms for sudden cardiac death and treatment. Sci Adv 8, eabo1272, doi:10.1126/sciadv.abo1272 (2022).

40 Sutko, J. L. & Willerson, J. T. Ryanodine alteration of the contractile state of rat ventricular myocardium. Comparison with dog, cat, and rabbit ventricular tissues. Circ Res 46, 332–343, doi:10.1161/01.res.46.3.332 (1980).

41 Chiesi, M., Wrzosek, A. & Grueninger, S. The role of the sarcoplasmic reticulum in various types of cardiomyocytes. Mol Cell Biochem 130, 159–171, doi:10.1007/bf01457397 (1994).

42 Xia, Y., Zhang, X. H., Yamaguchi, N. & Morad, M. Point mutations in RyR2 Ca2+-binding residues of human cardiomyocytes cause cellular remodelling of cardiac excitation contraction-coupling. Cardiovasc Res 120, 44–55, doi:10.1093/cvr/cvad163 (2024).

43 Chung, J. H., Canan, B. D., Whitson, B. A., Kilic, A. & Janssen, P. M. L. Force-frequency relationship and early relaxation kinetics are preserved upon sarcoplasmic blockade in human myocardium. Physiol Rep 6, e13898, doi:10.14814/phy2.13898 (2018).

44 Priori, S. G. et al. Clinical and molecular characterization of patients with catecholaminergic polymorphic ventricular tachycardia. Circulation 106, 69–74, doi:10.1161/01.cir.0000020013.73106.d8 (2002).

45 Baig, S. M. et al. Loss of Ca(v)1.3 (CACNA1D) function in a human channelopathy with bradycardia and congenital deafness. Nat Neurosci 14, 77–84, doi:10.1038/nn.2694 (2011).

46 Al-Attar, R. et al. Casein Kinase 1 Phosphomimetic Mutations Negatively Impact Connexin-43 Gap Junctions in Human Pluripotent Stem Cell-Derived Cardiomyocytes. Biomolecules 14, doi:10.3390/biom14010061 (2024).

47 Yavari, A. et al. Mammalian γ2 AMPK regulates intrinsic heart rate. Nat Commun 8, 1258, doi:10.1038/s41467-017-01342-5 (2017).

48 Gintant, G. et al. Repolarization studies using human stem cell-derived cardiomyocytes: Validation studies and best practice recommendations. Regul Toxicol Pharmacol 117, 104756, doi:10.1016/j.yrtph.2020.104756 (2020).

49 Abdelsayed, M., Bytyçi, I., Rydberg, A. & Henein, M. Y. Left Ventricular Contraction Duration Is the Most Powerful Predictor of Cardiac Events in LQTS: A Systematic Review and Meta-Analysis. J Clin Med 9, doi:10.3390/jcm9092820 (2020).

50 Toischer, K. et al. K201 improves aspects of the contractile performance of human failing myocardium via reduction in Ca2+ leak from the sarcoplasmic reticulum. Basic Res Cardiol 105, 279–287, doi:10.1007/s00395-009-0057-8 (2010).

51 Romero, A. et al. Discovery of Nelutroctiv (CK-136), a Selective Cardiac Troponin Activator for the Treatment of Cardiovascular Diseases Associated with Reduced Cardiac Contractility. J Med Chem 67, 7825–7835, doi:10.1021/acs.jmedchem.3c02413 (2024).

52 Rhoden, A. et al. Comprehensive analyses of the inotropic compound omecamtiv mecarbil in rat and human cardiac preparations. Am J Physiol Heart Circ Physiol 322, H373–h385, doi:10.1152/ajpheart.00534.2021 (2022).

53 Dashwood, A. et al. Effects of omecamtiv mecarbil on failing human ventricular trabeculae and interaction with (-)-noradrenaline. Pharmacol Res Perspect 9, e00760, doi:10.1002/prp2.760 (2021).

54 Teerlink, J. R. et al. Dose-dependent augmentation of cardiac systolic function with the selective cardiac myosin activator, omecamtiv mecarbil: a first-in-man study. Lancet 378, 667–675, doi:10.1016/s0140-6736(11)61219-1 (2011).

55 Voors, A. A. et al. Effects of danicamtiv, a novel cardiac myosin activator, in heart failure with reduced ejection fraction: experimental data and clinical results from a phase 2a trial. Eur J Heart Fail 22, 1649–1658, doi:10.1002/ejhf.1933 (2020).

56 Yuan, P. et al. Single-Cell RNA Sequencing Uncovers Paracrine Functions of the Epicardial-Derived Cells in Arrhythmogenic Cardiomyopathy. Circulation 143, 2169–2187, doi:10.1161/circulationaha.120.052928 (2021).

57 Howden, S. E., Thomson, J. A. & Little, M. H. Simultaneous reprogramming and gene editing of human fibroblasts. Nat Protoc 13, 875–898, doi:10.1038/nprot.2018.007 (2018).

58 Mills, R. J. et al. BET inhibition blocks inflammation-induced cardiac dysfunction and SARS-CoV-2 infection. Cell 184, 2167–2182.e2122, doi:10.1016/j.cell.2021.03.026 (2021).

59 Laurent, G. et al. Effects of chronic gap junction conduction-enhancing antiarrhythmic peptide GAP-134 administration on experimental atrial fibrillation in dogs. Circ Arrhythm Electrophysiol 2, 171–178, doi:10.1161/circep.108.790212 (2009).

60 Alexanian, M. et al. A transcriptional switch governs fibroblast activation in heart disease. Nature 595, 438–443, doi:10.1038/s41586-021-03674-1 (2021).

61 Fujiwara, Y. et al. ERRγ agonist under mechanical stretching manifests hypertrophic cardiomyopathy phenotypes of engineered cardiac tissue through maturation. Stem Cell Reports 18, 2108–2122, doi:10.1016/j.stemcr.2023.09.003 (2023).

62 Sakamoto, T. & Kelly, D. P. Cardiac maturation. J Mol Cell Cardiol 187, 38–50, doi:10.1016/j.yjmcc.2023.12.008 (2024).

63 Li, M. O. & Flavell, R. A. Contextual regulation of inflammation: a duet by transforming growth factor-beta and interleukin-10. Immunity 28, 468–476, doi:10.1016/j.immuni.2008.03.003 (2008).

64 Smith, E. D. et al. Desmoplakin Cardiomyopathy, a Fibrotic and Inflammatory Form of Cardiomyopathy Distinct From Typical Dilated or Arrhythmogenic Right Ventricular Cardiomyopathy. Circulation 141, 1872–1884, doi:10.1161/circulationaha.119.044934 (2020).

65 Wang, W. et al. Clinical characteristics and risk stratification of desmoplakin cardiomyopathy. Europace 24, 268–277, doi:10.1093/europace/euab183 (2022).

66 Nademanee, K. et al. Fibrosis, Connexin-43, and Conduction Abnormalities in the Brugada Syndrome. J Am Coll Cardiol 66, 1976–1986, doi:10.1016/j.jacc.2015.08.862 (2015).

67 Voges, H. K., Mills, R. J., Porrello, E. R. & Hudson, J. E. Generation of vascularized human cardiac organoids for 3D in vitro modeling. STAR Protoc 4, 102371, doi:10.1016/j.xpro.2023.102371 (2023).

68 Mills, R. J. et al. Development of a human skeletal micro muscle platform with pacing capabilities. Biomaterials 198, 217–227, doi:10.1016/j.biomaterials.2018.11.030 (2019).

69 Takasato, M. et al. Kidney organoids from human iPS cells contain multiple lineages and model human nephrogenesis. Nature 526, 564–568, doi:10.1038/nature15695 (2015).

70 Ariotti, N. et al. Molecular Characterization of Caveolin-induced Membrane Curvature. J Biol Chem 290, 24875–24890, doi:10.1074/jbc.M115.644336 (2015).

71 Rappsilber, J., Mann, M. & Ishihama, Y. Protocol for micro-purification, enrichment, pre-fractionation and storage of peptides for proteomics using StageTips. Nat Protoc 2, 1896–1906, doi:10.1038/nprot.2007.261 (2007).

72 Pino, L. K., Just, S. C., MacCoss, M. J. & Searle, B. C. Acquiring and Analyzing Data Independent Acquisition Proteomics Experiments without Spectrum Libraries. Mol Cell Proteomics 19, 1088–1103, doi:10.1074/mcp.P119.001913 (2020).

73 Humphrey, S. J., Karayel, O., James, D. E. & Mann, M. High-throughput and high-sensitivity phosphoproteomics with the EasyPhos platform. Nat Protoc 13, 1897–1916, doi:10.1038/s41596-018-0014-9 (2018).

74 Kuleshov, M. V. et al. Enrichr: a comprehensive gene set enrichment analysis web server 2016 update. Nucleic Acids Res 44, W90–97, doi:10.1093/nar/gkw377 (2016).

75 Pervolaraki, E., Dachtler, J., Anderson, R. A. & Holden, A. V. The developmental transcriptome of the human heart. Sci Rep 8, 15362, doi:10.1038/s41598-018-33837-6 (2018).

76 Hahn, V. S. et al. Myocardial Gene Expression Signatures in Human Heart Failure With Preserved Ejection Fraction. Circulation 143, 120–134, doi:10.1161/circulationaha.120.050498 (2021).

77 Kao, T. et al. GAPTrap: A Simple Expression System for Pluripotent Stem Cells and Their Derivatives. Stem Cell Reports 7, 518–526, doi:10.1016/j.stemcr.2016.07.015 (2016).

