## Supplementary Data for "Maturation of human cardiac organoids enables complex disease modelling and drug discovery"

**Extended Data Results**

This section provides an overview of how the maturation factors for the combinatorial screening in Fig. 1 were identified.

*Metabolism regulators for maturation*

We initially explored increasing loading using electrical pacing in our original hCO protocols[1]. Chronic pacing using electrodes in our platform results in media oxidation and hCO death due to the low media volumes and exchanges rates[1]. As our hCOs contain 6% pacemaker cells[1], we used 1 µM isoprenaline (ISO) to chronically increase rate (Extended Data 2). This resulted in a 25% increase in rate (to ~50 bpm), which declined to < 40 bpm over time. Even with this modest rate increase MLC2v was significantly increased. To increase rate more substantially we explored improving the pacemaker capability. We used an adapted nodal cardiomyocyte differentiation protocol[2] and were able to generate 'pacemaker' hCOs with substantially increased rates >150 bpm (Extended Data 3a-e). *In vivo* the sinoatrial nodal cells (SANCs) require semi-insulated 'nodes' to facilitate their automaticity capability, so we added a 5,000 hPSC-derived SANC 'node' to each hCO (Extended Data 3f,g), but this was also unable to achieve greatly enhanced rates.

To directly increase rate using alternative pacing methods similar to our skeletal muscle platform[3], we used optogenetic pacing where we added red shifted channel rhodopsin into the AAVS1 locus (Extended Data 4). While we were able to pace at 60 bpm at the start of the protocol, by the end of the maturation phase we were unable to continue to capture rate at 60 bpm even with the addition of 100 nM ISO (Extended Data 4).

In order to consistently and robustly increase hCO metabolic demand we next explored 'pharmacological exercise'[4] in our new serum-free conditions (SF-hCO)[5]. For metabolism we initially screened small molecules acting on key controllers of metabolism[6,7] including ERRβ/γ (GSK4716[8] and DY131[9]) or AMPK (O304 and MK8722[10]). These influenced different aspects of maturation (Extended Data 5), 10 µM MK8722 increased cTnI protein levels, while 10 µM DY131 increased force with a 2 days treatment at the end of the protocol. O304 may not have had any effect because it inhibits de-activation of AMPK in contrast to direct activation by MK8722.

We then compared MK8722 and DY131 to progesterone which we recently identified as a key metabolic regulator[11]. These were we added for 4 days followed by 4 days of recovery before analysis to assess whether the effects were sustained (Extended Data 6). Both progesterone and DY131 increased force when present, but only progesterone led to a sustained increase in force. This may be due to some overlapping gene control in the NR3C family of which these are both members[12]. MK8722 initially decreased force, which recovered after its removal. MK8722 also led to sustained reduction in rate and increased cTnI levels. Therefore, different metabolic regulators can drive different aspects of cardiac maturation.

*Screening interferons for maturation*

We have previously identified that maturation of non-myocytes, including fibroblasts, endothelial cells and epicardial cells, is associated with increased interferon signalling[11]. To determine if interferons can also drive hCO maturation, we tested IFN-γ, IFN-λ1, IFN-λ2, IFN-β and IFN-ω (Extended Data 7). The different IFN members led to profound differences in hCO response. IFN-γ increased force, reduced rate and transiently increased Tr50, including a 57% increase for 100 ng/ml IFN-γ at 48h ($P < 0.0001$, Mann-Whitney, $n = 21\text{-}28$ from 4 experiments) similar to our previous results[13], which returned to baseline at later time-points. IFN-β and IFN-ω substantially reduced force by 54% and 50%, respectively. In contrast, IFN-λ1 did not negatively impact function, an as such it was included in further screens to assess putative combinatorial effects. Surprisingly, even though cTnI levels were not statistically altered, the expression of cTnI was almost negligible in some of the hCOs treated with IFN-γ, IFN-β and IFN-ω (Extended Data 7h). Therefore, despite there being a key interferon signature of non-myocytes in the maturing heart[11], interferons appear to counteract cardiomyocyte maturation and may be a by-product of activation of the immune system rather than the cause of physiological heart maturation.

*Addition of CHIR99021 for hCO formation*

Prior to full factorial screening, we discovered that addition of 2 μM CHIR99021 during the first 2 days of hCO formation led to increased cellularity and improved consistency between batches as it promotes cell survival and proliferation[14,15] (Extended Data 8). There was a trend to increased force due to increase cell number, decreased rate and slightly elevated Tr50 without any impact on the maturation marker cTnI.

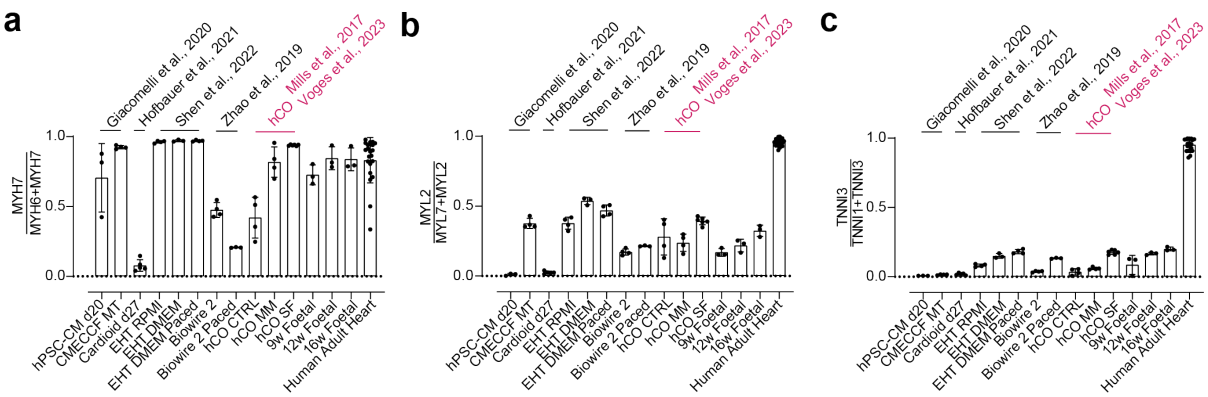


**Extended Data Fig. 1 Comparison of mRNA expression of sarcomeric protein ratios that correlate with maturation across multiple human pluripotent stem cell derived cardiomyocyte platforms. a**, Fraction of MYH7. **b**, Fraction of MYL2. **c**, Fraction of TNNI3. Gene counts expressed as a ratio RNA-sequencing data collected from hPSC-CM cultures including GSE93841[1], GSE148025[16], GSE116464[17], GSE201437[18], GSE114976[19] and human hearts including ERP109940[20] and Hahn et al., 2021[21].

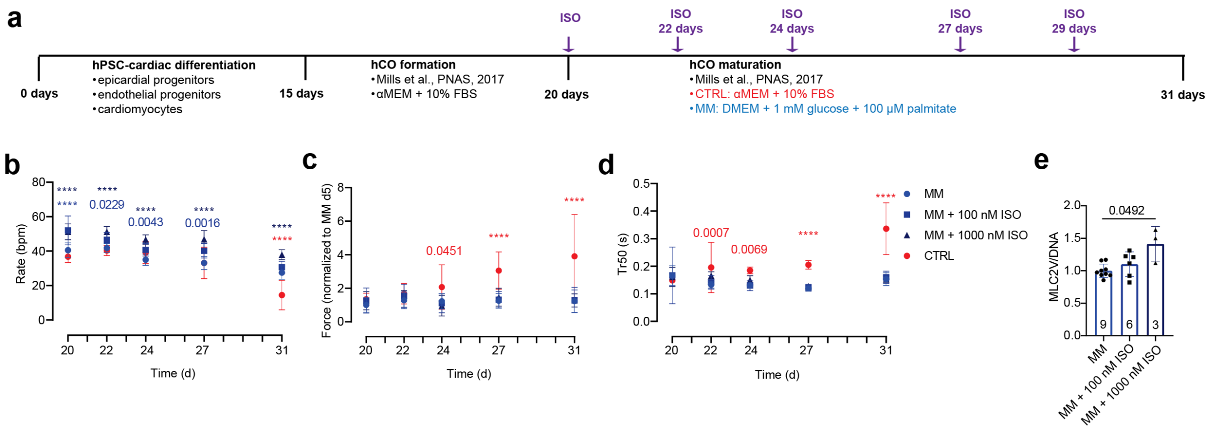


**Extended Data Fig. 2 Increased rate in hCOs by chronic dosing of ISO. a**, Schematic of the protocol. **b**, Rate. **c**, Force of contraction. **d**, Time from peak to 50% relaxation (Tr50). n = 14-16 (MM), 9-14 (MM + 100 nM ISO), 7-14 (MM + 1000 nM ISO), 8-10 (CTRL), 2 experiments (**b-d**). **e**, MLC2V intensity normalized to DNA. 1-2 experiments. Two-way ANOVA with Dunnett's multiple comparison to MM (**b-d**) and Krusksal-Wallis test with Dunn's multiple comparison to MM (**e**). **** P < 0.0001.

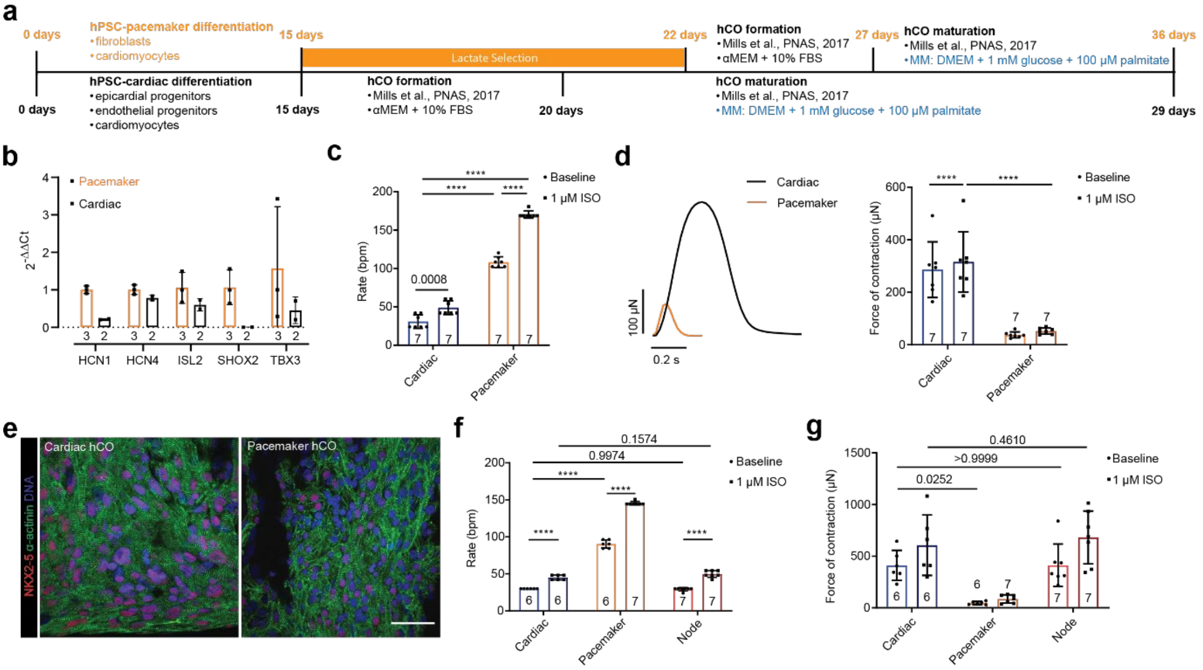


**Extended Data Fig. 3 Increased rate in hCOs using pacemaker cardiomyocytes. a**, Schematic of the protocol. **b**, Expression of key pacemaker genes. n = 2-3 experiments. **c**, Rate. **d**, Force of contraction trace and peak force. 1 experiment (**c-d**). **e**, Expression of NKX2-5 in cardiomyocytes (α-actinin). **f**, Rate. **g**, Force of contraction. 1 experiment (**f-g**). Two-way ANOVA with Sidaks's multiple comparison test (**c,d,f,g**). Bar = 20 μm. **** P < 0.0001.

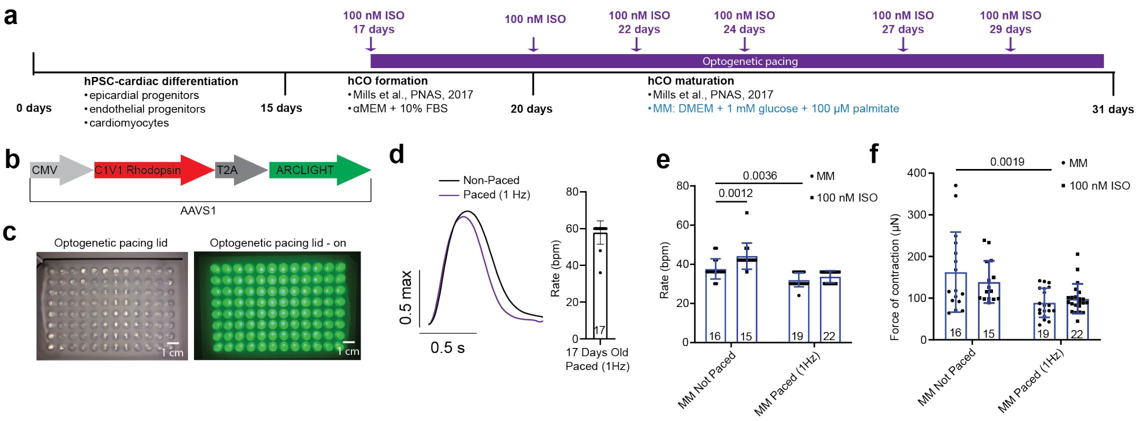


**Extended Data Fig. 4 Increased rate in hCOs using optogenetic pacing. a**, Schematic of the protocol. **b**, Construct inserted into AAVS1 (H9 hPSCs)**. c**, LED array lid used to stimulate pacing. **d**, Rate during pacing at the start of the protocol (17 days). 1 experiment. **e**, Rate without pacing at the end of the experiment (31 days). **f**, Force of contraction. 1 experiment (**e**,**f**). Two-way ANOVA with Sidak's multiple comparison test (**e,f**).

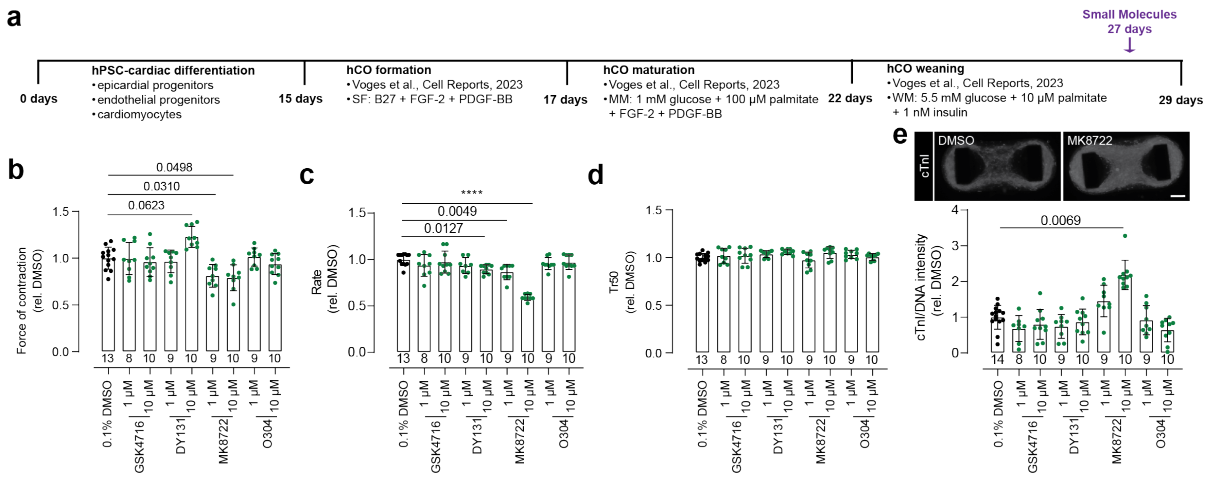


**Extended Data Fig. 5 Increased maturation with ERR and AMPK activators.** **a**, Schematic of the protocol. **b**, Force of contraction. **c**, Rate. **d**, Time from peak to 50% relaxation (Tr50). Functional data normalized to pre-treatment at 27 days. **e**, Cardiac troponin I (cTnI) intensity normalized to DNA. 2 experiments (**b-e**). Kruskal-Wallis test with Dunn's multiple comparison to DMSO (**b**-**e**). **** P < 0.0001. Bar = 200 µm.

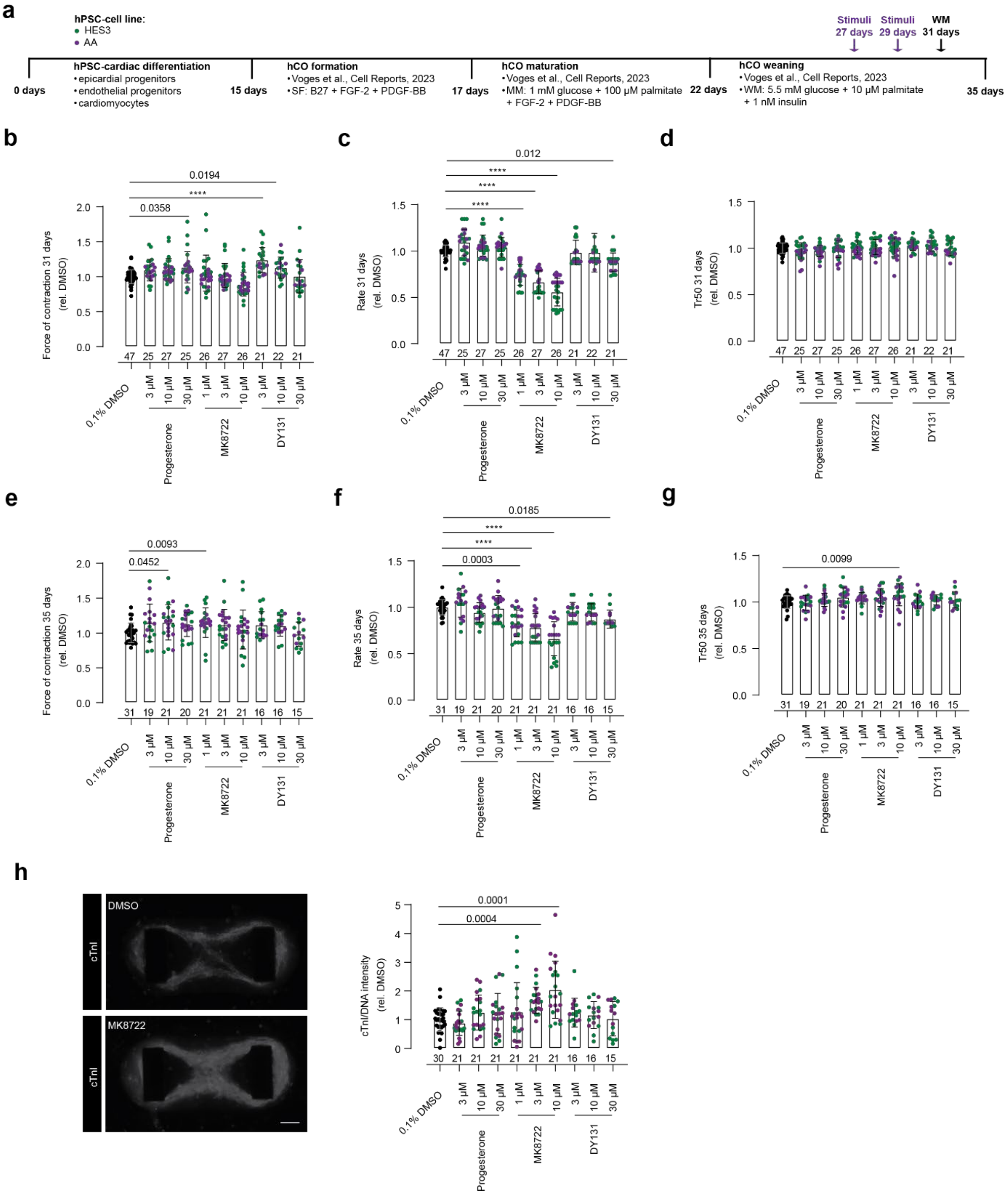

**Extended Data Fig. 6 Increased maturation with progesterone, DY131 and MK8722. a**, Schematic of the protocol. **b**, Force of contraction at 31 days. **c**, Rate at 31 days. **d**, Time from peak to 50% relaxation (Tr50) at 31 days. **e**, Force of contraction at 35 days. **f**, Rate at 35 days. **g**, Time from peak to 50% relaxation (Tr50) at 35 days. Functional data normalized to pre-treatment at 27 days. **h**, Cardiac troponin I (cTnI) intensity normalized to DNA. **i**, DNA intensity. 4 experiments, 2 HES3 (green) and 2 AA (purple), controls for all are in black (**b-i**). Kruskal-Wallis test with Dunn's multiple comparison to DMSO (**b**-**i**). **** P < 0.0001. Bar = 200 µm.

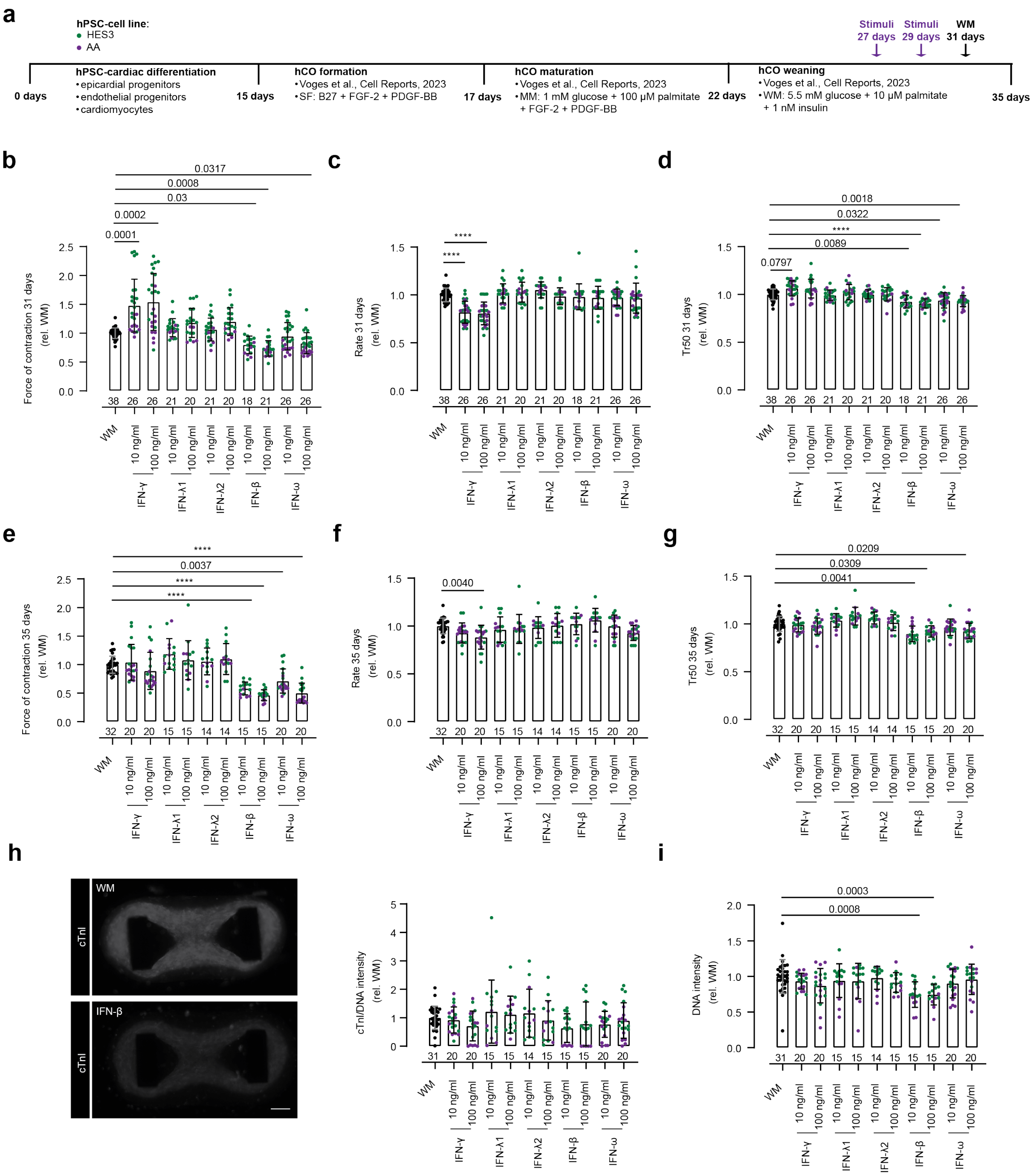


**Extended Data Fig. 7 Interferons have differential effects on hCO function. a**, Schematic of the protocol. **b**, Force of contraction at 31 days. **c**, Rate at 31 days. **d**, Time from peak to 50% relaxation (Tr50) at 31 days. **e**, Force of contraction at 35 days. **f**, Rate at 35 days. **g**, Time from peak to 50% relaxation (Tr50) at 35 days. Functional data normalized to pre-treatment at 27 days. **h**, Cardiac troponin I (cTnI) intensity normalized to DNA. **i**, DNA intensity. 4 experiments, 2 HES3 (green) and 2 AA (purple), controls for all are in black (**b-i**). Krusksal-Wallis test with Dunn's multiple comparison to DMSO (**b-i**). **** P < 0.0001. Bar = 200 μm.

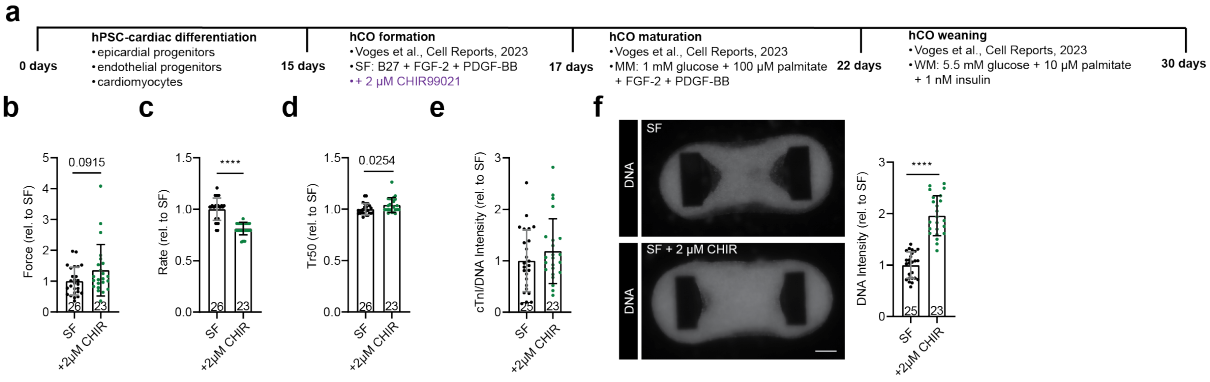


**Extended Data Fig. 8 Addition of CHIR99021 during hCO formation improves cell number. a**, Schematic of the protocol. **b**, Force of contraction at 30 days. **c**, Rate at 30 days. **d**, Time from peak to 50% relaxation (Tr50) at 30 days. Functional data normalized to the average of the SF condition for each experiment. **f**, DNA intensity. 2 experiments (**b-f**). Mann-Whitney test (**b-f**). **** P < 0.0001. Bar = 200 µm.

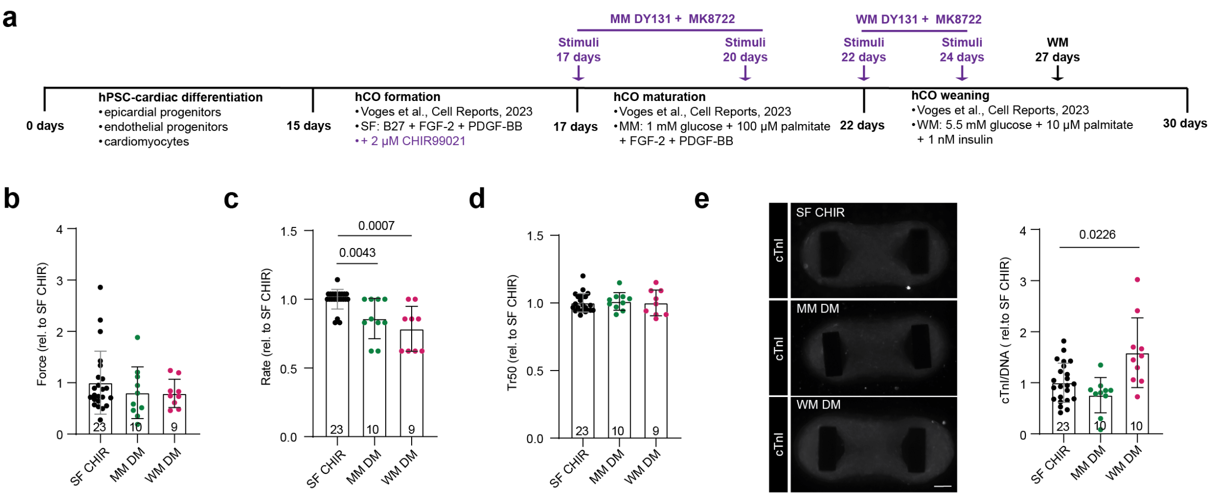


**Extended Data Fig. 9 Addition of DY131 and MK8722 are optimal during the WM phase. a**, Schematic of the protocol. **b**, Force of contraction at 30 days. **c**, Rate at 30 days. **d**, Time from peak to 50% relaxation (Tr50) at 30 days. **e,** Cardiac troponin I (cTnI) intensity normalized to DNA. (**b-e**). 2 experiments. Krusksal-Wallis test with Dunn's multiple comparison to SF + CHIR (**b-f**). Bar = 200 µm.

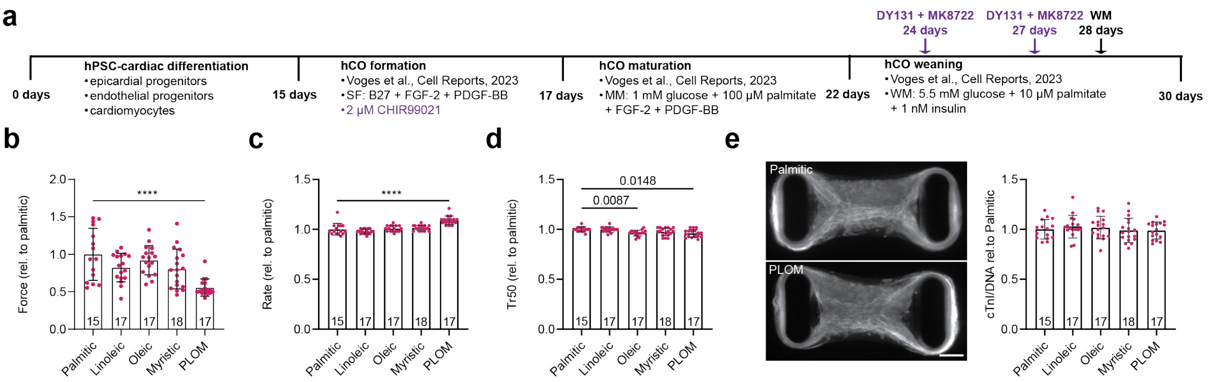


**Extended Data Fig. 10 Type of fatty acid has limited impact on DM-hCO function and cTnI expression.** **a**, Schematic of the protocol. **b**, Force of contraction at 30 days. **c**, Rate at 30 days. **d**, Time from peak to 50% relaxation (Tr50) at 30 days. **e,** Cardiac troponin I (cTnI) intensity normalized to DNA. (**b-e**). 2 experiments. Krusksal-Wallis test with Dunn's multiple comparison to palmitic acid (**b**-**e**). **** P < 0.0001. Bar = 200 μm. PLOM – 25 μM of each fatty acid.

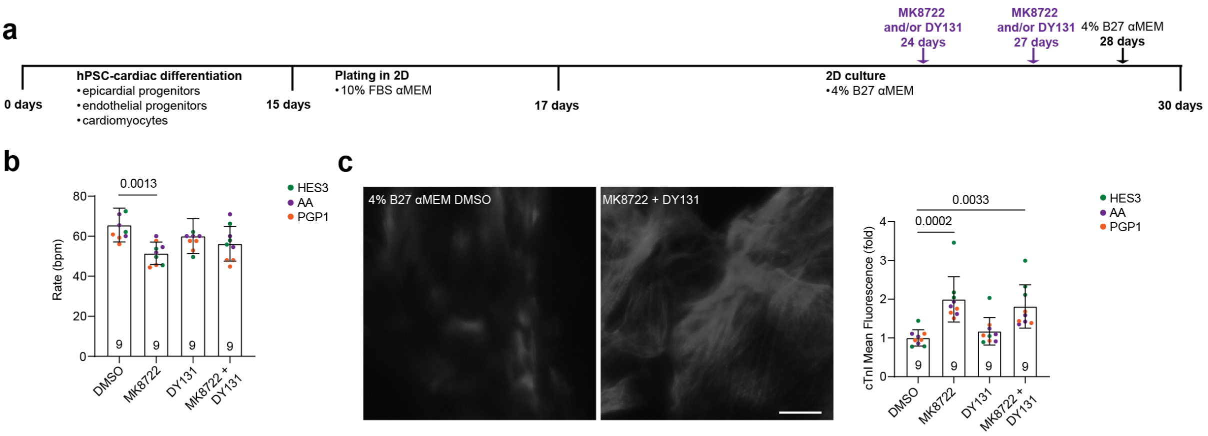


**Extended Data Fig. 11 Addition of DY131 and MK8722 improve maturation features in 2D. a**, Schematic of the protocol. **b**, Rate at 30 days. **c**, Normalized (to DMSO) mean fluorescence intensity (MFI) of cardiac troponin I (cTnI). 1 experiment per cell line. Krusksal-Wallis test with Dunn's multiple comparison to DMSO (**b,c**). Bar = 20 µm.

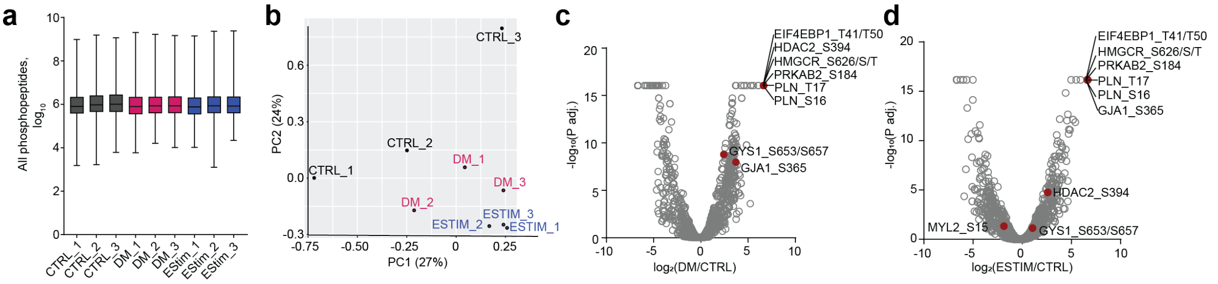


**Extended Data Fig. 12 Phosphoproteomics comparing between electrically paced and DM treated hCOs. a,** Intensity of phosphopeptides across the different samples. Each sample is 15 hCOs pooled. **b,** Samples plotted on their principal components PC1 and PC2. **c,** Volcano plot of DM treatment versus control hCOs. **d,** Volcano plot of ESTIM versus control hCOs. DM – 10 µM MK8722 + 3 µM DY131, ESTIM – 120 bpm paced hCOs, both for 5 minutes.

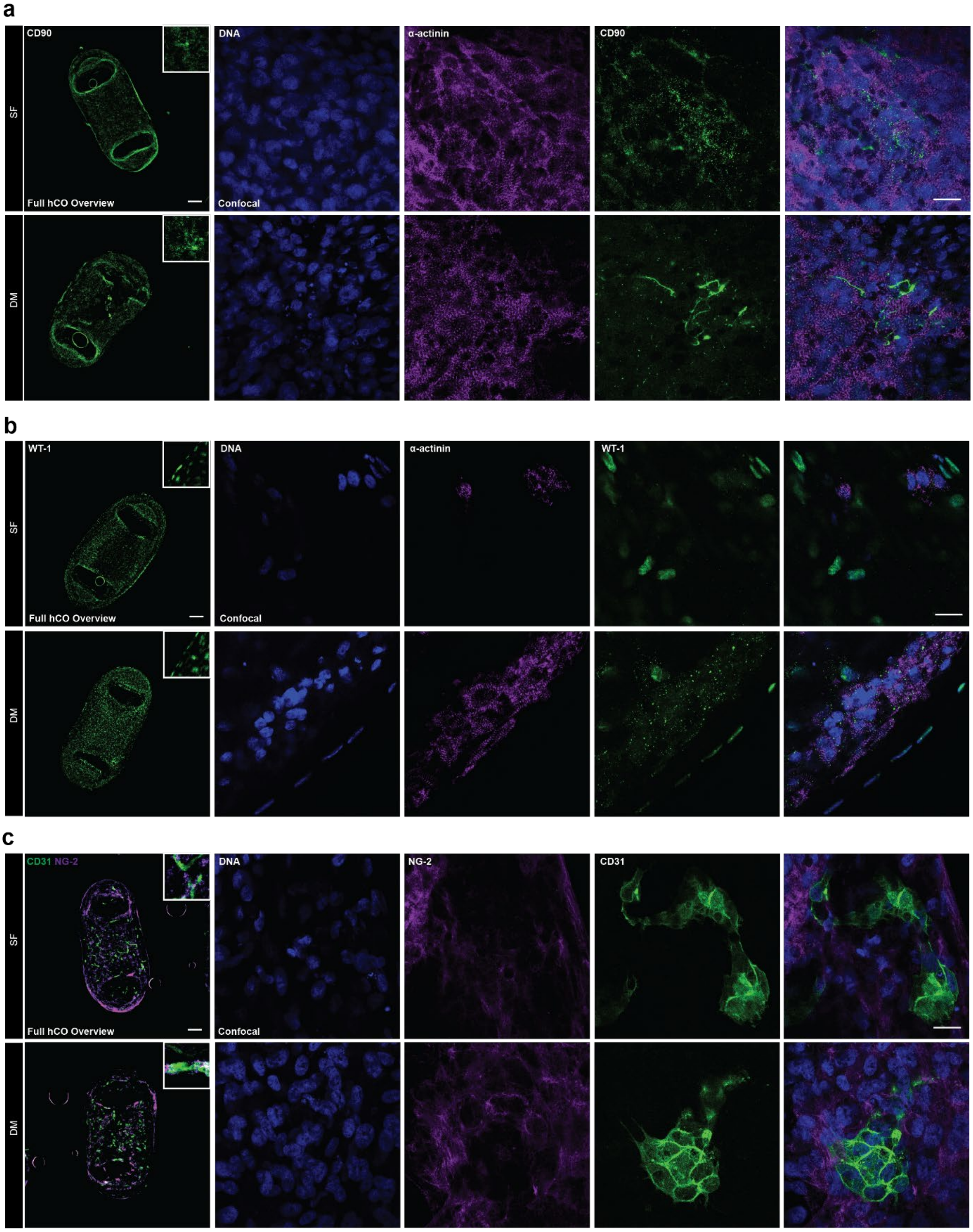


**Extended Data Fig. 13 Multiple cell populations are present in both SF- and DM-hCOs. a**, Fibroblast (CD90) and cardiomyocyte (α-actinin) staining. **b**, Epicardial cell (WT-1) and cardiomyocyte (α-actinin)

staining. **c**, Endothelial (CD31) and pericyte (NG-2) staining. Full hCO overview bar = 200 µm. Confocal bar = 20 µm.

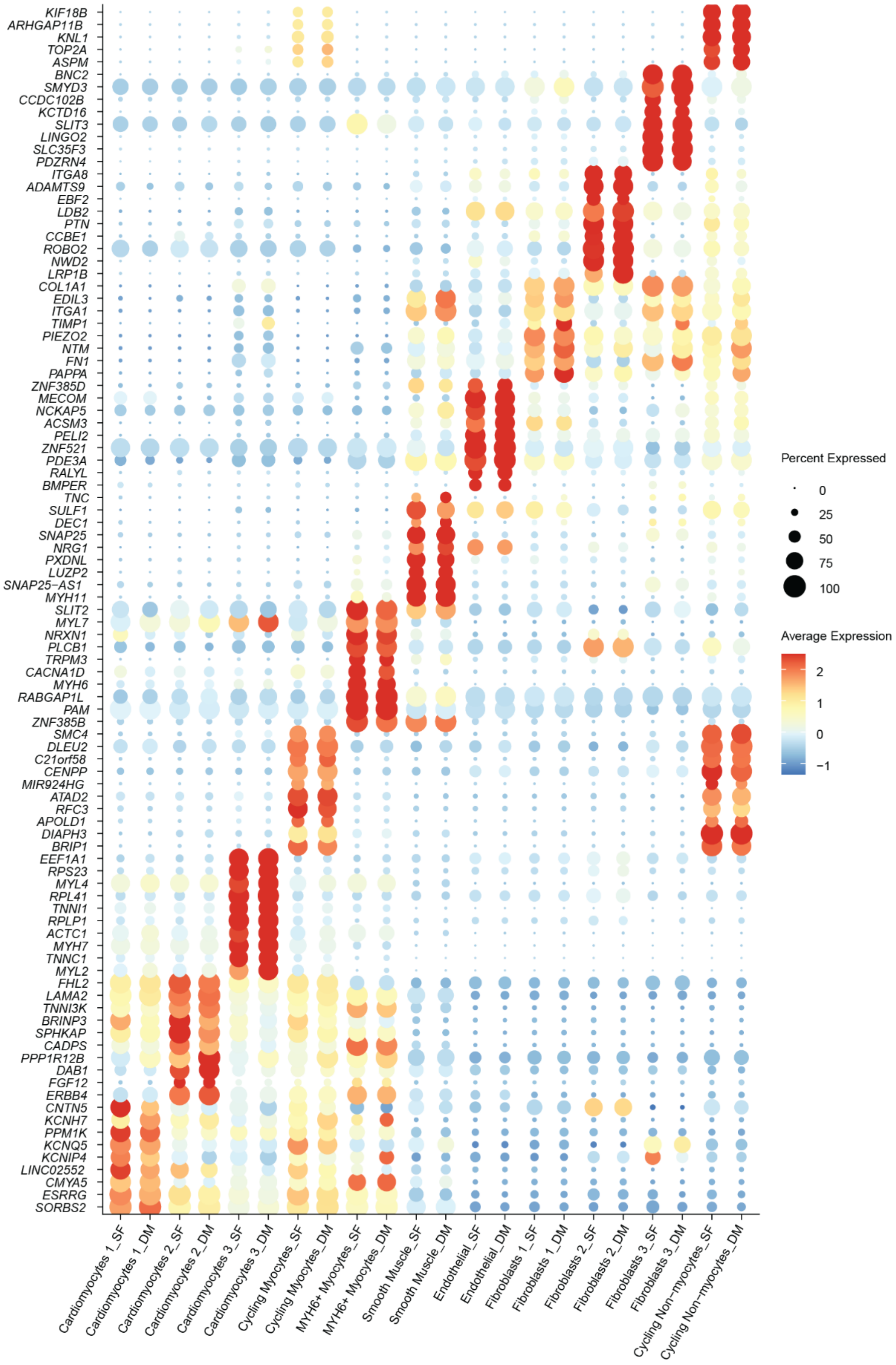


**Extended Data Fig. 14 Top 10 genes demarcating each cell population in SF- and DM-hCOs.**

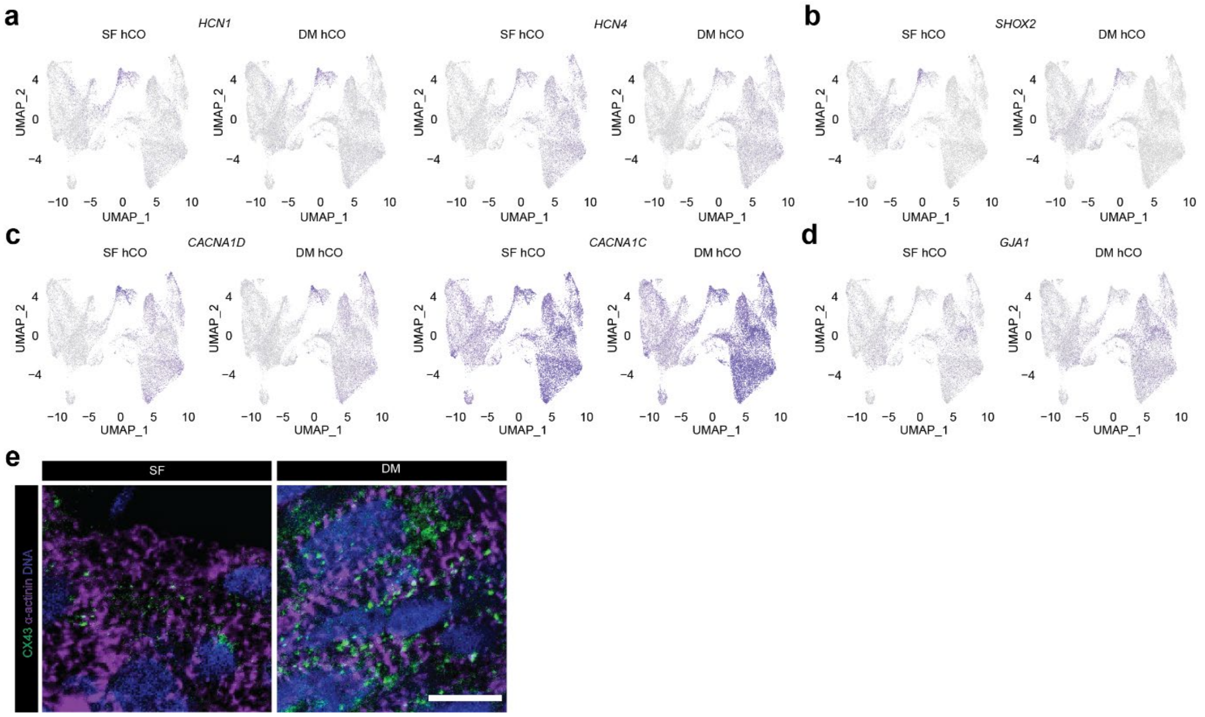


**Extended Data Fig. 15 Critical pace controlling genes in both SF- and DM-hCOs. a**, $I_f$ channel genes *HCN1* and *HCN4*. **b**, A key nodal cell transcription factor. **c**, Calcium channel genes for Cav1.2 (*CACNA1C*) and Cav1.3 (*CACNA1D*). **d**, Cell-cell junction gene for CX43 (*GJA1*). **e**, Immunostaining of CX43 in SF versus DM-hCOs. Scale bar = 10 µm.

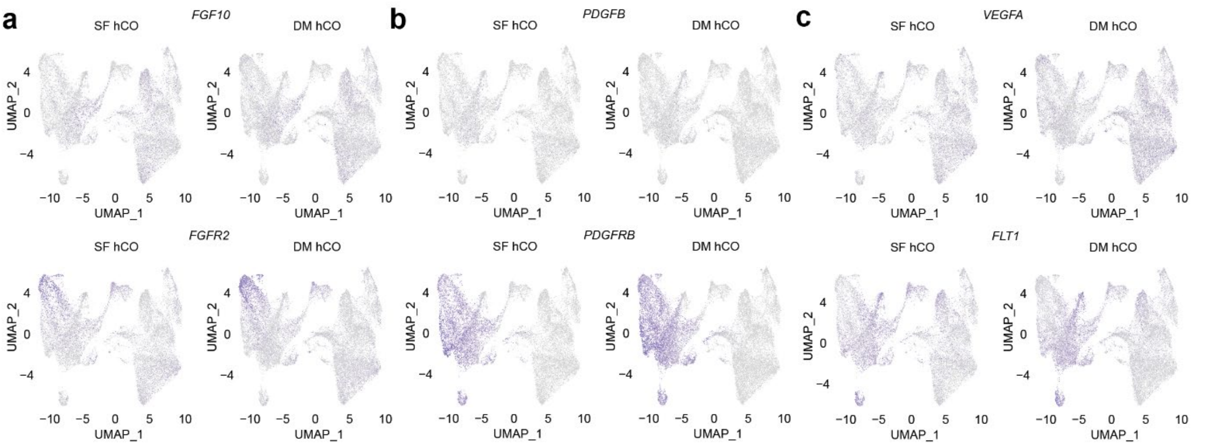


**Extended Data Fig. 16 Paracrine interactions in SF- and DM-hCOs.** **a**, *FGF10-FGFR2*. **b**, *PDGFB-PDGFRB*. **c**, *VEGFA-FLT1.*

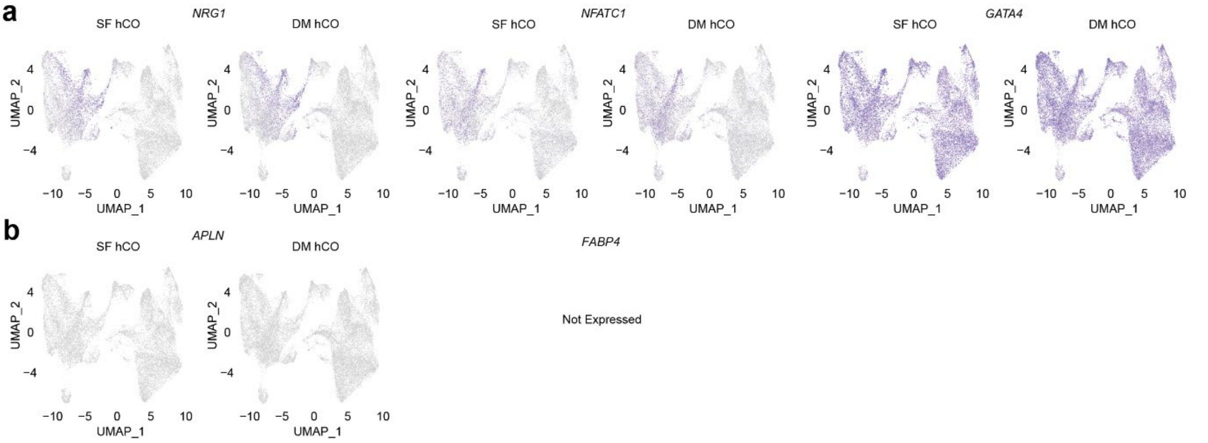


**Extended Data Fig. 17 Expression of endocardial versus coronary marker genes in SF- and DM-hCOs.** **a**, Endocardial markers. **b**, Coronary markers.

**a**

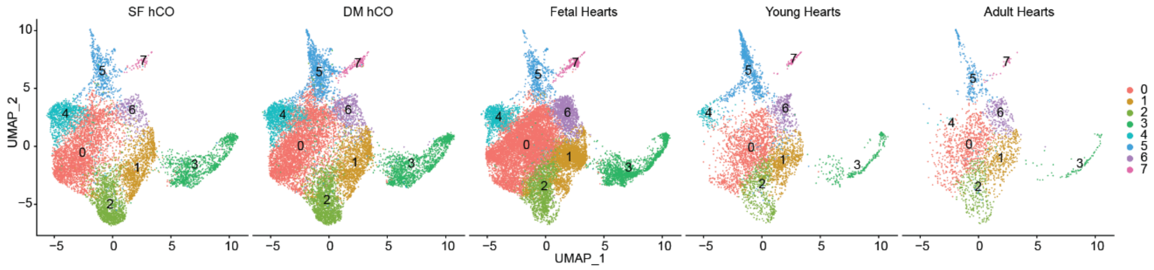


**Extended Data Fig. 18 Co-clustering of cardiomyocytes from SF and DM-hCOs and human heart cardiomyocytes form GSE156707[11].**

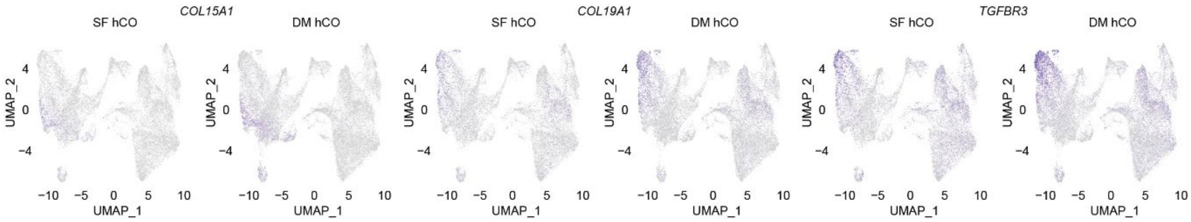


**Extended Data Fig. 19 Regulated genes in specific fibroblast populations in SF- and DM-hCOs.**

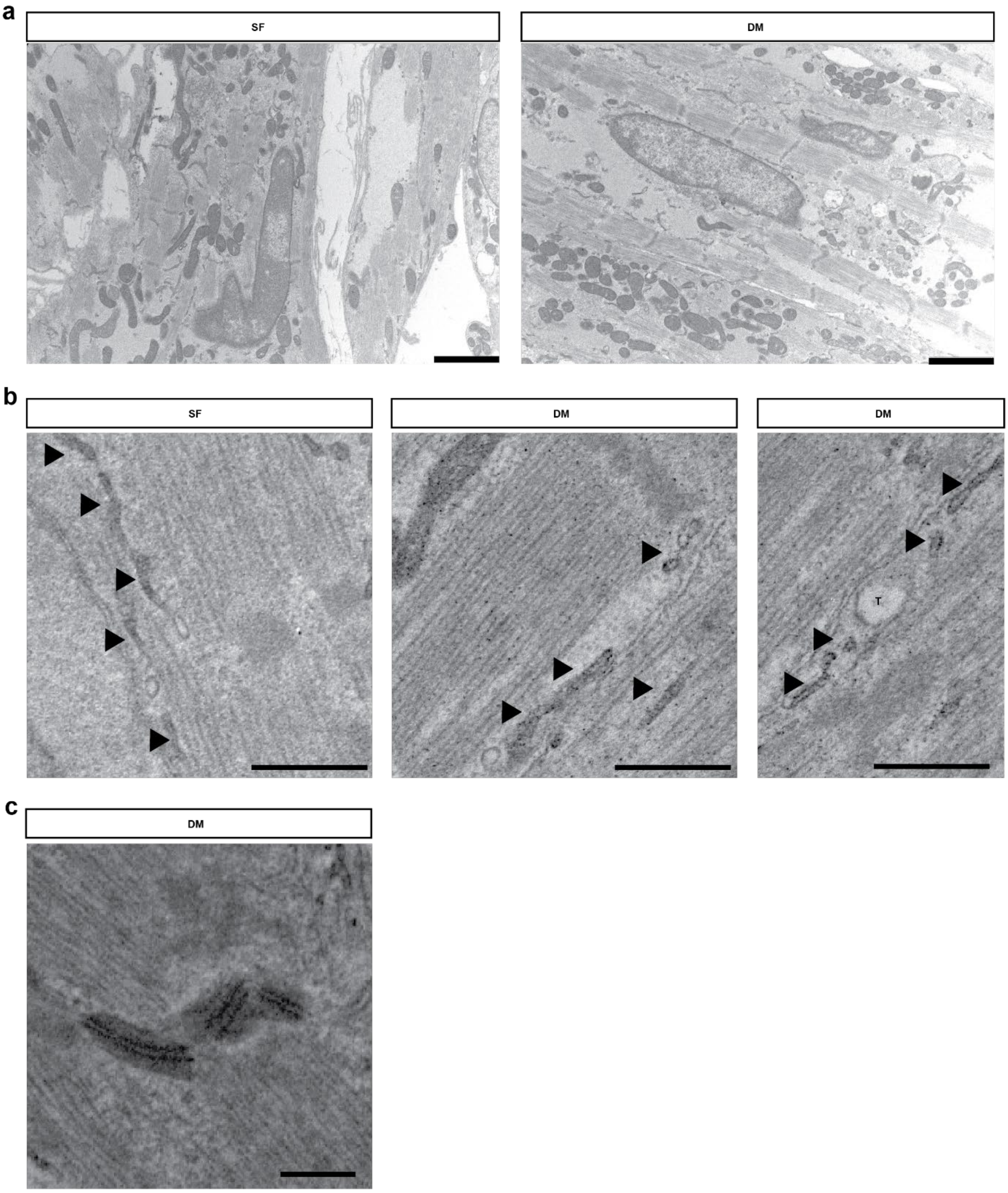


**Extended Data Fig. 20 SR and desmosomes are present in SF- and DM-hCOs. a**, Overview of a hCO section. Bar = 2 µm. **b**, Electron dense SR structures (indicated by arrows) and t-tubule (T). Bar = 500 nm. **c**, Electron dense intercalated discs. Bar = 500 nm.

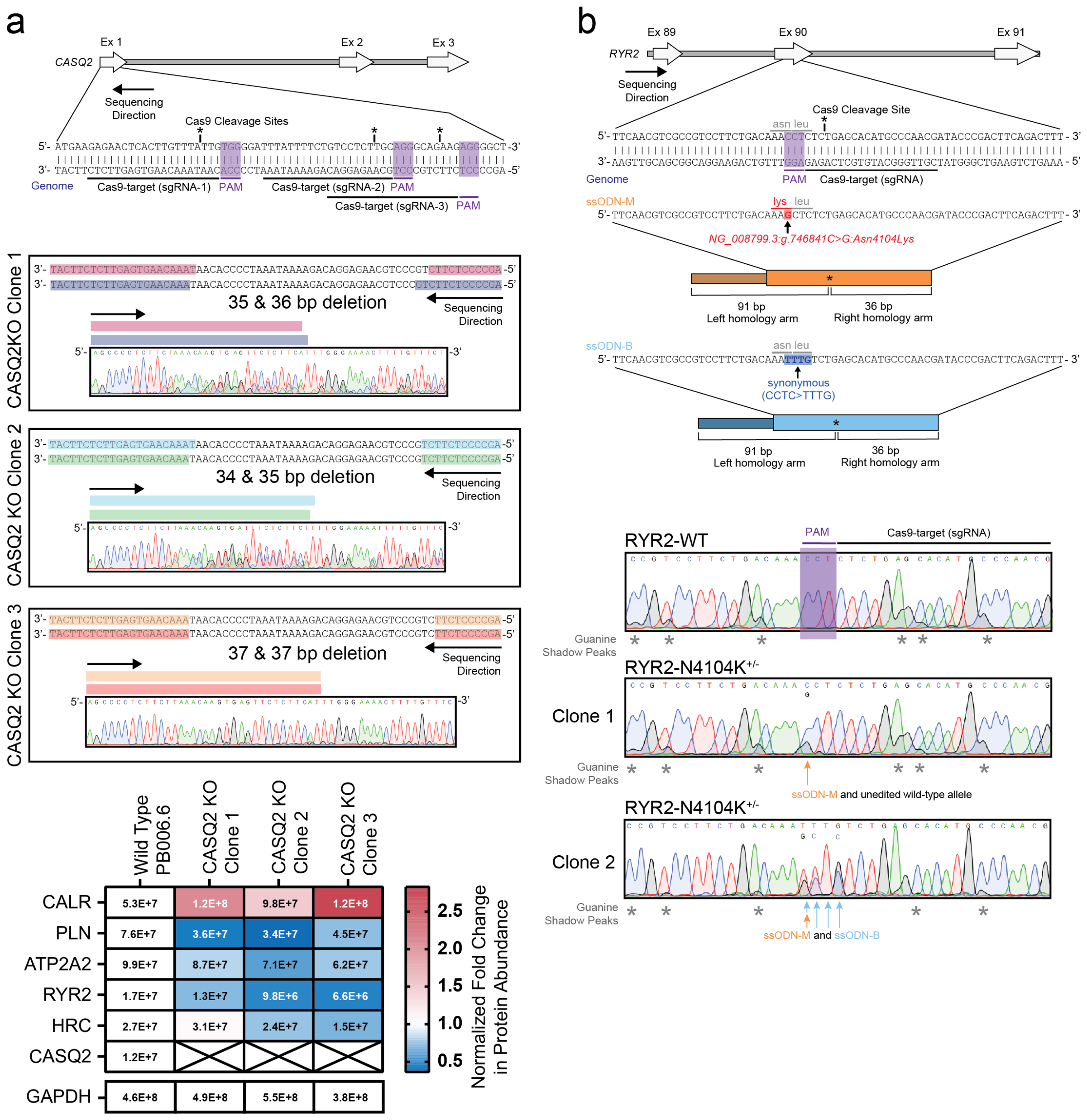


**CASQ2 amino acid sequence**

MKRTHLFIVGIYFLSSCRAEEGLNFPTYDGKDRVVSLSEKNFKQVLKK**YDLLCLYYHEPV**
**SSDK**VTQKQFQLKEIVLELVAQVLEHK**AIGFVMVDAK**KEAKLAKK**LGFDEEGSLYILK**GD
RTIEFDGEFAADVLVEFLLDLIEDPVEIISSK**LEVQAFER**IEDYIKLIGFFKSEDSEYYK
**AFEEAAEHFQPYIKFFATFDK**GVAKKLSLKMNEVDFYEPFMDEPIAIPNKPYTEEELVEF
VKEHQRPTLRRLRPEEMFETWEDDLNGIHIVAFAEK**SDPDGYEFLEILK**QVARDNTDNPD
LSILWIDPDDFPLLVAYWEKTFKIDLFRPQIGVVNVTDADSVWMEIPDDDDLPTAEELED
WIEDVLSGKINTEDDDEDDDDDDNSDEEDNDDSDDDDDE

**Peptide positions**

| | |
|---|---|
| YDLLCLYYHEPVSSDK | 48-86 |
| AIGFVMVDAK | 88-97 |
| LGFDEEGSLYILK | 106-118 |
| LEVQAFER | 152-160 |
| AFEEAAEHFQPYIK | 180-194 |
| FFATFDK | 195-201 |
| SDPDGYEFLEILK | 227-289 |

**Extended Data Fig. 21 Generation of CASQ2$^{-/-}$ and RYR2$^{N4104K/+}$cell lines using CRISPR. a,** Schematic of the CRISPR editing protocol and SANGER sequencing for CASQ2$^{-/-}$. This includes proteomics on DM-hCOs confirming knockdown of CASQ2 and other SR related proteins and identified CASQ2 peptides. **b,** Schematic of the CRISPR editing protocol and SANGER sequencing for RYR2$^{N4104K/+}$.

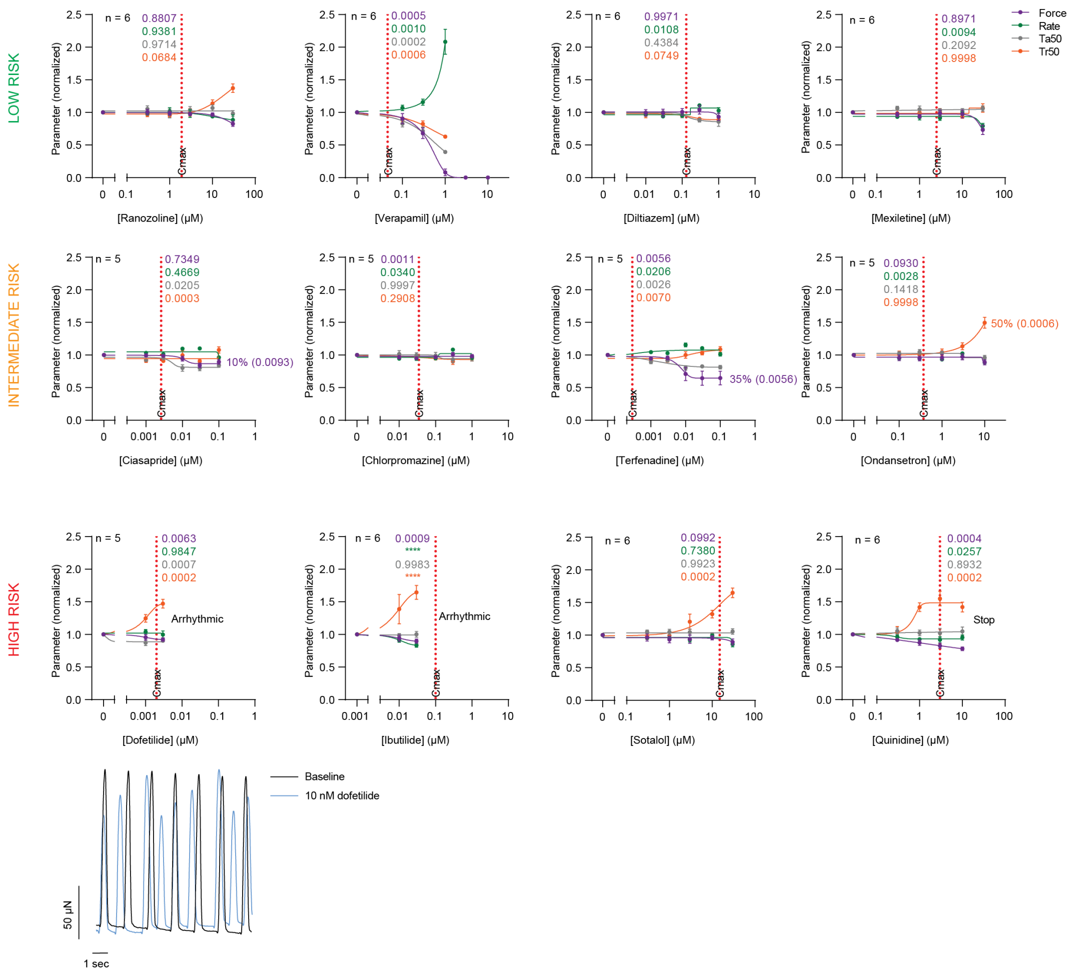


**Extended Data Fig. 22 DM-hCOs predict CiPA compound low and high risk.** Mixed –effects testing with Dunnett's post-hoc analysis in comparison to baseline. Only the statistics for concentration closest to Cmax are shown. Cmax values were derived from the literature. DM-hCOs did not alter their force, rate or Ta50 more than 10% in response to the CiPA compounds at the closest concentrations to Cmax, except for high quinidine (force), ibutilide (force and rate), verapamil (force and rate) and terfenadine (Ta50). Richard's 5 parameter dose-response was used to fit the curves.

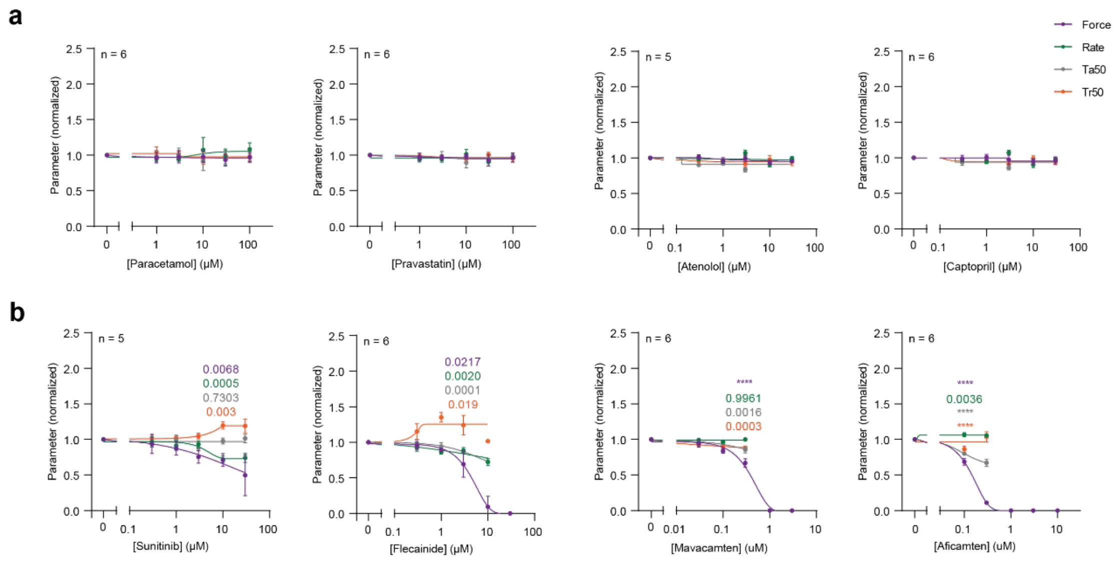


**Extended Data Fig. 23 DM-hCOs predict negative inotropes. a,** DM-hCO responses to benign drugs (paracetamol and pravastatin) or drugs inhibiting the impact of systemic factors (atenolol and captopril). **b**, DM-hCO responses to negative inotropes. Mixed –effects testing with Dunnett's post-hoc analysis in comparison to baseline. **** P < 0.0001. Only the statistics for lowest concentration with a decline in force are shown, as further force decline will impact the other parameters. n indicates number of DM-hCOs. Richard's 5 parameter dose-response was used to fit the curves.

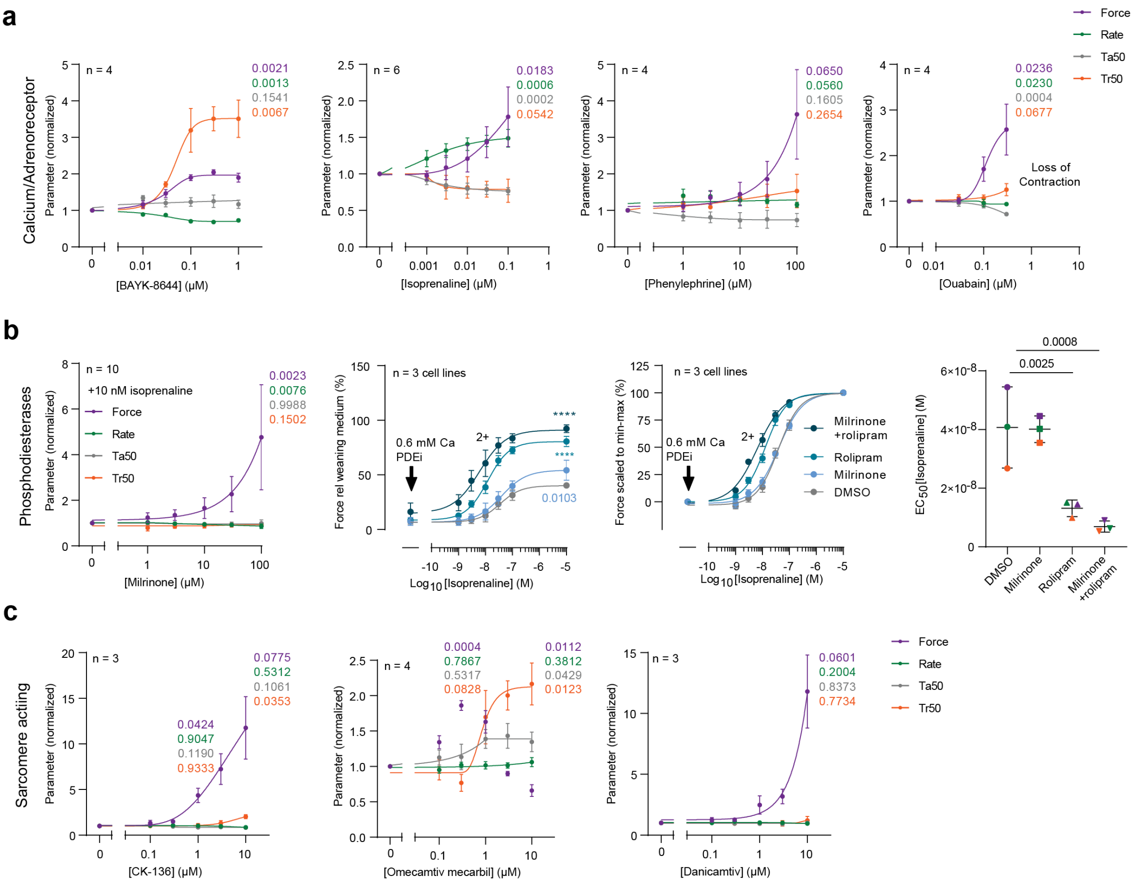


**Extended Data Fig. 24 DM-hCOs to predict positive inotropes. a,** DM-hCO responses to drugs modulating L-type calcium channel (BAYK-8644), β-adrenergic signalling (isoprenaline), $\alpha_1$-adrenergic signalling (phenylephrine) or Na+, K+ - ATPase (ouabain). **b**, DM-hCO responses to phosphodiesterase inhibition using milrinone (PDE3/PDE4) and rolipram (PDE4). For isoprenaline curves 10 µM and 30 µM milrinone were used. **c**, DM-hCO responses to sarcomeric acting inotropes targeting troponin (CK-136) or myosin (omecamtiv mecarbil and danicamtiv). All experiments were performed at 0.6 mM $Ca^{2+}$ by mixing weaning medium made in RPMI and DMEM base. Mixed –effects testing with Dunnett's post-hoc analysis in comparison to baseline (**a**-**c**), two-way ANOVA with Dunnett's post hoc analysis relative to DMSO (isoprenaline curves in **b**), one-way ANOVA with Dunnett's post hoc analysis relative to DMSO (EC50 in **b**). **** P < 0.0001. Statistics for highest concentration are shown, with the addition of intermediate concentrations where Tr50 is impacted. n indicated number of DM-hCOs unless stated otherwise. Richard's 5 parameter dose-response was used to fit the curves.

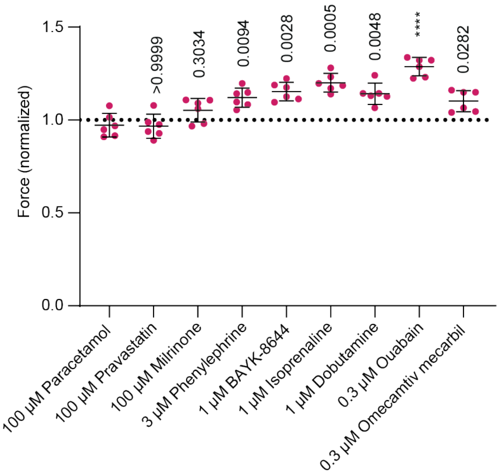


**Extended Data Fig. 25 DM-hCOs predict positive inotropes at 1.8 mM $Ca^{2+}$.** DM-hCO responses to inotropes modulating L-type calcium channel (BAYK-8644), β-adrenergic signalling (isoprenaline, dobutamine), $\alpha_1$-adrenergic signalling (phenylephrine) or Na+, K+ - ATPase (ouabain), milrinone (PDE3/PDE4) and myosin (omecamtiv mecarbil). Paracetamol and pravastatin were used as inert control drugs. n = 5-6 DM-hCOs. Brown-Forsythe and Welch's ANOVA test with Dunnett T3 post-hoc analysis relative to paracetamol. **** $P < 0.0001$.

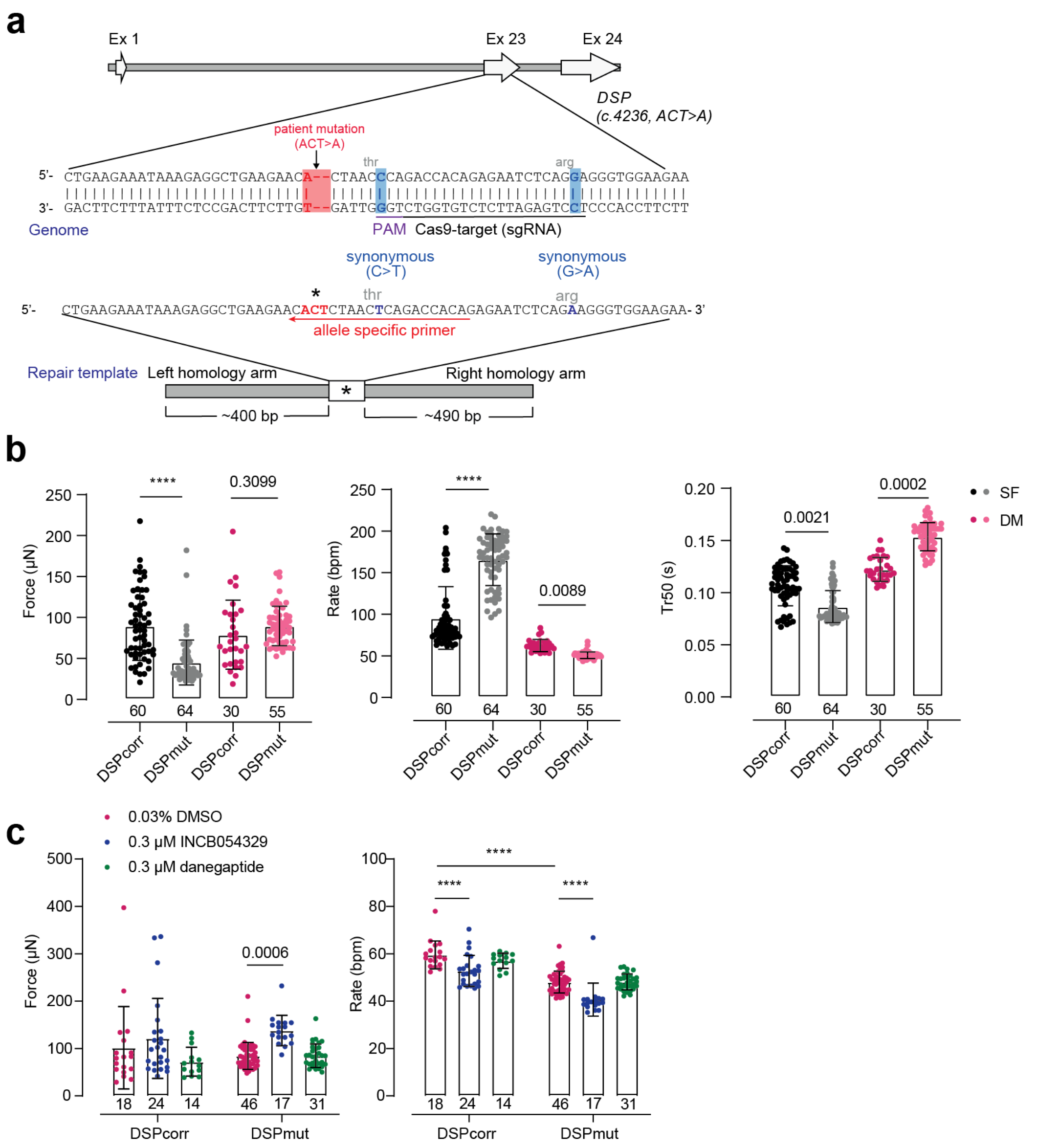


**Extended Data Fig. 26 DSPmut (MCHTB11) and DSPcorr (CRISPR corrected H.3) function in SF- and DM-hCOs. a,** Schematic of the CRISPR correction protocol**. b,** Contraction parameters of SF- and DM-hCOs. n = 3-4 experiments. **c,** Contraction parameters with treatment of INCB054329 and danegaptide in DM-hCOs (as per Fig. 6d). n = 3-4 experiments. Kruskal-Wallis test with Dunn's post-hoc analysis (**b**). Two-way ANOVA with Sidak's post-hoc test between lines for DMSO or relative to DMSO for DSPmut (**c**). **** $p < 0.0001$.

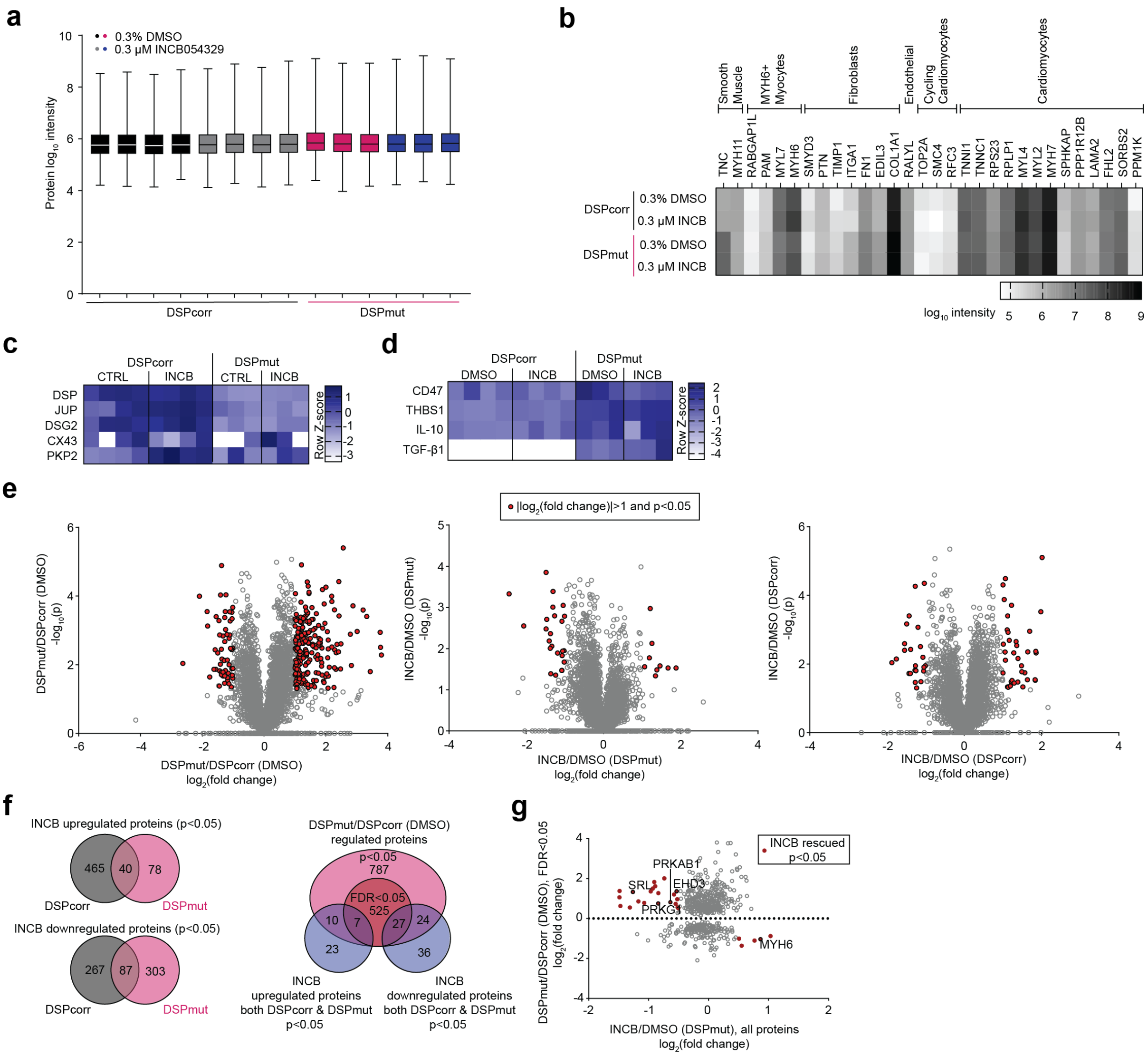

**Extended Data Fig. 27 Proteomic analysis of DSPmut and DSPcorr DM-hCOs treated with INCB0554329 (INCB). a,** Protein abundance. **b,** Protein abundance of cell markers**. c,** Heatmap of critical desmosomal proteins. **d**, Heatmap of critical immunomodulatory proteins. **e,** Volcano plots for different comparisons. **f,** Proteins regulated by INCB in DSPmut and DSPcorr DM-hCOs. **g,** Regulated proteins in DSPmut that are reverted by INCB. n = 3-4 experiments, each with 3 pooled DM-hCOs.

**Extended Data Table 1: Comparison of DM-hCO drug sensitivities with *in vivo* derived heart muscle or cardiomyocytes.**

| Compound | Inotrope | Target | IC50 or EC50 (µM) | Reference | Assay | DM hCO Half Maximal Effect (µM) | Maximum Effect Dose |
|---|---|---|---|---|---|---|---|
| Milrinone | Pos | PDE3/PDE4 | 62 (estimated) | Böhm et al., 1988[22] | Human left ventricle papillary muscle force | 46 | 100 µM |
| Phenylephrine | Pos | α1-AR | 1.5 (estimated) | Grimm et al., 2005[23] | Human ventricular trabecular muscle force | 48 | 100 µM |
| Paracetamol | None | Prostaglandin | | | | None | |
| Pravastatin | None | HMGCR | | | | None | |
| Ouabain | Pos | NKA | 0.45 | Brown and Erdmann, 1985[24] | Guinea pig papillary muscle force | 0.1 | 0.3 µM |
| Sunitinib | Neg | RTK | 7 | Harmer et. al., 2012[25] | Beagle dog isolated ventricular cardiomyocyte fractional shortening | 6.6 | |
| Flecainide | Neg | $I_{Na}$ | 7.8 | Harmer et. al., 2012[25] | Beagle dog isolated ventricular cardiomyocyte fractional shortening | 4.5 | |
| Verapamil | Neg | LTCC | 0.8 | Schwinger et al., 1990[26] | Human papillary muscle force | 0.4 | |
| Captopril | None | ACE | No basal stimulation in hCO | | | None | |
| Atenolol | None | β1-AR | No basal stimulation in hCO | | | None | |
| Aficamten | Neg | Myosin | 0.23 (free) | Chuang et al., 2021[27] | Beagle dogs fractional shortening in vivo | 0.14 | |
| Clonidine | None | α2-AR | No basal stimulation hCO | Saleem et al., 2020[28] | Human engineered heart tissues | None | |
| BAYK8644 | Pos | LTCC | 0.04 | Näbauer and Erdmann, 1988[29] | Human papillary muscle force | 0.03 | |
| Isoprenaline | Pos | β-AR | 0.02 | Schwinger et al., 1991[30] | Human papillary muscle force | 0.04 | |
| Mavacamten | Neg | Myosin | 0.2-0.3 | Green et al., 2016[31] | Rat isolated ventricular cardiomyocyte fractional shortening | 0.4 | |
| Nelutroctiv | Pos | Troponin | 2.7 (estimated) | Romero et al., 2024[32] | Rat left ventricle fractional shortening in vivo | 2.2 | 10 µM |
| Danicamtiv | Pos | Myosin | 6 | Voors et al., 2020[33] | Left ventricle Yucatan mini-pig sarcomere ATP hydrolysis | 5.9 | 10 µM |
| Omecamtiv Mecarbil | Pos | Myosin | 0.36 | Gollapudi et al., 2017[34] | Guinea pig muscle fibres force | 0.1 | 1 µM |

PDE – phosphodiesterase, AR – adreneoreceptor, HMGCR – HMG-CoA reductase, NKA – sodium potassium exchanger, RTK – receptor tyrosine kinase, LTCC – L-Type calcium channel, ACE – angiotensin converting enzyme.

**Extended Data References**


1 Mills, R. J. *et al.* Functional screening in human cardiac organoids reveals a metabolic mechanism for cardiomyocyte cell cycle arrest. *Proc Natl Acad Sci U S A* **114**, E8372-e8381, doi:10.1073/pnas.1707316114 (2017).

2 Protze, S. I. *et al.* Sinoatrial node cardiomyocytes derived from human pluripotent cells function as a biological pacemaker. *Nat Biotechnol* **35**, 56-68, doi:10.1038/nbt.3745 (2017).

3 Mills, R. J. *et al.* Development of a human skeletal micro muscle platform with pacing capabilities. *Biomaterials* **198**, 217-227, doi:10.1016/j.biomaterials.2018.11.030 (2019).

4 Needham, E. J. *et al.* Phosphoproteomics of Acute Cell Stressors Targeting Exercise Signaling Networks Reveal Drug Interactions Regulating Protein Secretion. *Cell Rep* **29**, 1524-1538.e1526, doi:10.1016/j.celrep.2019.10.001 (2019).

5 Voges, H. K. *et al.* Vascular cells improve functionality of human cardiac organoids. *Cell Rep* **42**, 112322, doi:10.1016/j.celrep.2023.112322 (2023).

6 Sakamoto, T. *et al.* A Critical Role for Estrogen-Related Receptor Signaling in Cardiac Maturation. *Circ Res* **126**, 1685-1702, doi:10.1161/circresaha.119.316100 (2020).

7 Arad, M., Seidman, C. E. & Seidman, J. G. AMP-activated protein kinase in the heart: role during health and disease. *Circ Res* **100**, 474-488, doi:10.1161/01.Res.0000258446.23525.37 (2007).

8 Zuercher, W. J. *et al.* Identification and structure-activity relationship of phenolic acyl hydrazones as selective agonists for the estrogen-related orphan nuclear receptors ERRbeta and ERRgamma. *J Med Chem* **48**, 3107-3109, doi:10.1021/jm050161j (2005).

9 Yu, D. D. & Forman, B. M. Identification of an agonist ligand for estrogen-related receptors ERRbeta/gamma. *Bioorg Med Chem Lett* **15**, 1311-1313, doi:10.1016/j.bmcl.2005.01.025 (2005).

10 Myers, R. W. *et al.* Systemic pan-AMPK activator MK-8722 improves glucose homeostasis but induces cardiac hypertrophy. *Science* **357**, 507-511, doi:10.1126/science.aah5582 (2017).

11 Sim, C. B. *et al.* Sex-Specific Control of Human Heart Maturation by the Progesterone Receptor. *Circulation* **143**, 1614-1628, doi:10.1161/circulationaha.120.051921 (2021).

12 Huss, J. M., Garbacz, W. G. & Xie, W. Constitutive activities of estrogen-related receptors: Transcriptional regulation of metabolism by the ERR pathways in health and disease. *Biochim Biophys Acta* **1852**, 1912-1927, doi:10.1016/j.bbadis.2015.06.016 (2015).

13 Mills, R. J. *et al.* BET inhibition blocks inflammation-induced cardiac dysfunction and SARS-CoV-2 infection. *Cell* **184**, 2167-2182.e2122, doi:10.1016/j.cell.2021.03.026 (2021).

14 Buikema, J. W. *et al.* Wnt Activation and Reduced Cell-Cell Contact Synergistically Induce Massive Expansion of Functional Human iPSC-Derived Cardiomyocytes. *Cell Stem Cell* **27**, 50-63.e55, doi:10.1016/j.stem.2020.06.001 (2020).

15 Quaife-Ryan, G. A. *et al.* β-Catenin drives distinct transcriptional networks in proliferative and nonproliferative cardiomyocytes. *Development* **147**, doi:10.1242/dev.193417 (2020).

16 Hofbauer, P. *et al.* Cardioids reveal self-organizing principles of human cardiogenesis. *Cell* **184**, 3299-3317.e3222, doi:10.1016/j.cell.2021.04.034 (2021).

17 Giacomelli, E. *et al.* Human-iPSC-Derived Cardiac Stromal Cells Enhance Maturation in 3D Cardiac Microtissues and Reveal Non-cardiomyocyte Contributions to Heart Disease. *Cell Stem Cell* **26**, 862-879.e811, doi:10.1016/j.stem.2020.05.004 (2020).

18 Shen, S. *et al.* Physiological calcium combined with electrical pacing accelerates maturation of human engineered heart tissue. *Stem Cell Reports* **17**, 2037-2049, doi:10.1016/j.stemcr.2022.07.006 (2022).

19 Zhao, Y. *et al.* A Platform for Generation of Chamber-Specific Cardiac Tissues and Disease Modeling. *Cell* **176**, 913-927.e918, doi:10.1016/j.cell.2018.11.042 (2019).

20 Pervolaraki, E., Dachtler, J., Anderson, R. A. & Holden, A. V. The developmental transcriptome of the human heart. *Sci Rep* **8**, 15362, doi:10.1038/s41598-018-33837-6 (2018).

21 Hahn, V. S. *et al.* Myocardial Gene Expression Signatures in Human Heart Failure With Preserved Ejection Fraction. *Circulation* **143**, 120-134, doi:10.1161/circulationaha.120.050498 (2021).

22 Böhm, M., Diet, F., Kemkes, B. & Erdmann, E. Enhancement of the effectiveness of milrinone to increase force of contraction by stimulation of cardiac beta-adrenoceptors in the failing human heart. *Klin Wochenschr* **66**, 957-962, doi:10.1007/bf01738110 (1988).

23 Grimm, M. *et al.* Key role of myosin light chain (MLC) kinase-mediated MLC2a phosphorylation in the alpha 1-adrenergic positive inotropic effect in human atrium. *Cardiovasc Res* **65**, 211-220, doi:10.1016/j.cardiores.2004.09.019 (2005).

24 Brown, L. & Erdmann, E. Concentration-response curves of positive inotropic agents before and after ouabain pretreatment. *Cardiovasc Res* **19**, 288-298, doi:10.1093/cvr/19.5.288 (1985).

25 Harmer, A. R. *et al.* Validation of an in vitro contractility assay using canine ventricular myocytes. *Toxicol Appl Pharmacol* **260**, 162-172, doi:10.1016/j.taap.2012.02.007 (2012).

26 Schwinger, R. H., Böhm, M. & Erdmann, E. Negative inotropic properties of isradipine, nifedipine, diltiazem, and verapamil in diseased human myocardial tissue. *J Cardiovasc Pharmacol* **15**, 892-899, doi:10.1097/00005344-199006000-00006 (1990).

27 Chuang, C. *et al.* Discovery of Aficamten (CK-274), a Next-Generation Cardiac Myosin Inhibitor for the Treatment of Hypertrophic Cardiomyopathy. *J Med Chem* **64**, 14142-14152, doi:10.1021/acs.jmedchem.1c01290 (2021).

28 Saleem, U. *et al.* Blinded, Multicenter Evaluation of Drug-induced Changes in Contractility Using Human-induced Pluripotent Stem Cell-derived Cardiomyocytes. *Toxicol Sci* **176**, 103-123, doi:10.1093/toxsci/kfaa058 (2020).

29 Näbauer, M., Brown, L. & Erdmann, E. Positive inotropic effects of the calcium channel activator Bay K 8644 on guinea-pig and human isolated myocardium. *Naunyn Schmiedebergs Arch Pharmacol* **337**, 85-92, doi:10.1007/bf00169482 (1988).

30 Schwinger, R. H., Böhm, M., Pieske, B. & Erdmann, E. Different beta-adrenoceptor-effector coupling in human ventricular and atrial myocardium. *Eur J Clin Invest* **21**, 443-451, doi:10.1111/j.1365-2362.1991.tb01393.x (1991).

31 Green, E. M. *et al.* A small-molecule inhibitor of sarcomere contractility suppresses hypertrophic cardiomyopathy in mice. *Science* **351**, 617-621, doi:10.1126/science.aad3456 (2016).

32 Romero, A. *et al.* Discovery of Nelutroctiv (CK-136), a Selective Cardiac Troponin Activator for the Treatment of Cardiovascular Diseases Associated with Reduced Cardiac Contractility. *J Med Chem* **67**, 7825-7835, doi:10.1021/acs.jmedchem.3c02413 (2024).

33 Voors, A. A. *et al.* Effects of danicamtiv, a novel cardiac myosin activator, in heart failure with reduced ejection fraction: experimental data and clinical results from a phase 2a trial. *Eur J Heart Fail* **22**, 1649-1658, doi:10.1002/ejhf.1933 (2020).

34 Gollapudi, S. K., Reda, S. M. & Chandra, M. Omecamtiv Mecarbil Abolishes Length-Mediated Increase in Guinea Pig Cardiac Myofiber Ca(2+) Sensitivity. *Biophys J* **113**, 880-888, doi:10.1016/j.bpj.2017.07.002 (2017).
